# Differential turnover of apicobasal regulators drives emergent mechano-response and shape homeostasis

**DOI:** 10.64898/2026.08.12.744353

**Authors:** Jesús M. López-Gay, Uday Ram Gubbala, Priscillia Pierre-Elies, Ines Cristo, Diana Pinheiro, Edouard Hannezo, Yohanns Bellaiche

## Abstract

Epithelial cell shape plays a fundamental role in tissue dynamics. Numerous studies have established how cells drastically change their shape to promote epithelial tissue morphogenesis. However, the mechanisms enabling cells to maintain their shape remain far less understood. Here, leveraging live imaging in *Drosophila* epithelial tissue and theoretical modeling, we identify an emergent mechano-chemical feedback that ensures junction length and cell shape stability, without requiring a dedicated molecular force sensor. We find that an increase in junction length is associated with a passive dilution of E-Cadherin, followed by an increase in Myosin-II-dependent contractility that reduces junction length. Theoretically, we show that this regulation of junction length generically emerges when negative and positive regulators of contractility have distinct kinetics. Experiments confirm that E-Cadherin acts as a negative regulator with slow turnover. Mechanistically, local dilution of E-Cadherin passively lifts an inhibition on lateral apicobasal polarity components, allowing the RhoGEF Cyst — with its fast turnover — to accumulate and increase contractility. Perturbing this feedback results in aberrant cell junction and shape regulation, thereby compromising the ability of the tissue to buffer local mechanical fluctuations and global mechanical stresses. Altogether, we propose that differential turnover between apical and lateral polarity complexes provides an emergent mechano-response for junction length and cell shape homeostasis.

## Introduction

How cells and organisms regulate their size and shape is one of the key outstanding questions in biology. In recent years, our understanding of the sensing and feedback mechanisms that ensure size control has improved substantially ^1–3^. At the single-cell scale, numerous studies have delineated how cell volume is sensed to promote cell cycle progression and division, thereby controlling cell size within tissues ^4–9^. Yet each cell within a tissue must regulate not only its volume but also its shape to control tissue morphogenesis. For instance, many epithelial morphogenetic movements, such as tissue elongation or invagination, are rooted in the ability of cells to precisely control local contractions and elongations of their apical cell-cell junctions ^10–14^. The actomyosin cytoskeleton is a key force-generating machinery driving these junctional length changes. These forces are transmitted at the supra-cellular level via cell-substrate and cell-cell junctions, specifically integrins and E-Cadherins (E-Cad) based adherens junctions (AJ), leading to tissue-scale deformations ^15–18^. However, it remains poorly understood, from both theoretical and experimental perspectives, how a given cell shape is maintained at short and long timescales.

Theoretical models of epithelial tissues (e.g., Vertex or Potts models), based on junctional actomyosin contractility and cell-cell adhesion, have shown strong predictive power: they explain a range of morphogenetic movements, predict the steady-state geometry and topology of confluent epithelial monolayers, and can even infer whether a living tissue behaves as a fluid or a solid ^19–23^. Interestingly, one of their key underlying assumptions is that each individual cell is able to precisely regulate its apical perimeter to a defined value ^24,25^, as small changes in this preferred parameter drive tissue-level rigidity transitions ^26–29^. Whether and how this “homeostatic” perimeter is biologically maintained over long timescales remains unclear, both mechanistically and conceptually. Since cell shape is regulated by mechanical forces ^30^, a homeostatic mechanism should be able to sense and respond to a fluctuating mechanical environment ^11,31–37^. Over the last decades, extensive work has been devoted to the study of cell and tissue mechano-response ^17,18,26,38,39^. This has been primarily characterized as mechanisms involving force-induced unfolding of specific junctional and cytoskeletal proteins, which then trigger downstream responses ^40–51^. Yet how to link these junctional mechanical responses, which occur at molecular length scales and on timescales of seconds, to the maintenance of mesoscopic junctional length and cell shape over developmental timescales (tens of minutes to hours) remains an outstanding challenge in the field.

Here, by combining a theoretical mechano-chemical model with live imaging and genetic perturbations in *Drosophila* epithelial tissue, we show that E-Cad AJs exhibit a collective, self-organized mechanical response that ensures junction length and cell shape maintenance, without requiring a dedicated molecular force sensor. Theoretically, we show that this control of junction length generically emerges when negative and positive regulators of contractility have distinct kinetics. Experimentally, E-Cad acts as a negative regulator with slower turnover, and its local dilution due to junction elongation passively lifts an inhibition on positive regulators of contractility with fast turnover. Mechanistically, the core apicobasal polarity machinery, typically associated with maintaining the apicobasal positioning of junctional complexes ^52^, is critical for such regulation of in-plane junction length. This reveals an uncharacterized role for apicobasal polarity machinery in mechano-chemical feedback. Finally, we find that the proposed mechano-chemical model provides a generic force-response mechanism that ensures junction length regulation under both low and high mechanical stress.

## Results

### Local E-Cad and Myo-II dynamics correlate with rapid changes in junction length

*Drosophila* epithelial tissues have provided fundamental insights into the conserved mechanisms controlling junction dynamics during development ^16,17^. To experimentally explore how junction length is regulated, we performed high-resolution time-lapse imaging of junction dynamics using both E-Cad:GFP and the Myosin II light-chain tagged with mKate2, Myo-II: mKate2x3, in the *Drosophila* pupal dorsal thorax (Extended Data Fig. 1A). The time-lapses were conducted over 40 minutes (min) with a time interval of 20 seconds (s) in the anterior region of the tissue between 18 and 20 hours after pupa formation (h APF, Supplementary Video 1). We then plotted each junction’s dynamics as a kymograph. Visual inspection of the time-lapse movie or the kymograph revealed a stereotypical dynamic of E-Cad, Myo-II, and junction length (Fig. 1A,A’). We observed instances of a local decrease in the junctional E-Cad and cortical Myo-II signals concomitant with junction lengthening, shortly followed by an increase in Myo-II, a recovery of E-Cad levels, and junction shortening (Fig. 1A,A’). The membrane PH:GFP marker remained along the junction at the sites of local E-Cad decreases, indicating that changes in E-Cad signal are not caused by membrane rupture (Extended Data Fig. 1B,C); such local E-Cad signal decrease will hereafter be referred to as an E-Cad signal gap.

**Figure 1.**
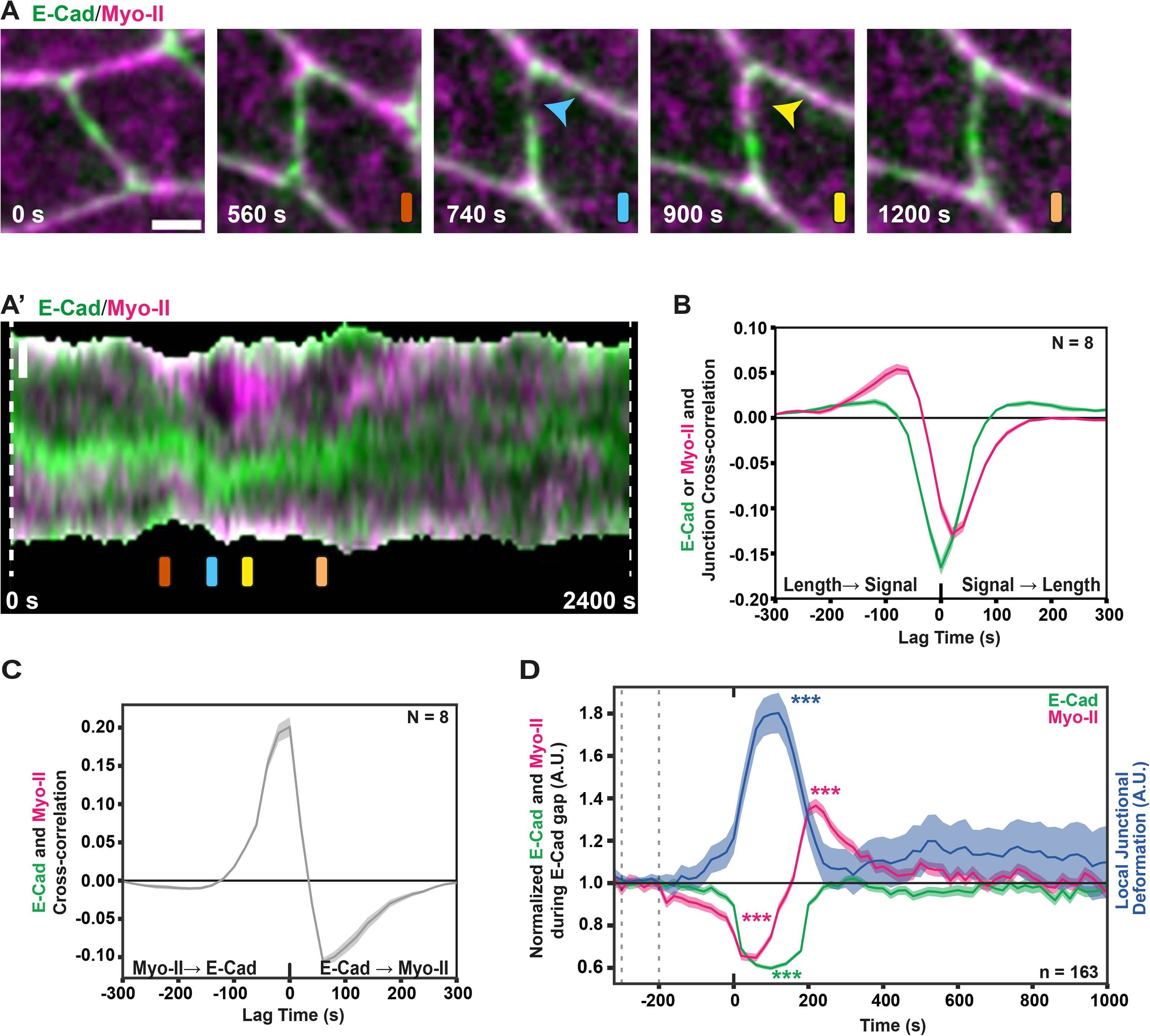
E-Cadherin and Myosin-II dynamics at epithelial junctions in *Drosophila*. **(A, A’)** Time-lapse images of E-Cad:GFP and Myo-II:mKate2x3 junction (A). Colored bars (orange, blue, yellow, peach) within panels indicate the corresponding time points in the kymograph in (A’). Blue and yellow arrowheads mark the position of E-Cad gap and the subsequent Myo-II accumulation. Kymograph of E-Cad:GFP and Myo-II:mKate2x3 along the junction shown in (A) over 2400 s (40 min, A’). Colored bars indicate the time points displayed in (A). **(B)** Temporal cross-correlation between junction length and E-Cad intensity or Myo-II intensity (mean ± SEM). Positive time lags indicate that the E-Cad or Myo-II signal precedes changes in junction length. Minimum number of junctions analyzed per animal was 161 junctions. **(C)** Temporal cross-correlation between E-Cad:GFP and Myo-II:mKate2x3 intensities along the junction (mean ± SEM). Positive time lags indicate that the E-Cad signal change precedes a change in Myo-II signal. Minimum number of junctions analyzed per animal was 161 junctions. **(D)** Normalized E-Cad:GFP, Myo-II:mKate2x3, and local junction deformation (median ± SEM) aligned to local E-Cad gap events (t = 0 s). E-Cad and Myo-II intensities, as well as local junction deformation, were normalized to their median baseline values between -300 and -200 s, indicated by dotted lines. Statistical significance was assessed using paired Wilcoxon signed-rank tests against baseline. n: total E-Cad gaps. Minimum number of animals analyzed for this quantification was 8. Scale bars: 2 µm (A, A’). Time is shown in seconds (s). Unless otherwise indicated, n and N indicate the number of junctions and animals, respectively. Asterisks indicate statistical significance: *p < 0.05, **p < 0.01, ***p < 0.001; ns, not significant.

To characterize the dynamics of E-Cad and Myo-II more quantitatively, we performed two sets of analyses. First, we used cross-correlation for different time delays between the E-Cad and Myo-II signals across the entire kymographs, while also keeping track of junctional length (see Methods). We found a robust anticorrelation between junction length and Myo-II/E-Cad at short timescales (0-20 s, Fig. 1B), as well as, with a delay of 60 s, an anticorrelation between E-Cad and Myo-II signals (Fig. 1C). This is consistent with the qualitative observation that both Myo-II and E-Cad decreased upon junction lengthening, and that the E-Cad decrease was followed by a subsequent Myo-II increase. Second, we used the kymographs to average the dynamics of E-Cad, Myo-II, and junction deformation, specifically in the regions where an E-Cad signal gap occurs (Fig. 1D and see Methods). This analysis established similar stereotypical dynamics of E-Cad, Myo-II, and local junction deformation following E-Cad gaps: 1) a local decrease of E-Cad and Myo-II, that was concomitant with junction lengthening; 2) an increase in Myo-II that overshot, around 200 s later, above its value prior to the local E-Cad decrease; and 3) junctional shortening within the next 100–200 s as Myo-II, E-Cad, as well as junctional length, recovered close to their original values. Importantly, and confirming previous results in the dorsal thorax and other epithelial tissues ^53–55^, we found that upon a partial reduction of E-Cad levels by RNAi (*e*-*cad^RNAi^*), cortical Myo-II levels increased, suggesting that an E-Cad decrease is sufficient to promote a local Myo-II increase (Extended Data Fig. 1D,E). Collectively, these results showed that junction lengthening and shortening dynamics concur with the formation and closure of local E-Cad signal gaps associated with stereotypical Myo-II dynamics. To understand how this cycle arises, and whether it could robustly mediate junctional stability (hereafter referred to as junction length homeostasis), we then sought to model it theoretically.

### An emergent response to passive dilution can maintain junction length homeostasis

For this, we developed a minimal active gel model for the dynamics of a junction of fluctuating length *l*(*t*) and concentration *m*(*t*) of Myo-II (which sets junctional contractility, see SI Theory, Fig. 2A). However, as previously described ^56–58^, this model alone, in the absence of any stabilizing feedback, typically leads to unstable junctions: junctional stretching leads to Myo-II dilution, leading in turn to smaller contractile forces that promote further junctional elongation. We therefore added E-Cad dynamics *c*(*t*) to the model, with the assumption, based on our results and the literature ^51,53–55^, that it modulates the steady-state Myo-II levels with coupling strength *γ* (this model is hereafter referred to as the E-Cad/Myo-II model). Altogether, the equations then read:

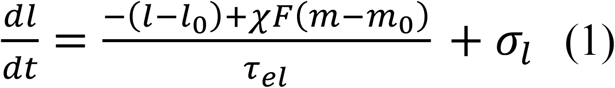

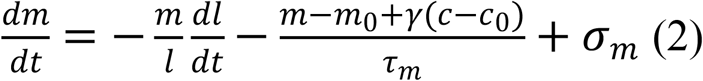

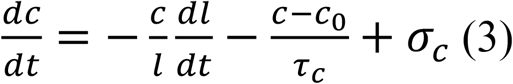

**Figure 2.**
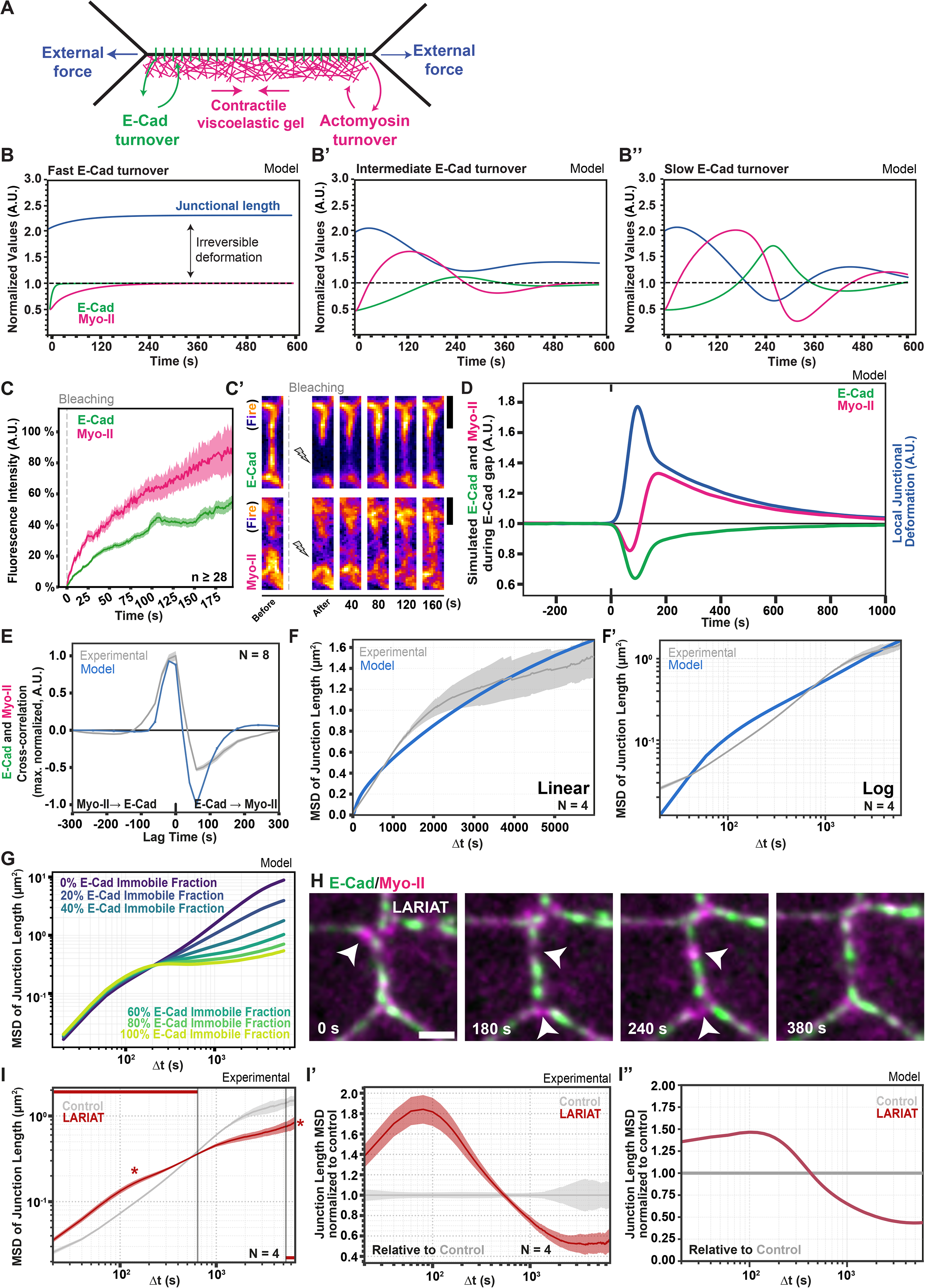
Theoretical model of emergent junction length homeostasis via differential dilution of junctional components. **(A)** Schematic of the theoretical model of an epithelial junction. Junction length is set by a balance of forces between junctional viscoelasticity, active Myo-II contractility and stochastic external forces on the junction. We consider that both E-Cad and Myo-II levels can vary due to either turnover or dilution/concentration from length fluctuations. **(B-B’’)** Simulated temporal profiles of junctional length, E-Cad levels and Myo-II levels after an abrupt junctional deformation from its homeostatic value of 1. If E-Cad turns much faster than Myo-II (B), junctional length does not go back to its homeostatic value. Intermediate values of E-Cad turnover allow for a mode of dynamical stabilization where irreversible length deformation is minimized (B’), while very slow values of E-Cad turnover predict junctional length oscillations (B’’). (**C,C’**) FRAP recovery curves of E-Cad:GFP and Myo-II:GFPx3 are shown as percentage of pre-bleach intensity (mean ± SEM). Time 0 corresponds to bleaching (vertical dashed line). Representative examples of E-Cad and Myo-II FRAP are shown in (C’). Arrowheads indicate FRAP regions on cell junctions. Myo-II and E-Cad are displayed with Fire LUT in (C’). n indicates the lowest number of FRAP measurements among the conditions shown. **(D)** Simulated dynamics of junction deformation, E-Cad levels, and Myo-II levels after an E-Cad gap in the best-fit simplified model of junction length homeostasis (see SI Theory). **(E)** Normalized cross-correlation function of E-Cad and Myo-II across the entire junction kymograph, in experiments (mean ± SEM) and theory. Minimum number of junctions analyzed per animal was 161 junctions. (**F,F’**) MSD of junction length over time (mean ± SEM), in experiments and theory, in linear **(F)** and log-log (F’) plots. Minimum number of junctions analyzed per animal was 187 junctions. **(G)** Theoretical predictions for how junction length MSD changes with varying immobile fractions of E-Cad. More immobile E-Cad predicts both increased short-term fluctuations and lower long-term fluctuations. **(H)** Time-lapse of E-Cad:GFP and Myo-II:mKate2x3 junction upon E-Cad:GFP clustering using the LARIAT optogenetic system. LARIAT was activated by 488 nm exposure 10 min before t = 0 s. **(I-I’’)** Experimental junction length MSD in LARIAT (mean ± SEM), shown without normalization to control junction length MSD (I), normalized to control junction length MSD at each time interval (I’), and as normalized simulated junction length MSD (I’’). Statistical significance was assessed by comparing the mean short-timescale MSD (first 10%; 0-640 s) and long-timescale MSD (last 20%; 3960-6620 s) using Welch’s t-test. Minimum number of junctions analyzed per animal was 187 junctions. Scale bars: 2 µm (C’,H). Time is shown in seconds (s). Unless otherwise indicated, n and N indicate the number of junctions and animals, respectively. Asterisks indicate statistical significance: *p < 0.05, **p < 0.01, ***p < 0.001; ns, not significant

The first equation is the force balance on the junction: *χ* is the strength of active contractility which depends on Myo-II concentration *F*(*m*), and *τ*_*el*_ is the timescale of short-term junctional relaxation, while *l*_0_ is the equilibrium length of the junction, which can remodel in time in a viscoelastic manner over a minute timescale,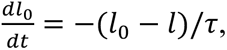 as previously measured experimentally ^59^ (see SI Theory for details). The next two equations are the conservation equations for Myo-II and E-Cad concentrations (with subscripts *m* and *c*), which can change owing to two processes: turnover on the timescale *τ*_*m*,*c*_, with associated noise *σ_i_*, but also junctional length changes 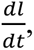 which tend to dilute/concentrate local Myo-II and E-Cad. As detailed further in SI Theory, this latter term is typically neglected in models of junction mechanics ^23,59^, but is important upon fast junctional length changes, and can naturally account for the near-instantaneous anticorrelation between junction length and Myo-II/E-Cad observed experimentally (Fig. 1B).

We first investigated the dynamics of this mechano-chemical model in response to an abrupt junctional lengthening. As expected, junction lengthening was associated with a near-instantaneous drop of both Myo-II and E-Cad arising from passive dilution, and we tested whether the model could also reproduce a return to the initial junction length. Incorporating the proposed E-Cad to Myo-II feedback confers stability to the system: as junction stretching dilutes E-Cad, this temporarily lifts the negative feedback on Myo-II, which then counteracts its own dilution. Notably, this compensatory mechanism from passive dilution does not require any explicit mechano-sensing terms. Furthermore, we found that this only occurs if the *τ*_*c*_ is large compared with the Myo-II turnover *τ*_*m*_ (Fig. 2B-B’’). Indeed, for fast turnover, E-Cad recovers too quickly to its initial concentration, providing little feedback to Myo-II (Fig. 2B). Less intuitively, for very slow E-Cad turnover, although the dilution of E-Cad acts as a long timescale memory of the original junction length prior to stretching, the model predicts a strong Myo-II overshoot, resulting in an oscillatory instability that prevents rapid return to homeostatic junctional lengths and can lead in extreme cases to unstable junctional collapse (Fig. 2B’’). This incompatibility between achieving both efficient return to a homeostatic length value and junction stability holds for most parameters of the model considered, as well as for different extensions of the model (see below and SI Theory and Extended Data Fig. 2A-G). We also found similar results when taking into account the possibility of differential dilution arising from different E-Cad immobile fractions, rather than from different turnover times (Extended Data Fig. 2C): (i) an intermediate immobile E-Cad fraction is highly beneficial to junctional length homeostasis; (ii) at a low E-Cad immobile fraction, there is little memory of past deformations over long time scales; (iii) but a high immobile fraction can result in Myo-II overshoot and short-term junctional oscillations (see SI Theory). Thus, our modelling approach suggests an efficient self-organized mechanism for junctional stability, in which junctional responses to stretch do not rely on any particular force sensor, but instead emerges due to the differential dilution of positive and negative regulators of Myo-II-dependent contractility.

We then aimed to test a key assumption of this model: E-Cad dynamics should be slower than those of Myo-II for any significant homeostatic regulation to occur. First, we performed FRAP experiments on Myo-II and E-Cad. This confirmed that Myo-II recovered fully within a few minutes (87 ± 3 s) with little immobile fraction, whereas E-Cad showed an intermediate immobile fraction (45 ± 2%) that did not recover over the timescale of the experiments, consistent with previous results ^53,60–62^ (Fig. 2C,C’). We then proceeded to constrain the parameters of the model beyond E-Cad and Myo-II dynamics. In particular, we used laser ablations to infer the time scale *τ*_*el*_ of junctional relaxation as *τ*_*el*_ = 5 ± 2 s (Extended Data Fig. 2G-H’), a value consistent with previous measurements^59^ and the correlation between junctional length and Myo-II level over time to obtain the rescaled contractility parameter *χ* = 0.30 ± 0.05 (Extended Data Fig. 2I, see SI Theory).

We then proceeded to quantitatively test the model with this parameter set, keeping the strength of the E-Cad to Myo-II feedback *γ* as the only fitting parameter. First, we imposed an abrupt force on the junction, causing lengthening and, thus, mimicking the formation of an E-Cad gap due to dilution. We found that the model could well reproduce the recovery of junctional length, as well as Myo-II and E-Cad levels (Fig. 2D), with *γ* = 1.5. Indeed, it could predict well the magnitude of the initial decrease in both E-Cad and Myo-II levels (with a stronger dilution of E-Cad), but also the timescale of Myo-II overshoot (at around 250 s) and of the return to initial values of E-Cad levels (closure of the gap on timescales of 400 s), as the junction recovers to its pre-gap value. Second, to probe the long-term dynamics and stochasticity of junction length regulation, we ran the model over long timescales without imposing a specific external force at t = 0, instead letting the noisy system evolve dynamically (see SI Theory for details). Mirroring the experimental quantifications, we then computed auto- and cross-correlation functions of junctional length, E-Cad and Myo-II levels (see SI Theory). These simulations showed good agreement with the experimental auto- and cross-correlations (resp. R^2^=0.95 and R^2^=0.66): both E-Cad and Myo-II exhibited anticorrelations with junctional length at small time delays (Fig. 1B and Extended Data Fig. 2J,K). Furthermore, in both model and experiments, the maximal anticorrelation with length peaked at dt = 0 s for E-Cad (passive dilution effect), but around dt = -20 s for Myo-II, consistent with a delay between Myo-II recruitment and junctional shrinkage due to the timescale of junctional elasticity (Fig. 1B and Extended Data Fig. 2J,K). The cross-correlation between E-Cad and Myo-II, in both model and experiments, also showed a positive correlation at delays of dt = - 20 s (the same mechanism as above, whereby Myo-II increase leads to junctional shrinkage and passive E-Cad increase), but also an anticorrelation at longer delays of dt = 60 s (Fig. 2E). Third, we sought to test our hypothesis of junctional length homeostasis more quantitatively, by computing the mean-square displacement (MSD) of junctional length across all parameter regimes of the model. For unregulated junctional length, random fluctuations should build up over time, leading to long-term diffusive behavior (corresponding to an MSD scaling linearly with time, see Extended Data Fig. 2A-D and SI Theory for details). In the presence of feedback and an E-Cad immobile fraction, homeostatic memory manifested itself as a sub-diffusive scaling of the MSD, which tended to plateau at long timescales, in agreement with the model (R^2^=0.94, Fig. 2F-G).

Overall, the mechano-chemical model suggested that junctional homeostasis can be achieved by differential dilution of positive and negative regulators of contractility as long as the negative regulators have slower turnover. In the current model, the negative regulator is E-Cad, while the positive regulator remains to be identified. To experimentally test the model prediction about the role of slow turnover of negative regulators, we sought to further increase the difference between E-Cad and Myo-II turnover. For this, we performed optogenetic clustering of E-Cad:GFP using LARIAT ^54,63^ (Supplementary Video 2), which nearly fully abrogated E-Cad:GFP turnover dynamics, as assessed by FRAP experiments (Extended Data Fig. 2L). We then ran our fully parameterized model with immobile E-Cad to match this experimental condition, which predicted better junctional homeostasis over long timescales, as characterized by a lower long-term MSD for junctional length, but with higher short-term fluctuations (Fig. 2G), as discussed above. Performing junctional tracking in the LARIAT condition agreed with this theoretical prediction (R^2^=0.59, Fig. 2H-I’’). This is best visualized by plotting the LARIAT MSD normalized to the control condition at each time interval (Fig. 2I’,I’’). In both data and theory, LARIAT had larger noise at short-time scale (below 500 s) and smaller noise at long-time scale (above 500 s), underlining that the robust long-term junction homeostasis in LARIAT conditions is not simply due to lower short-term noise.

Altogether, these findings were consistent with an emergent mechano-response process, arising from the passive dilution of regulators with intrinsically different turnover rates. However, several key questions raised by the model remained open. From a more mechanistic perspective, what are the positive regulators of contractility and are their dynamics fast compared with E-Cad? Can we perturb these regulators to test more functionally whether the proposed mechano-chemical feedback is key for long-term junctional homeostasis?

### Junction length stability is controlled by the dynamics of the RhoGEF Cyst

To identify the positive regulators of contractility in junctional homeostasis, we first considered the classical AJ mechano-sensors, Vinculin and Ajuba (Jub, *Drosophila* LMD1), which are known to locally increase actomyosin at AJs under mechanical loads ^17,18,26,38,39,42,44,47,50,51,64–68^. However, we did not detect significant enrichment within gaps or anticorrelation with E-Cad that would be expected for a positive regulator of contractility in junctional homeostasis (Extended Data Fig. 3A-E). In addition, loss of Jub or Vinc function did not affect the stereotypical E-Cad and Myo-II dynamics (Extended Data Fig. 3F) and did not increase junction fluctuations as revealed by junction length MSD analyses (Extended Data Fig. 3G). These results led us to envision that distinct mechanisms may contribute to junction length homeostasis.

We previously reported that during cytokinesis, the decrease in E-Cad promoted by contraction of the cytokinetic ring was associated with a local increase in the RhoGEF Cyst and Myo-II ^54^. Additionally, the optogenetic clustering of E-Cad using LARIAT promoted local E-Cad decreases where Cyst accumulated ^54^. Therefore, Cyst is a potential candidate to understand how local E-Cad dilution can lead to the emergence of a mechano-chemical feedback between E-Cad and Myo-II. To analyze this hypothesis, we first investigated the co-dynamics of Cyst and E-Cad at cell junctions. In contrast to Vinc and Jub (Extended Data Fig. 3A-E), we observed that the local decreases of E-Cad were associated with a local increase of Cyst (Fig. 3A). We then quantified the dynamics of E-Cad, Cyst, and Myo-II during E-Cad gap formation and found that E-Cad gaps were followed by a rapid increase in Cyst signal, which preceded the Myo-II peak (Fig. 3B). This was also observed by cross-correlating the Cyst signal with Myo-II or E-Cad in the analysis of whole junction dynamics (Fig. 3C,D), suggesting that Cyst could promote the local increase of Myo-II in E-Cad gaps. Consistent with this hypothesis, loss of Cyst function reduced Myo-II junctional level and FRAP experiments revealed that Cyst recovered even faster than Myo-II (Extended Data Fig. 3K-M). Furthermore, partial E-Cad knockdown (*e*-*cad^RNAi^*) led to a junctional increase in Cyst, as observed for Myo-II (Extended Data Fig. 3H-H’’) and, notably, reducing Cyst function in *e*-*cad^RNAi^* clones abrogated the Myo-II accumulation observed in these clones (Extended Data Fig. 3I-J’). Together, these results agreed with the notion that Cyst acts downstream of the local E-Cad decrease to promote Myo-II accumulation. After incorporating Cyst into our mechano-chemical model (see SI Theory for details on equations and parameter fitting, Extended Data Fig. 3R-S and Extended Data Fig. 4 for detailed sensitivity analysis), we first tested whether we could reproduce Cyst dynamics during E-Cad gap appearance and closure. As observed experimentally, in the model, Cyst showed little dilution and rapidly increased as E-Cad was diluted, with a peak within 100-150s after the onset of the E-Cad gap, before that of Myo-II (peak at 200 s, Fig. 3E). This resulted in an efficient closure of the E-Cad gap by 300s, in both model and experiments (Fig. 3E). We then turned to computing the auto- and cross-correlation functions of the full kymograph between Cyst, Myo-II, E-Cad and junctional length, in both model and experiments (resp. R^2^=0.84 and R^2^=0.59), which again showed a good agreement despite the simplicity of the model (Fig. 3C,D and Extended Data Fig. 5A-J). Notably, the model including Cyst captures better the strength of the E-Cad/Myo-II anti-correlation peak than the E-Cad/Myo-II only model.

**Figure 3.**
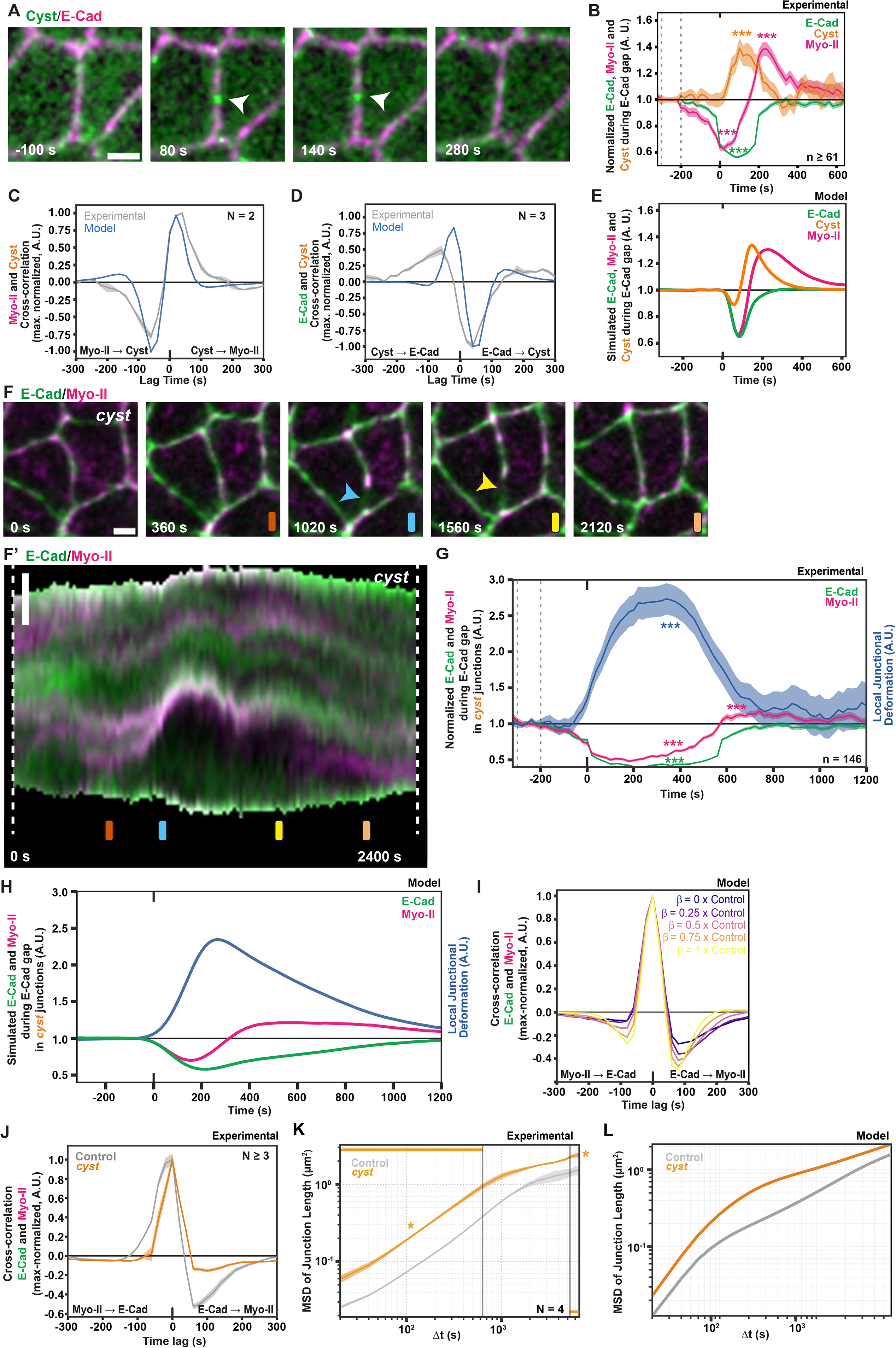
The RhoGEF Cyst controls E-Cad, Myo-II, and junction length dynamics. **(A)** Time-lapse images of Cyst:GFP and E-Cad:mKate2x3. White arrowheads mark the position of the E-Cad gap and Cyst accumulation. Timestamps are aligned to E-Cad gap onset. **(B)** Normalized E-Cad, Myo-II, and Cyst intensities (median ± SEM) aligned to local E-Cad gap events (t = 0). E-Cad, Myo-II, and Cyst intensities were normalized to their median baseline values between -300 and -200 s, indicated by dotted lines. Statistical significance was assessed using paired Wilcoxon signed-rank tests against baseline. n: E-Cad gaps (61 for E-Cad and Cyst, and 163 for E-Cad and Myo-II). Minimum number of animals analyzed for this quantification was 3. **(C)** Comparison of model predictions (blue) and experimental data (grey) for cross-correlations between Myo-II and Cyst (mean ± SEM). Minimum number of junctions analyzed per animal was 116 junctions. **(D)** Comparison of model predictions (blue) and experimental data (grey) for cross-correlations between E-Cad and Cyst (mean ± SEM). Minimum number of junctions analyzed per animal was 119 junctions. **(E)** Simulated dynamics of E-Cad, Myo-II, and Cyst levels after an E-Cad gap in the best-fit full model of junction length homeostasis (see SI Theory). **(F,F’)** Time-lapse images of E-Cad:GFP and Myo-II:mKate2x3 at a junction in a *cyst* mutant clone. Colored bars (orange, blue, yellow, peach) above each panel indicate the corresponding time points in the kymograph in (F’). Blue and yellow arrowheads mark the formation of the E-Cad gap, the difficulty in detecting a clear Myo-II:mKate2x3 accumulation, and the persistence of the E-Cad gap 680 s later. Kymograph of E-Cad:GFP and Myo-II:mKate2x3 along the junction shown in (F) over 2400 s (40 min, F’). Colored bars indicate the time points displayed in (F). **(G)** Normalized E-Cad, Myo-II, and local junction deformation (median ± SEM) aligned to local E-Cad gap events (t = 0) in *cyst* mutant junctions. E-Cad and Myo-II intensities, as well as local junction deformation, were normalized to their median baseline values between -300 and -200 s, indicated by dotted lines. Statistical significance was assessed using paired Wilcoxon signed-rank tests against baseline. n: total E-Cad gaps. Minimum number of animals analyzed for this quantification was 6. **(H)** Simulated dynamics of local junction deformation, E-Cad, and Myo-II in cyst mutant junctions after an E-Cad gap (see SI Theory). **(I)** Simulation of the normalized cross-correlation between E-Cad and Myo-II intensities, for decreasing values of the feedback strength *β* compared to the best-fit value from control junctions. **(J)** Normalized cross-correlation between E-Cad and Myo-II intensities (mean ± SEM) in control and *cyst* mutant junctions. Cross-correlation values were normalized to the maximum cross-correlation value. Minimum number of junctions analyzed per animal was 161 junctions. **(K)** MSD of junction length (mean ± SEM) over time intervals for control and *cyst* mutant junctions. Statistical significance was assessed by comparing the mean short-timescale MSD (first 10%; 0-640 s) and long-timescale MSD (last 20%; 3960-6620 s) using Welch’s t-test. Minimum number of junctions analyzed per animal was 187 junctions. **(L)** Predicted MSD of junction length over time intervals for control and *cyst* mutant junctions. Scale bars: 2 µm (A,F,F’). Time is shown in seconds (s). Unless otherwise indicated, n and N indicate the number of junctions and animals, respectively. Asterisks indicate statistical significance: *p < 0.05, **p < 0.01, ***p < 0.001; ns, not significant.

To functionally test whether Cyst controls E-Cad, Myo-II, and junction length dynamics, we then recorded the dynamics of E-Cad, Myo-II, and junction length in *cyst* mutant cells (Supplementary Video 3). We found that the loss of Cyst function caused the formation of larger and longer-lasting gaps in the E-Cad signal (Fig. 3F,F’ and Extended Data Fig. 3N-N’’). Within such gaps, the recovery of the Myo-II signal was greatly reduced (Fig. 3G). Therefore, removing Cyst activity perturbed junction length dynamics by delaying the closure of E-Cad gaps (Fig. 3G and Extended Data Fig. 3N-N’’). Importantly, as observed for E-Cad signal gaps in the control condition, the E-Cad gaps observed in *cyst* mutant cells were not associated with membrane rupture (Extended Data Fig. 3O). Next, we simulated junction dynamics with impaired mechano-chemical feedback strength to mimic the *cyst* mutant condition. We also repeated FRAP and laser ablation experiments in *cyst* mutant cells to constrain turnover and contractility parameters in this condition (Extended Data Fig. 3P,Q and see SI Theory). In response to an E-Cad gap, simulations using *cyst* mutant parameters reproduce the weaker Myo-II overshoot and slower junctional contraction seen experimentally (Fig. 3H). Furthermore, computing auto- and cross-correlation functions across the full stochastic simulations also showed good agreement with data (resp. R^2^=0.88 and R^2^=0.78), in particular a weaker anticorrelation between E-Cad and Myo-II (Fig. 3I,J and Extended Data Fig. 5K-P). Finally, to explore the consequences of Cyst loss-of-function for junction length homeostasis, we compared the junction dynamics using MSD analysis of junction length in control and Cyst loss-of-function conditions and found a larger MSD at all time-points for *cyst* mutant junctions (Fig. 3K), which was well recapitulated by our model (Fig. 3L, R^2^=0.86).

As an orthogonal test of the model, we also explored its core assumption that Cyst passively responds to the dilution of its negative regulator E-Cad and, thus, that cell contractility *per se* is not necessary for active Cyst recruitment. Indeed, we found that *rock^RNAi^*did not decrease Cyst accumulation at the AJ (Extended Data Fig. 3T-T’’), nor did it modify the E-Cad and Cyst cross-correlation, as predicted by the model (Extended Data Fig. 3U). Therefore, the previously parametrized model could explain the auto- and cross-correlation functions observed in *rock^RNAi^* (resp. R^2^=0.96 and R^2^=0.48), simply by lowering the contractility parameter *χ* and keeping all other parameters constant (Extended Data Fig. 5Q-V). To directly test that the inhibitory effect of E-Cad on Cyst did not require contractility, we examined *e-cad^RNAi^*, *rock^RNAi^*clones, and confirmed that Cyst remained enriched at the cell-cell junction in these clones (Extended Data Fig. 3V,V’). This further illustrates that Cyst-dependent response to E-Cad gap is distinct from classical mechano-sensing pathways that depend on cell contractility and E-Cad ^17,18,26,38,39,42,44,47,50,64–67^. Overall, we concluded that Cyst fulfils the theoretical model requirements for a positive regulator of Myo-II with fast dynamics compared with E-Cad, and that this underlies junction length homeostasis.

### Impact of emergent mechano-response for long-term junctional and shape homeostasis

Next, we sought to test the model further by quantifying junctional length homeostasis upon mechanical challenges. We previously reported that down-regulating α-Spectrin (α-Spec), a key cytoskeletal component of cell-cell junctions, leads to abrupt junction elongation ^67^, and our MSD analysis confirmed significantly larger short-term length fluctuations, with numerous gaps in E-Cad signal (Fig. 4A-B and Supplementary Video 4). As in control cells, these junctional gaps were not associated with membrane rupture (Extended Data Fig. 6A). To check that other junctional parameters were not affected, we performed FRAP and laser ablation experiments in *α-spec^RNAi^* cells, and found that E-Cad turnover dynamics and actomyosin contractility were not different from control cells (Extended Data Fig. 6B-D). This made *α-spec^RNAi^*cells an ideal condition to investigate how increasing short-term mechanical noise (parameter *σ*_*m*_) affects long-term junction length homeostasis. Interestingly, the model predicted that in contrast to Cyst loss of function, which showed high MSD at both short and long timescales, *α-spec^RNAi^*cells should show noise-buffering with a convergence towards control long-term fluctuations (Fig. 4C). Notably, this was closely mirrored experimentally (R^2^=0.94), with long-term MSD in *α-spec^RNAi^* cells converging to values close to control condition (Fig. 4B). Furthermore, we found that (i) the junction gaps observed in *α-spec^RNAi^* tissue were quickly repaired following E-Cad dilution as in control conditions, with Cyst and Myo-II recruitment within the E-Cad gap (Fig. 4A and Extended Data Fig. 6E-H); and (ii) auto-and cross-correlation functions between E-Cad and Myo-II, and Cyst were qualitatively unaffected and followed model predictions (resp. R^2^=0.96 and R^2^=0.84, Fig. 4D and Extended Data Fig. 7A-I). Furthermore, reducing Cyst function in *α-spec^RNAi^* led to additive defects in junction length homeostasis as revealed by MSD analysis (Supplementary Video 5), again in agreement with the model (R^2^=0.72, Fig. 4B,C and Extended Data Fig. 6I,I’). We therefore concluded that the emergent mechano-response via differential dilution enables long-term junction homeostasis even in the presence of enhanced short-term fluctuations.

**Figure 4.**
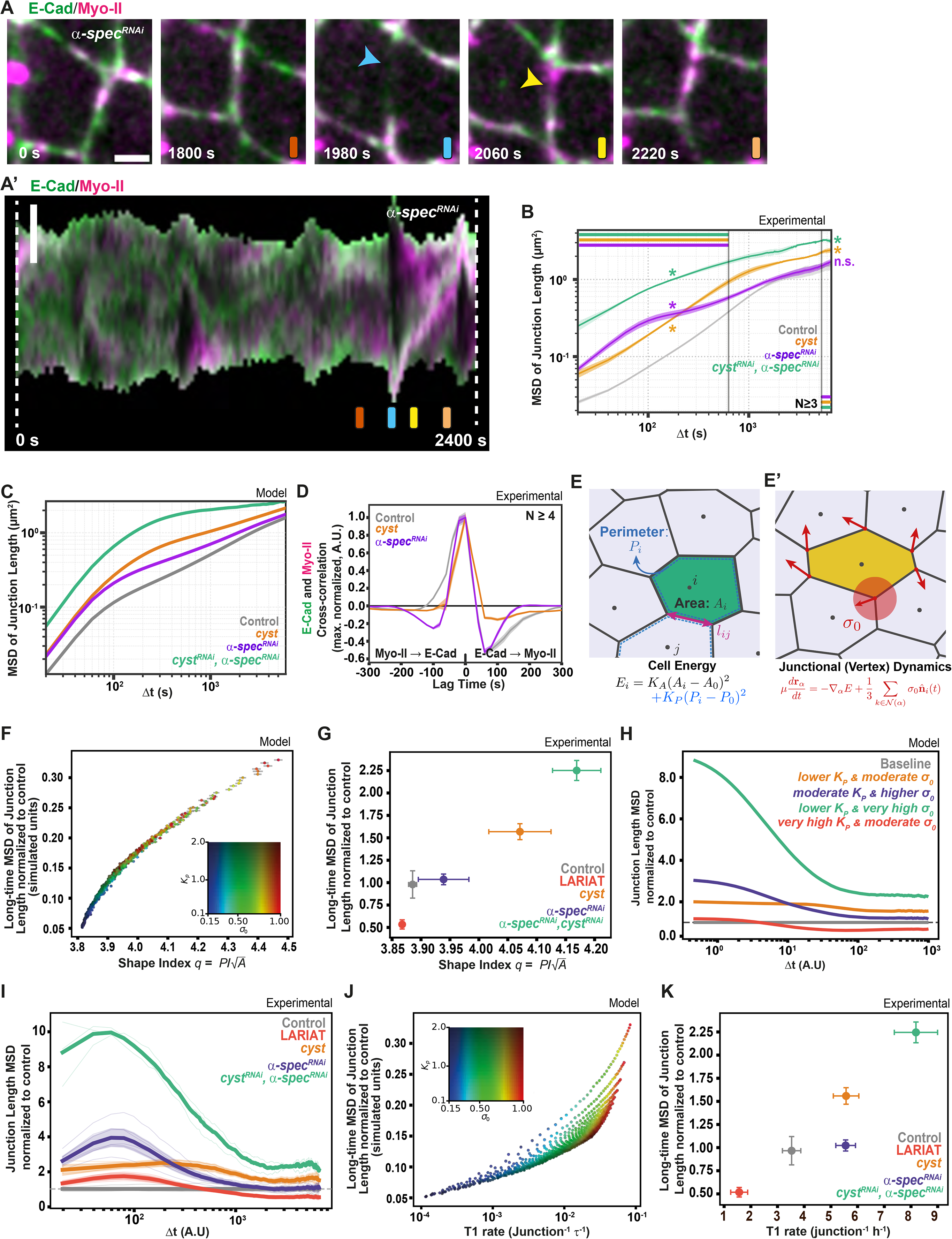
α-Spectrin controls short-term junction dynamics. **(A,A’)** Time-lapse images of E-Cad:GFP and Myo-II:mKate2x3 at a junction in an *α-spec^RNAi^* clone. Colored bars (orange, blue, yellow, peach) above each panel indicate the corresponding time points in the kymograph in (A’). Blue and yellow arrowheads mark the position of the E-Cad:GFP gap and the subsequent Myo-II:mKate2x3 accumulation. Kymograph of E-Cad:GFP and Myo-II:mKate2x3 along the junction shown in (A) over 2400 s (40 min) is shown in (A’). **(B)** MSD of junction length over time intervals (t) in control, *cyst mutant*, *α-spec^RNAi^*, and *α-spec^RNAi^*, *cyst^RNAi^* junctions. Statistical significance was assessed by comparing the mean short-timescale MSD (first 0-10%; 0-640 s) and long-timescale MSD (last 20%; 3960-6620 s) using Welch’s t-test. Minimum number of junctions analyzed per animal was 106 junctions. **(C)** Simulated MSD of junction length over time intervals in control, *cyst mutant*, *α-spec^RNAi^* and *α-spec^RNAi^*, *cyst^RNAi^* junctions. **(D)** Normalized cross-correlation function of E-Cad and Myo-II intensities (mean ± SEM) across control, *cyst* mutant, and *α-spec^RNAi^* junction kymographs. Cross-correlation values were normalized to the maximum cross-correlation value. Minimum number of junctions analyzed per animal was 106 junctions. **(E-E’)** Schematic of the active vertex model (AVM). Left: a representative cell *i*, with neighboring cell *j*, showing cell area *A_i_*, perimeter *P_i_*, and shared junction length *l_ij_* between cells *i* and *j*. Cell geometry is governed by the energy *E_i_* = *K_A_*(*A_i_* – *A*_0_)^2^ + *K_p_*(*P_i_* – *P*_0_)^2^ (E). Right: schematic of the isotropic active noise term *σ*_0_ at a representative vertex *α* (red dot); short arrows indicate independent noise contributions at each cell vertex, and the shaded red circle indicates the isotropic (undirected) character of the noise at *α*. Vertex motion follows 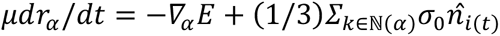 (E’). See SI Theory for full model details. **(F)** Simulated long-time MSD of junction length (mean ± SEM, simulation units), as a function of the observed shape index 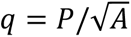 in AVM simulations (preferred shape index parameter *q*_0_ = 3.8). Each point represents one (*K_p_, σ*_0_) combination; color encodes both parameters (hue: *σ*_0_, blue to red, from 0.15 to 1.00; brightness: *K_p_*, light to dark, from 0.1 to 2.0). **(G)** Long-time MSD of junction length as a function of shape index *q* in control, LARIAT, *cyst mutant*, α-*spec^RNAi^*, and α-*spec^RNAi^*, *cyst^RNAi^* conditions. One point is shown per condition; error bars show the SD of *q* (x-axis) and the SEM of long-time junction MSD across replicates (y-axis). MSD values for all conditions are normalized to control MSD. **(H)** Simulated MSD of junction length normalized to the baseline parameter set (*K*_p_ = 0.30, *σ*_0_ = 0.275) over time intervals (simulation units). Dashed line at ratio = 1 marks the baseline. Five parameter combinations are shown: baseline (*K_p_* = 0.30, *σ*_0_ = 0.275); lower *K_p_*, moderate *σ*_0_ (*K_p_* = 0.10, *σ*_0_ = 0.375); moderate *K_P_*, higher *σ_0_* (*K_p_* = 0.50, *σ*_0_ = 0.500); lower *K_P_*, very high *σ_0_* (*K*_p_ = 0.10, *σ*_o_ = 0.775); and higher *K_P_*, moderate *σ_0_* (*K_p_* = 2.00, *σ*_0_ = 0.325). Higher *σ*_0_ enhances short-timescale fluctuations, while higher *K_p_* lowers long-timescale fluctuations. **(I)** Normalized junction length MSD over time intervals in control, LARIAT, *cyst*, *α-spec^RNAi^*, and *α-spec^RNAi^*, *cyst*^RNAi^ junctions. Normalization is done to the control MSD for each time interval. Thin lines show individual replicates; thick lines show the condition mean; shaded bands show the propagated error. Dashed line at ratio = 1 marks the control reference. Minimum number of junctions analyzed per animal was 106 junctions. **(J)** Simulated long-time MSD of junction length (mean ± SEM, simulation unit) as a function of T1 transition rate (junction⁻¹ τ⁻¹) in AVM simulations. As in (F), each point represents one (*K_p_, σ*_0_) combination, colored using the same hue (*σ*_0_) and brightness (*K_p_*) as in (F). **(K)** Long-time MSD of junction length as a function of T1 transition rate in Control, LARIAT, *cyst*, *α-spec^RNAi^* and *α-spec^RNAi^, cyst^RNAi^* conditions, colored as in (I). MSD values for all conditions are normalized to control MSD. Minimum number of junctions analyzed per animal was 106 junctions. Scale bars: 2 µm (A, A’). Time is shown in seconds (s). Unless otherwise indicated, n and N indicate the number of junctions and animals, respectively. Asterisks indicate statistical significance: *p < 0.05, **p < 0.01, ***p < 0.001; ns, not significant.

Finally, we asked whether differences between control, LARIAT, *cyst* mutant, *α-spec^RNAi^*and *cyst^RNAi^*, *α-spec^RNAi^*junctional dynamics also had consequences for cell shape. Vertex models assume tight shape regulation of each cell to a reference shape index *q_0_* (i.e., the cell’s perimeter normalized by area, Fig. 4E,E’), with an associated perimeter elasticity coefficient *K_P_*, which is predicted to be dependent on Cyst function (see also SI Theory). Running active Vertex model simulations revealed that as *K_P_* is decreased, the observed cell shape index *q* increases away from its target value *q_0_*, as expected from previous works on solid-fluid transitions ^69,70^ (Fig. 4F and Extended Data Fig. 7J-O, see SI Theory for details). Computing the same metrics for experimental data confirmed this prediction, with *cyst* mutant cells (lower *K_P_*) having a larger shape index *q* than control and *α-spec^RNAi^* cells (Fig. 4G). To test for the generality of this finding, we went back to the simulations and found that we could derive a generic master curve relating long-term junctional MSD to static cell shape *q* (Fig. 4F), which holds both when varying perimeter elasticity coefficient *K_P_* and mechanical noise *σ*_7_. Strikingly, computing both metrics across the control, LARIAT, *cyst* mutant, *α-spec^RNAi^* as well as *cyst^RNAi^*, *α-spec^RNAi^*conditions reproduces the trend of this simple prediction (Fig. 4G and Extended Data Fig. 7P). We also found that we could capture both long-term and short-term differences in junctional MSD between control cells and the different perturbations with the Vertex model (Fig. 4H,I), showing that we can map our junctional parameters on tissue-level properties (see also SI Theory). As a final test of whether junctional length homeostasis can predict tissue-scale features such as tissue fluidity, we found in the Vertex model a relationship between long-term junctional MSD and the rate of junctions undergoing a T1 transition (Fig. 4J). Computing both quantities across the control, LARIAT, *cyst* mutant, *α-spec^RNAi^*and *cyst^RNAi^*, *α-spec^RNAi^*conditions agrees with these predicted correlations (Fig. 4K and Extended Data Fig. 7Q). This showed that impaired junction length homeostasis impacts cell shape and tissue fluidity.

### Local decrease in aPKC levels controls Cyst recruitment and junction shortening

Having established the relevance of the mechano-chemical feedback for the control of junction, cell shape homeostasis and tissue fluidity, we then sought to analyze how Cyst is recruited to E-Cad gaps. We first analyzed the function of Crumbs (Crb) and Yurt (Yrt, known as Lulu1 and 2 in vertebrates), which have been reported to control Cyst membrane localization ^71^. While Crb knock-down (*crb^RNAi^*) led to a partial decrease in Cyst levels at the junction, we did not observe an obvious junctional Myo-II decrease or the formation of long-lasting E-Cad gaps in *crb^RNAi^* clones (Extended Data Fig. 8A-J). While we cannot fully exclude a role for Crb, we found that Yrt knock-down (*yrt^RNAi^*) led to a pronounced decrease in Cyst levels at the AJ (Fig. 5A,A’). Furthermore, *yrt^RNAi^* was associated with a decrease in junctional Myo-II and the formation of E-Cad gaps (Fig. 5B-B’’ and Extended Data Fig. 9A). Accordingly, in the absence of Yrt function, gap dynamics as well as the cross-correlation between E-Cad and Myo-II were changed as observed in *cyst* mutant tissues (Fig. 5C-D). Furthermore, reducing Yrt function in *e*-*cad^RNAi^*clones decreased the Myo-II accumulation observed in these clones (Extended Data Fig. 9B-C’). Last, Yrt dynamics were consistent with a function upstream of Myo-II, since Yrt rapidly accumulated in E-Cad gaps and it is characterized by both a high mobile fraction and turnover as shown by FRAP experiments (Fig. 5E,F and Extended Data Fig. 8J). Yrt membran localization is regulated by the activity of aPKC, which directly phosphorylates Yrt to prevent its membrane accumulation ^52,72,73^. Accordingly, Yrt membrane localization is an *in situ* readout of aPKC activity ^72,73^. This regulation has previously been proposed in the early *Drosophila* embryo as a key mechanism to establish the apical and basolateral epithelial domains ^72,74^. Yet, the role of this regulation in E-Cad dynamics is unknown. aPKC activity is regulated by its binding to Par6 ^52,75–79^. We therefore explored both aPKC and Par6 dynamics during E-Cad gap formation. We found that both aPKC and Par6 were mildly but significantly decreased within the E-Cad gap (Fig. 5G-H and Extended Data Fig. 9D-G). We then found that aPKC loss of function led to a strong apical junctional accumulation of Yrt and Cyst (Fig. 5I-I’’ and Extended Data Fig. 9H,H’) that is consistent with the strong accumulation of Myo-II and apical constriction previously reported upon aPKC loss of function ^80^. Notably, partial loss of E-Cad function using *e-cad^RNAi^* led to a reduction of aPKC level at cell junction (Extended Data Fig. 9I-J). To further test whether E-Cad gaps were sufficient to promote downregulation of aPKC activity, we analyzed whether optogenetically promoting local E-Cad decrease would be sufficient to promote Yrt accumulation. Using the LARIAT system, we clustered E-Cad:GFP to generate ectopic regions of E-Cad gaps. Strikingly, we found that Yrt accumulated in the induced regions of E-Cad decrease (Fig. 5J,J’). Collectively, these results suggested a model whereby the decrease in E-Cad induces a local aPKC activity decrease that controls Myo-II contractility via the rapid turnover and junctional accumulation of Yrt and Cyst.

**Figure 5.**
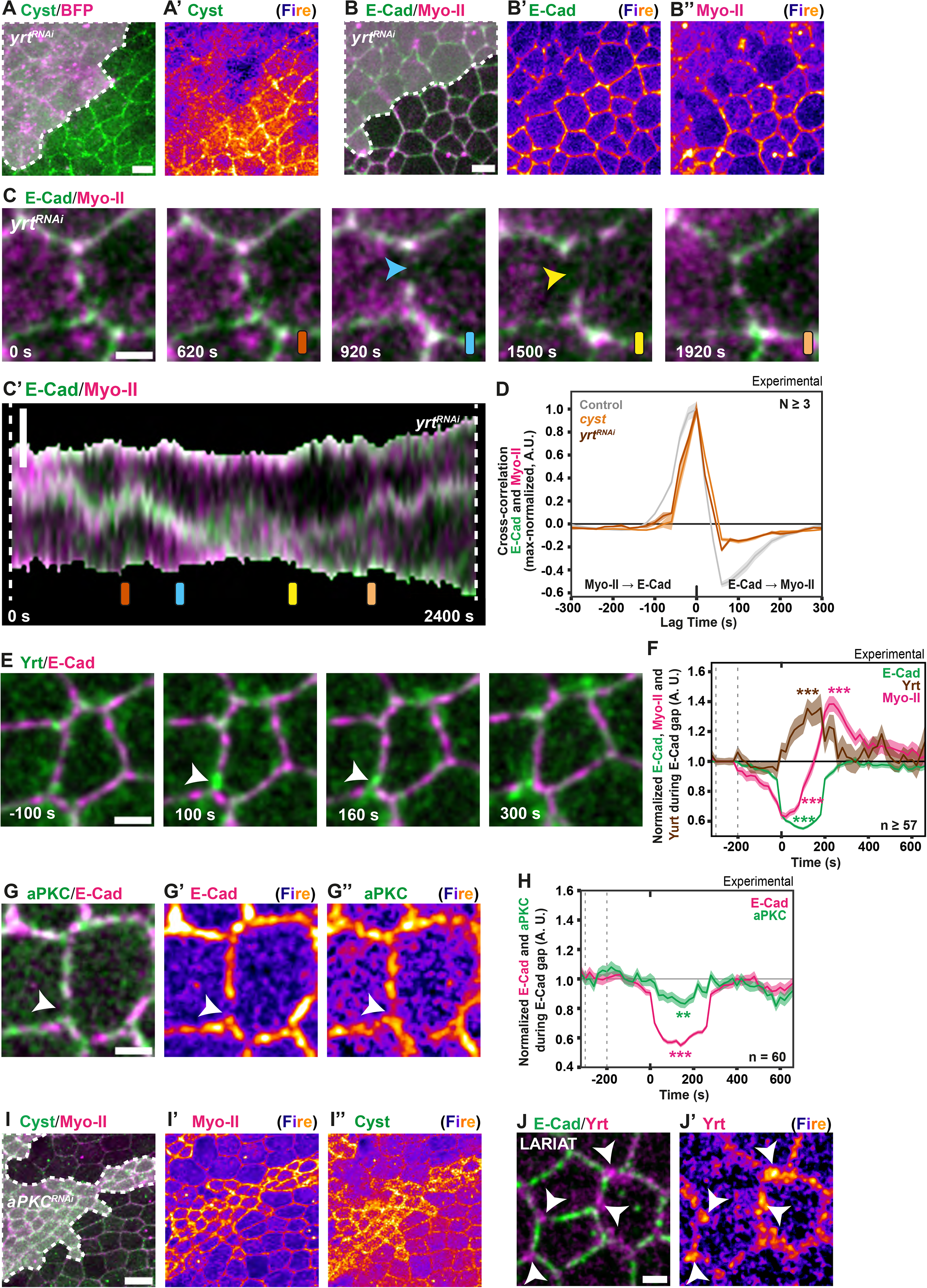
aPKC and Yrt control Cyst accumulation at E-Cad gaps. **(A, A’)** Cyst:GFP and CAAX:BFP2 in tissue with *yrt^RNAi^* cells. Dotted line (A) marks the position of *yrt^RNAi^* cells identified by the expression of CAAX:BFP2 (not shown). Cyst:GFP displayed with Fire LUT (A’). **(B-B’’)** E-Cad:GFP and Myo-II:mKate2x3 in tissue with *yrt^RNAi^*cells. Dotted line (B) marks the position of *yrt^RNAi^* cells identified by the expression of CAAX:BFP2 (not shown). E-Cad:GFP (B’) and Myo-II:mKate2x3 (B’’) displayed with Fire LUT. **(C, C’)** Time-lapse images of E-Cad:GFP and Myo-II:mKate2x3 in *yrt^RNAi^* tissue. Colored bars (orange, blue, yellow, peach) above each panel indicate the corresponding time points in the kymograph in (C’). Blue and yellow arrowheads mark the position of the E-Cad:GFP gap, the low level Myo-II:mKate2x3 accumulation, and the persistence of the E-Cad gap 880 s later. Kymograph of E-Cad:GFP and Myo-II:mKate2x3 along the junction shown in (C) over 40 min is shown in (C’). **(D)** Normalized cross-correlation between E-Cad and Myo-II intensities (mean ± SEM) in control, *cyst* mutant and *yrt^RNAi^* junctions. Cross-correlation values were normalized to the maximum cross-correlation value. Minimum number of junctions analyzed per animal was 82 junctions. **(E)** Time-lapse images of Yrt:GFPx3 and E-Cad:mKate2x3 in control tissue. White arrowheads mark E-Cad gap and local increase of Yrt. Timestamps are aligned to E-Cad gap onset. **(F)** Normalized E-Cad, Myo-II, and Yrt intensities (median ± SEM) aligned to local E-Cad gap events (t = 0). E-Cad, Myo-II, and Yrt intensities were normalized to their median baseline values between −300 and −200 s, indicated by dotted lines. Statistical significance was assessed using paired Wilcoxon signed-rank tests against baseline. n: E-Cad gaps (57 E-Cad gaps for E-Cad and Yrt from 3 animals, and 163 E-Cad gaps for E-Cad and Myo-II from 8 animals). **(G-G’’)** aPKC:GFP and E-Cad:mKate2x3 in control tissue (G), with aPKC:GFP (G’) and E-Cad:mKate2x3 (G’’) displayed with Fire LUT. White arrowhead marks E-Cad gap and the local decrease of aPKC signal. **(H)** Normalized E-Cad:mKate2x3 and aPKC:GFP intensities (median ± SEM) aligned to local E-Cad decrease events (t = 0). E-Cad and aPKC intensities were normalized to their median baseline values between -300 and -200 s, indicated by dotted lines. Statistical significance was assessed using paired Wilcoxon signed-rank tests against baseline. n: total E-Cad gaps. Minimum number of animals analyzed for this quantification was 3. **(I-I’’)** Cyst:GFP and Myo-II:mKate2x3 in tissue containing *aPKC^RNAi^* cells, identified by the expression of CAAX:BFP2 (I; dashed outline), Myo-II:mKate2x3 (I’) and Cyst:GFP (I’’) displayed with Fire LUT. **(J-J’)** E-Cad:GFP and Yrt:mKate2x3 in tissue where E-Cad:GFP clustering was induced using the LARIAT optogenetic system. LARIAT was activated by 488 nm exposure 10 min before t = 0 s. Scale bars: 5 µm (A,B,C,C’,G,J); 10 µm (I). Time is shown in seconds (s). Unless otherwise indicated, n and N indicate the number of junctions and animals, respectively. Asterisks indicate statistical significance: *p < 0.05, **p < 0.01, ***p < 0.001; ns, not significant.

### Emergent mechano-response to increased stress anisotropy

Lastly, we explored whether the proposed mechano-chemical feedback provides an emergent mechano-response to an increase in anisotropic stress. So far, our analyses had been performed between 18 and 20 h APF, a developmental stage at which the tissue is characterized by a low isotropic mechanical stress^81^. Laser ablation revealed that in the anterior domain of the thorax, mechanical stress increased in magnitude and anisotropy between 18 and 26 h APF, with stress becoming more pronounced along the AP axis (Fig. 6A-C). Using this endogenous mechanical stress increase, we explored the contribution of the proposed mechano-chemical feedback to a mechano-response. To this end, we compared E-Cad dynamics in control and *cyst* mutant tissues at 18 and 26 h APF (Extended Data Fig. 10A-C). In control tissues, the frequency and duration of E-Cad gaps were slightly increased between 18 and 26 h APF, with gaps being slightly more pronounced on junctions oriented along the AP axis at 26 h APF (Fig. 6D,E). As expected from the above results, we found that the frequency and duration of gaps were increased in *cyst* mutant cells at 18 h APF for junctions both parallel and orthogonal to the AP axis (Fig. 6D,E). More interestingly, at 26 h APF, (namely in the presence of anisotropic stress oriented along the AP axis), we observed that loss of Cyst function resulted in a drastic increase in E-Cad junction gaps along the AP axis (Fig. 6D,E). Together, these results indicated that the formation of E-Cad gaps can be induced by higher mechanical stress anisotropy and that Cyst contributes to the tissue’s mechanical response by buffering increases in junction length. Accordingly, we found that *cyst* mutant cells were more elongated than control cells along the AP axis at 26 h APF (Fig. 6F-G’). Altogether, we concluded that local dilution of E-Cad passively relieves the inhibition on lateral apicobasal polarity components, allowing the RhoGEF Cyst, with its fast turnover, to accumulate and increase contractility to buffer junction length fluctuations and prevent junction elongation under mechanical stress.

**Figure 6.**
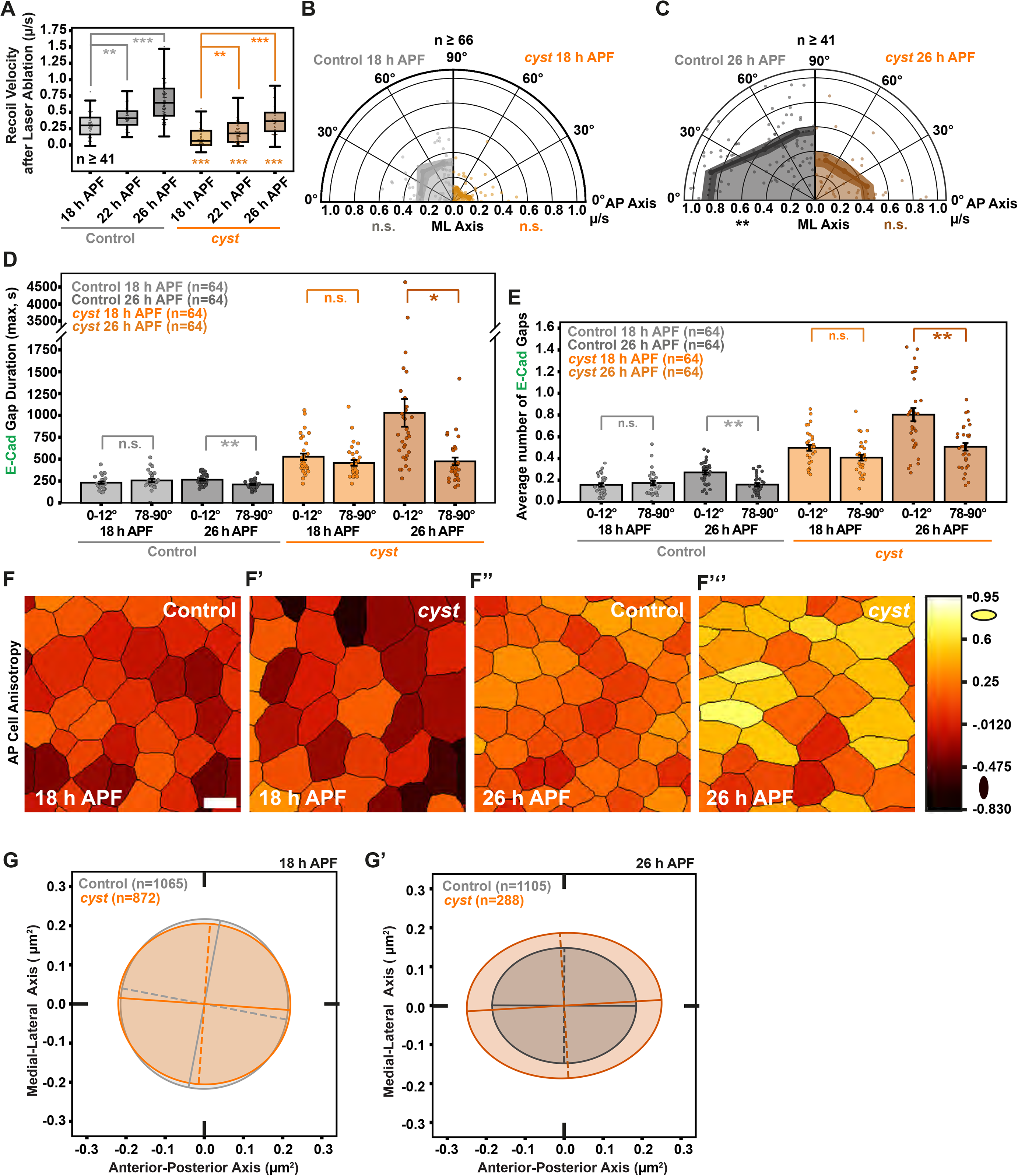
Epithelial mechano-response upon increase in endogenous tissue anisotropic stress. **(A)** Recoil velocity after laser ablation in control and *cyst* junctions at 18, 22, and 26 h APF. Box plots show the median, interquartile range, and whiskers at 1.5 × IQR; individual junctions are overlaid as dots. Statistical significance was assessed using Welch’s t-test. n refers to minimum number of junctions analyzed across all experimental conditions. **(B,C)** Polar plot of recoil velocity after laser ablation (mean ± SEM) as a function of junction orientation relative to the AP and ML axes in control (left half) and *cyst* mutant (right half) cells at 18 h APF (B) and 26 h APF (C). Statistical significance was assessed using Welch’s t-test. n is the minimum number of junctions analyzed across all experimental conditions. **(D,E)** E-Cad gap duration (mean ± SEM, D) and number of E-Cad gaps per 2 h (mean ± SEM, E) for junctions oriented along the AP axis (0-12° relative to the AP axis) and along the ML axis (78-90° relative to the AP axis) in control and *cyst* at 18 and 26 h APF. Minimum number of animals analyzed for this quantification was 2. **(F-F’’’)** Cell anisotropy maps along the AP axis in control (F,F’’) and cyst (F’,F’’’) at 18 h APF (F,F’) and 26 h APF (F’’,F’’’). Color scale indicates cell anisotropy along the AP axis. **(G, G’)** Cell anisotropy ellipses derived from the time-averaged inertia tensors of control and cyst mutant cells at 18 and 26 h APF. Minimum number of animals analyzed for this quantification was 2. Scale bars: 5 µm (F). Asterisks indicate statistical significance: *p < 0.05, **p < 0.01, ***p < 0.001; ns, not significant.

## Discussion

Understanding how epithelial cells regulate their shape over short and long timescales is central to understanding the dynamics and robustness of epithelial tissues. This question has been made particularly topical by several emerging lines of evidence of extensive mechanical fluctuations and heterogeneities affecting developing tissues in a variety of multicellular systems ^11,31–37^. Although this suggests that active sensing and feedback mechanisms would be necessary to buffer these different sources of noise and ensure robust shape regulation, this hypothesis had remained largely untested. In this work, we have proposed a minimal model of junctional length regulation, based on core mechano-chemical components of epithelial junctions, resulting in emergent mechano-responses. This is qualitatively distinct from previous findings considering explicit mechano-sensing mechanisms at the molecular scale (i.e. assembly or disassembly rates dependent on local mechanical stresses ^40–49,68,82–92^). In the proposed mechanism, as long as negative regulators of contractility have slower turnover dynamics than positive ones, they will recover differentially to any passive dilution arising from junctional elongation, resulting in temporary overactivation of contractility. This provides a dynamical stabilization to junction length, and could be a mechanistic basis for past proposals of emergent tissue elasticity via active stabilization of tensions between neighboring junctions ^93^.We demonstrate the quantitative predictive power of this model for *Drosophila* junctional dynamics. Indeed, we identify a robust mechano-chemical cycle: upon stochastic fluctuations leading to junctional elongation, local gaps are observed, which locally dilute junctional components. Given that E-Cad has slow dynamics with large immobile fraction, its dilution is maintained over time, allowing for the local Cyst-dependent recruitment of Myo-II, which overshoots until it contracts the junction to its original value, which restores high local E-Cad concentration and stops the cycle. Our results also suggest that this emergent mechanism is well-suited to explain theoretical findings from vertex models at the tissue scale, which posit that cells must be able to precisely maintain their junction perimeters ^24,69,94^, although mechanisms for this had so far been elusive.

While the full molecular interactions that sense local E-Cad dilution remain to be dissected further, we provide strong evidence that it is directly linked to the apical–basal polarization machinery. Indeed, local dilution of E-Cad is sufficient to recruit the RhoGEF Cyst through a local decrease in aPKC activity, as indicated by the accumulation of Yrt, which is normally excluded from the apical junctional domain by direct aPKC phosphorylation^72,73^. Since the apical exclusion of the lateral polarity complex plays a conserved role in the establishment and maintenance of epithelial polarity ^52,72,73,95–99^, our work reveals that this emerging mechano-response is rooted in the core mechanisms that govern epithelial polarization. Given that aPKC and polarity proteins also play key roles in regulating contractility in other non-epithelial contexts, for instance the cytoskeletal flows regulating the symmetry breaking of the *C. elegans* zygote ^100–104^ we anticipate that our finding would have broad relevance in other cell types. Furthermore, as previous studies in epithelial cell culture showed that the Cysts orthologue p114RhoGEF is also involved in mechano-response at sites of E-cad accumulation upon acute mechanical stress^85^, analysing the contribution of p114RhoGEF, Par6/aPKC, Lulu1/2 and E-Cad dynamics *in vitro* might provide important insights into the mechanical control of junction length homeostasis. Last, in the early *Drosophila* embryo Cyst contributes to AJ integrity, cell-cell rearrangements and cell ingression ^71,105,106^; processes that are known to be regulated by the Par6/aPKC complex and are associated with the production of mechanical forces and the remodelling of the actomyosin cytoskeleton ^16,17,107^. One could therefore envision that the contribution of the Par6/aPKC complex to cytoskeleton and junction dynamics in developmental processes might also depend on the emerging Cyst-dependent mechanochemical feedback. Accordingly, recent evidence indicates that aPKC, Yrt and Cyst may contribute to the response to mechanical forces in the *Drosophila* follicular epithelium^108^. Altogether, our findings indicate that the antagonism between the apical and lateral polarity complexes ensures junction length homeostasis and mechano-responses.

Although we have concentrated here on the biochemical consequences of E-Cad dilution, recent work underlines that E-Cad dilution can also have emergent mechanical consequences, for instance due to the potential role of E-Cad in regulating local friction ^53,109^. During cytokinesis, we had previously shown that E-Cad dilution in neighbors of dividing cells results in Cyst-dependent and self-organized actomyosin flows due to locally decreased friction with the cortex ^53^. Notably, this friction-dependent mechanism can be theoretically unified with our current homeostatic mechanism by considering both biochemical and mechanical contributions of E-Cad in the model (Extended Data Fig. 10D,E and SI Theory). Upon formation of new cell-cell contacts in zebrafish, contact expansion also dilutes both E-Cad and the actomyosin cortex, resulting in flows that contribute to the formation of a contact rim ^110^. Interestingly, alternative forms of mechano-responses have started to emerge over recent years, where sensing arises as an emergent property at the cellular, rather than molecular scale. For instance, changes in cytoplasmic crowding in response to compressive forces can drive global change in translation and cell growth ^111,112^ and the nucleus has been proposed to function as a mechanical sensor of stresses acting on entire cells ^113–118^. Furthermore, active materials, such as the actomyosin cortex, have been theoretically shown to generate collective responses to mechanical forces, for instance due to global network reorganization or active instabilities ^119,120^. These observations could help to explain recent findings that not only the magnitude, but also the timescales of force application, are critical to elicit different types of mechano-responses ^17,87,121,122^. Collectively, our findings and other published works underline that changes in local actomyosin levels can arise due to three distinct processes: local recruitment due to molecular mechano-sensors, advection from cytoskeletal flows and passive dilution from junctional stretch. Integrating these various mechano-responses as well as nucleus mechanosensing will be an important future goal to achieve a comprehensive understanding of the biophysical principles driving tissue organization and dynamics.

## Supporting information

Theory Note

## Acknowledgments

We thank R. Basto, N. Brown, Y Hong, J. Knöblich, E. Morais-De-Sá, D. St Johnston, X. Wang, Bloomington Drosophila Stock Center, Transgenic RNAi Project at Harvard Medical School, Vienna Drosophila Resource Center, Kyoto Drosophila Stock Center, and Developmental Studies Hybridoma Bank for reagents; UMR3215/U934 imaging facility PICT-IBiSA@BDD for help with microscopy; Floris Bosveld for comments on the manuscript. This work was supported by Institut Curie, CNRS, INSERM, ERC Advanced Scaling-Sensitivity (101020243), ARC (SL220130607097), ANR (TiMecaDiv 20CE13000801), CANCERO-INCA (PLBIO2020/BELLAICHE), ANR Labex DEEP (11-LBX-0044, ANR-10-IDEX-0001-02), and FWF (International project I06360, to EH).

## Author contributions

Conceptualization: JGL, EH, YB

Methodology: JGL, YB

Software: JGL, URG

Formal analysis: JGL

Investigation: JGL, URG, IC, DP, PPE

Resources: PPE

Data curation: JGL, EH

Writing – original draft: YB, EH

Writing – review & editing: JGL, EH, YB, with inputs from all authors

Visualization: JGL, EH,

Supervision: EH, YB

Project administration, Funding acquisition: EH, YB

## Materials and Methods

### Fly stocks and genetics

A comprehensive list of the *Drosophila* melanogaster lines used throughout this study, together with their corresponding references, is provided in Extended Data Table 1. Loss- and gain-of-function experiments were performed using three complementary genetic approaches: the FLP/FRT recombination system, the Gal4/UAS expression system, and the temperature-sensitive Gal4/Gal80ts/UAS system ^123–126^. Somatic mitotic clones were induced by applying a 1-hour heat shock at 37°C during the second or third larval instar stages. Flip-out clones were induced by applying a 20-minute heat shock at 37°C during the second or third larval instar stages. To compare gene-function phenotypes at 18 h APF and 26 h APF, the time elapsed after clone induction (hACI) was kept constant between conditions in order to minimize potential effects arising from protein perdurance. Unless otherwise indicated, animals carrying UAS-RNAi constructs were maintained at 29°C from clone induction until imaging.

### Generation of the Yurt:GFPx3 and Yurt:mKate2x3 alleles

The Yurt:GFPx3 and Yurt:mKate2x3 alleles were both produced through CRISPR/Cas9-driven homologous recombination directly at the endogenous locus, taking advantage of the *vas-Cas9* line ^127^. Guide RNAs targeting the desired yurt genomic sites were inserted into the *pCFD5: U6:3-t::gRNA* backbone ^128^ using the following primers: 5’-GTTCGATTCCCGGCTGGTGCATAGAACATTTAGCTTATACGGTTTTAGAGCTAGAAATAGCAAGTTAAAAT-3’ and 5’-GCGGCCCGGGTTCGATTCCCGGCCGATGCATTAGCTTATACGCGGAGTAGGTTTTAGAGCTAGAAATAGCAAGTTAA-3’

The homology arms were inserted into recombination plasmids that contained a *hs-miniwhite* selection cassette flanked by *loxP* sites ^53,129^ and the 3xGFP- or 3xmKate2-tagging sequences (vector and map available upon request). Amplification of the two homology regions (HR1 and HR2) bracketing the CRISPR/Cas9 cut sites was carried out with the following primer pairs: (i) Yurt:3xGFP: HR1, 5’-ATTCAGTGCACCACTCACTCCG-3’ and 5’-CGAAAATCCGTTATTACAACCCAGTTGAGTTCGGGGTCCAGCGGTTCTTCAGGCAGT-3’; HR2, 5’- TCGGTAGCTATGGAAATTTAGGCAG-3’ and 5’-CTAGGTCTTGAACGAAGACTGTTTTCAGTCAATCGAGTTCAAGGGCGACACAAAATT-3’; (ii) Yurt:3xmKate2: HR1, 5’-CCGGGCTAATTATGGGGTGTCGCCCTTCGTATTCAGTGCACCACTCACTCCG-3’ and 5’-CGAAAATCCGTTATTACAACCCAGTTGAGTTCGGGGTCCAGCGGTTCTTCAGGCAGT-3’; HR2, 5’- TCCGGAAGTGGTAGCTCAGGGTCTAGTGGATCGGTAGCTATGGAAATTTAGGCAG-3’ and 5’- CTAGGTCTTGAACGAAGACTGTTTTCAGTCAATCGAGTTCAAGGGCGACACAAAATT- 3’; All embryo injections for transgenesis were carried out by Bestgene.

### Live imaging microscopy

Pupal samples were mounted for live imaging of the dorsal thorax as previously described ^67^. Unless otherwise specified, acquisitions were performed at 25°C or 29°C using inverted spinning-disk confocal microscopes (Nikon or Zeiss) equipped with one of the following objectives: 60× NA 1.4 Oil DIC N2 Plan Apo VC (binning 1), 63× NA 1.4 Oil DICII Plan Apo (binning 1), or 100× NA 1.4 Oil DIC N2 Plan Apo VC (binning 2), coupled to a Hamamatsu sCMOS camera. In addition, time-lapse acquisitions were performed on a Zeiss LSM880 NLO platform equipped with a 63× NA 1.4 Oil DICII Plan Apo objective using 5× or 7.5× optical zoom for laser-ablation experiments (see Laser Ablation section), as this setup was coupled to a two-photon Ti laser source (Mai Tai DeepSee, Spectra Physics). Experiments typically consisted of 40-minute or 2-hour movies acquired at 20-second intervals from the anterior medio-lateral region of the notum (see Extended Data Figure 1A,A’) at 18 h APF using spinning-disk confocal microscopy. Alternatively, movies were acquired from a similar region at 26 h APF. E-Cad/Myo-II imaging experiments typically consisted of 13 z-steps (0.5 µm spacing) at 18 h APF or 23 z-steps at 26 h APF, when tissue curvature is more pronounced. For experiments involving weaker fluorescent signals, such as the ones associated with the acquisitions of Cyst, aPKC, Par6, Yrt or PH localizations, a 7–9 z-steps acquisition routine was used to minimize photobleaching.

### Optogenetics

E-Cad:GFPpupae expressing UAS-LARIAT using the Act-Gal4 driver in combination with Tub-Gal80ts were maintained at 25°C in complete darkness, and then animals were shifted to 29°C and mounted as previously described^130^, with the additional precaution that all manipulations were performed under red-light illumination^67,131^. Light-activation of LARIAT was triggered by exposure to a 488 or 491 nm laser, while E-Cad:GFP and Myo-II:mKate2x3 or Yrt:mKate2x3 dynamics were simultaneously followed using the 561 nm line. For analysis of junction dynamics upon clustering of E-Cad:GFP using LARIAT, LARIAT activation was performed by exposing the tissue to blue light for 1 to 2 min before the start of the time lapse movie and then re-exposed to blue light every 20 s to maintain E-Cad clustering. For the analysis of the dynamics of Myo-II:mKate2x3 or Yrt:mKate2x3 upon E-Cad clustering using LARIAT blue light exposure was done every 20 s during time-lapse acquisition.

### Laser ablations

To quantify the tissue recoil response following laser ablation, samples expressing E-Cad:GFP were imaged using a two-photon laser-scanning system (Carl Zeiss LSM780 or LSM880 NLO) equipped with a 63× NA 1.4 Oil DICII Plan Apo objective and 5× or 7.5× optical zoom. Image acquisition was performed using single-photon bidirectional scanning, with an acquisition time of t = 400 ms, repeated every 1 s for 12 timepoints: 2 timepoints before junction ablation and 10 timepoints after junction ablation. At t = 0 s, a rectangular region surrounding the target junction was defined, and junction ablation was performed using a Ti laser (Mai Tai DeepSee, Spectra Physics) tuned to 800 nm, delivering pulses shorter than 100 fs at an 80 MHz repetition rate, typically at 30% power. Following ablation, frames were collected every 1 s for 10 s. Recoil velocity was determined from the displacement of the two vertices connected by the ablated junction between the second and the fourth acquisitions. In addition, to estimate junction elasticity, the positions of all connected vertices involved in the local junctional network of the ablated junction were tracked over time, corresponding to the two vertices directly connected to the ablated junction and the two pairs of neighboring vertices linked to each vertex of the ablated junction. Junction orientations relative to the midline (i.e., bilateral symmetry axis of the dorsal thorax) were also measured to infer the anisotropy of mechanical stress exerted on junctions.

### FRAP experiments

FRAP assays were performed to assess the dynamics of E-Cad:GFP, Myo-II:GFPx3, Cyst:GFP, Crb:GFP, Yrt:GFPx3 tagged proteins. Bleaching was performed using an inverted spinning-disk confocal microscope equipped with a FRAP module (Roper/Nikon). The FRAP ROI covering approximately one-quarter to one-third of each junction was bleached using a 491 nm laser at 100% power for 40–50 iterations. For high-intensity fluorescently tagged proteins, such as E-Cad:GFP, Crb:GFP, and Myo-II:GFPx3, a single AJ plane was acquired at 1 s intervals after bleaching for at least 195 s. For fluorescently tagged proteins with lower signal-to-noise ratios, such as Yrt:GFPx3 or Cyst:GFP, images were acquired every 2 s or 5 s. Pre-bleaching fluorescence values were obtained by acquiring 4 to 6 timepoints before bleaching. When FRAP assays were performed in flip-out clones, the 405 nm channel was also acquired, typically every 6 to 12 timepoints, to track the position of RNAi- expressing cells based on CAAX:BFP2 labeling. Each FRAP movie was then corrected for photobleaching using histogram matching in Fiji/ImageJ. Intensities were measured over time in the FRAP ROI and normalized to a non-FRAP ROI junction.

### Image processing and segmentation

Movies were processed to generate two-channel junctional kymographs as follows: First, both imaging channels underwent signal pre-processing using custom Python scripts. Briefly, background subtraction was performed by subtracting the average projection background obtained from 3D image volumes acquired under identical imaging conditions, then denoised using a 3D Gaussian Blur filter [(x=1), (y=1), (z=0.5)], subsequently corrected for photobleaching using a histogram matching algorithm in Python. The resulting z-stack time-lapse images were then projected using an in-house MATLAB routine previously described ^81,132^. This pipeline allows us to project both channels within a 2 µm window centered around the AJ, defined by E-Cad signals. Projected stacks were then used for cell tracking, cell segmentation, and junction segmentation. Cell segmentations were generated using Cellpose^133^ with custom models trained on manually segmented control and *cyst* mutant tissues. Cell tracking was performed using previously described MATLAB pipelines^81^. Segmentation outputs were subsequently refined through iterative rounds of automatic and manual correction^81^. Junctional kymographs were generated by tracking individual junctions from the segmented and tracked cells. Signal integration along the junction was performed using a 3x3 weighted kernel: [(0.5,1, 0.5), (1,2,1), (0.5,1,0.5)] as junctions typically span approximately 3 pixels at this imaging resolution. Finally, projected time-lapse movies were converted to 8-bit images in Fiji using *Process > Enhance Contrast* followed by 8-bit conversion. For junctional signals, 0.05% saturated pixels were allowed, whereas approximately 0.3% saturated pixels were allowed for Myo-II and PH signals. As an additional output of the pipeline, CSV files containing the junction orientation at each time point were generated.

### E-Cad Gap Detection

E-Cad gaps (defined as local transient decreases in E-Cad signal intensity) were detected on 40-min or 2-h junction kymograph using custom FIJI/ImageJ macros and Python pipelines. Briefly, kymographs were binarized to generate masks restricting the analysis to the junctional region. E-Cad gaps were then detected on the inverted images using at least two alternative rounds of gap detection using different FindFoci Fiji plugin parameter combinations. Typical parameters included: background_parameter = 1, search_parameter = 0.15 or 0.375, peak_parameter = 0.04 or 0.1, and fraction_parameter = 0.4 or 0.775. Preliminary detections were refined through sequential morphological operations, including dilation, erosion, and opening, to reduce noise and regularize object boundaries. Candidate E-Cad gap masks were subsequently refined using local E-Cad intensity information extracted from the original kymographs. Threshold values of 0.6, 0.65, and 0.7 were typically used for control tissues. Since E-Cad signal values are lower in *cyst* mutant tissues, values of 0.7, 0.75, and 0.8 were used. Finally, E-Cad gap masks were filtered according to length (minimum of 0.490 µm) and temporal duration (minimum of 60 s). E-Cad gaps detected at the beginning or end of the kymograph time axis were excluded. Due to the high variability of E-Cad gap morphologies, particularly in *cyst* mutant tissues, as well as possible differences between E-Cad:mKate2x3 and E-Cad:GFP dynamics, candidate E-Cad gap masks were visually inspected using the E-Cad kymographs. This manual validation was performed blindly with respect to the non-E-Cad channel.

### Analysis of protein dynamics during gap formation using kymograph

Protein dynamics during E-Cad gap were analyzed using two-channel kymographs. For each movie, background subtraction was performed at each time point using apical-medial regions away from cell junctions. For each detected E-Cad gap, the selected ROI was extended before and after the event to include protein dynamics prior to and after the E-Cad gap formation. All events were aligned to the initial detection time point of the E-Cad gap. For each junction (*j*), channel (*c*), and time point (*t*), signal intensity inside the E-Cad gap region was normalized to the mean signal outside this region within the kymograph: 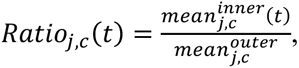 where *mean^inner^j,c*(*t*) is the mean signal inside the E-Cad gap ROI at time *t*, and *mean*^outer,global^*j*, *c* is the global mean signal outside the E-Cad gap ROI in the kymograph. To avoid edge or saturation artifacts, the first signal-containing pixel on each side of the kymograph, typically corresponding to tricellular junctions, was excluded from the analysis. *Ratio*_j,c_(*t*) were temporally rescaled to the mean E-Cad gap duration of each experimental condition using linear interpolation. E-Cad gap analyses typically included at least 50 E-Cad gaps from at least three animals. For display, the baseline of each signal was defined from the median between 300 and 200 s before the initial E-Cad gap detection: 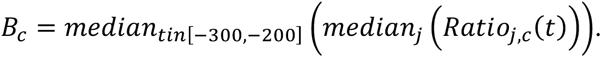 The displayed normalized signal was then calculated as: 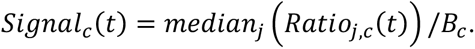 Dispersion was shown as the standard error of the mean (SEM) across junctions: 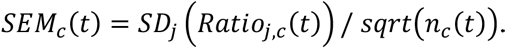 Local junctional deformation was estimated from changes in junction length rescaled by the initial relative size of the E-Cad gap within the junction: 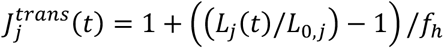 where 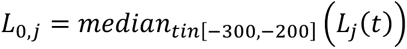 and 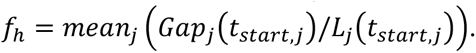 Here, *L_j_*(*t*) is junction length, *L*_0,*j*_ is the median of junction length between 300 and 200 s before the initial E-Cad gap detection, *Gap_j_*(*t_start,j_*) is the initial E-Cad gap size, and *f_h_*is the mean gap-to-junction length ratio at the time of gap detection. Statistical analyses were performed using sliding windows of three time points. For each junction and sliding window, ratio values were normalized to the baseline of each junction and were compared against the corresponding baseline values using paired Wilcoxon tests.

### Auto- and Cross-correlation analysis

To calculate the auto- and cross-correlation functions, we used our kymographs of junctional dynamics, and calculated, for each spatial position, how a given signal (e.g. E-Cad, Myo-II etc.) is correlated either with itself (auto-correlation) or with other signals (cross-correlation), with a delay interval dt. For details of the formula used and normalization methods, see the SI Theory Note.

### Mean Squared Displacement (MSD) Analysis of Junction Length

Junction length dynamics were extracted from kymographs by measuring junction length by binarizing non-zero signal pixels at each time point. For each kymograph, mean squared displacement (*MSD*) was calculated as the average squared change in junction length over increasing time intervals: *MSD*(*τ*) = mean_*t*_([*s*(*t* + *τ*) − *s*(*t*)]^6^), where *s*(*t*) is the junction length signal at time *t*, *τ* is the time interval, and the mean is taken over all valid time-point pairs separated by *τ*. MSD curves were computed up to 331 timepoints, corresponding to approximately 6620 s. For each animal, individual kymograph MSD curves were extracted at each time interval using a robust MAD-based z-score threshold of 10. This stringent filter was used to remove extreme MSD values likely caused by segmentation or tracking errors. For each experimental condition, the mean and SEM of the MSD were calculated as the mean and SEM of MSD extracted from each animal. When MSD curves were displayed as values normalized relative to a control condition, each experimental group MSD was divided by the corresponding control MSD curve at the same time interval. The uncertainty of the normalized MSD ratio was estimated by standard error propagation from each experimental condition.

### MSD of the Cell Shape index

Cell shape index was calculated only for cells that were reliably tracked throughout the movie and did not divide and touch the image border. Cell shape index was calculated from the polygon defined by the positions of the cell tricellular junctions (TCJs): 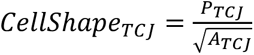 where *P_TCJ_*and *A_TCJ_* correspond to the perimeter and area, respectively, of the TCJ-defined polygon.

### T1 rate measurements in experimental data

To determine the rate of T1 events, the number of T1 were manually determined on the 2-h kymograph dataset. T1 events were defined as junctions that progressively shortened until disappearance. Junction disappearance associated with cell division or junction tracking errors were excluded from the analysis. See Theory SI for T1 measurements in Vertex simulations.

### Cell Anisotropy measurements

Cell shape anisotropy was estimated from the corrected Cartesian inertia tensor of each cell. For each cell, the three tensor components were first averaged over time: 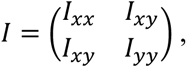 where *I*_{*xx*}_, *I*_{*xy*}_, and *I*_{*AA*}_ are the time-averaged corrected Cartesian inertia tensor components. The tensor was diagonalized to obtain its major and minor eigenvalues, *λ_major_* and *λ_minor_* and the corresponding eigenvectors. For visualization, each tensor was represented as an ellipse whose radii were defined from the square root of the eigenvalues: 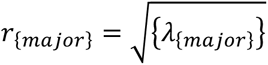 and 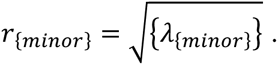 The ellipse orientation was given by the eigenvector associated with *λ_major_*. Since tensor components are in *μm*^4^, ellipse radii are measured in µm^2^.

### Statistics and Data Visualization

Sample sizes differed across experiments. Each experiment was repeated independently at least three times, except for the data presented in Fig. 3C and Fig. 6 for which two replicates were performed. The numbers of junction, ablations, animals, or cells included in each analysis are reported in the corresponding figures or figure legends. Unless otherwise indicated, error bars represent the SEM, and the statistical tests applied are specified in each figure legend. Statistical analyses were selected based on the type of comparison performed. Welch’s t-test, with Holm adjustment for multiple comparisons when appropriate, was used for comparisons between independent groups. Wilcoxon signed-rank tests were used for paired comparisons of temporal signal evolution against baseline. For these analyses, signals were normalized to the median baseline value to reduce sensitivity to noise. Statistical significance is represented by asterisks: *p < 0.05, **p < 0.01, ***p < 0.001; ns, not significant. All statistical computations were performed using the Python packages *scipy.stats*, *scikit_posthocs* and *netCDF4*. Simulation trajectories stored in NetCDF format were processed with custom scripts to extract junction connectivity graphs, compute FFT-based MSD curves, and aggregate metrics across ensembles. Visualization was performed using previously referred custom Python scripts based on NumPy, SciPy, Pandas, Matplotlib and Seaborn, and final figures were exported as .eps files and then assembled in Adobe Illustrator.

## Code availability

The MATLAB pipelines used cell tracking have been described in earlier publications ^81,134^. The MATLAB scripts, the Python Jupyter notebooks and scripts used for data analysis, plotting and statistics, custom Cellpose models^133^ and the Fiji macros developed for quantification as well as processed data will be made available upon publication.

## Use of AI tools

The text of the manuscript was proofread using Claude and ChatGPT. Codes for analysis were written with the help of Codex. All texts and codes were then manually checked.

---

**Extended Data Table 1.** Stocks used in this study.

| **Drosophila Stock** | **Promoter** | **Source** |
| --- | --- | --- |
| FRT40A, nls:GFP | Ubiquitin | Bloomington Drosophila Stock Center (5629) |
| Act-FRT-CD2-FRT-Gal4 | *Actin5C* | Bloomington Stock Center (3953) |
| Act-FRT-y+-FRT-Gal4 | *Actin5C* | Bloomington Stock Center (4780) |
| Jub:GFP | Endogenous | Bloomington Stock Center (56806) |
| E-Cad:GFP | Knock-in | Huang et al., 2009 <sup>135</sup> |
| Vinc:GFP | Endogenous | Klapholz et al, 2015 <sup>136</sup> |
| E-Cad:mKate2x3 | Knock-in | Pinheiro et al., 2017 <sup>53</sup> |
| Myo-II:GFPx3 | Endogenous | Pinheiro et al., 2017 <sup>53</sup> |
| Myo-II:mKate2x3 | Endogenous | Pinheiro et al., 2017 <sup>53</sup> |
| PH:GFP | αTub84B | Bloomington Stock Center (8164) |
| Crb:GFP | Knock-in | Huang et al., 2009 <sup>135</sup> |
| Cyst:GFP | Endogenous | di Pietro et al., 2023 <sup>54</sup> |
| Par6:GFP | Endogenous | Wirtz-Peitz et al., 2008 <sup>137</sup> |
| aPKC:GFP | Endogenous | Chen et al., 2018 <sup>138</sup> |
| Yrt:GFP | Endogenous | Vienna Drosophila Resource Center ( 318067) |
| Yrt:mKate2x3 | Knock-in | This paper |
| Yrt:GFPx3 | Knock-in | This paper |
| aPKC<sup>as4</sup>:mScarlet | Knock-in | Osswald et al, 2022 <sup>80</sup> |
| *vinc*<sup>D3</sup> | - | Pinheiro et al., 2017 <sup>53</sup> |
| *cyst*<sup>ARMCE</sup> | - | di Pietro et al., 2023 <sup>54</sup> |
| *rock*<sup>RNAi</sup> | UAS | Bloomington Stock Center (28797) |
| *a-spec*<sup>RNAi</sup> | UAS | Bloomington Stock Center (31209) |
| *aPKC*<sup>RNAi</sup> (Chr II) | UAS | Bloomington Stock Center (67204) |
| *cyst*<sup>RNAi</sup> (Chr II) | UAS | Bloomington Stock Center (38942) |
| *cyst*<sup>RNAi</sup> (Chr III) | UAS | Bloomington Stock Center (33047) |
| *yrt*<sup>RNAi</sup> | UAS | Bloomington Stock Center (36118) |
| *jub*<sup>RNAi</sup> | UAS | Bloomington Stock Center (41938) |
| *w*<sup>RNAi</sup> | UAS | Bloomington Stock Center (33644) |
| *e-cad*<sup>RNAi</sup> | UAS | Vienna Drosophila Resource Center (v103962) |
| *crb*<sup>RNAi</sup> | UAS | Bloomington Stock Center (27697) |
| CAAX:BFP2 (Chr II) | UAS | Bloomington Stock Center (from 55082 by H.S.) |
| CAAX:BFP2 (Chr III) | UAS | Bloomington Stock Center (from 55090 by H.S.) |
| LARIAT (Chr II) | UAS | Park et al, 2014 <sup>139</sup> |
| LARIAT (Chr III) | UAS | Qin et al, 2017 <sup>140</sup> |
| hs:FLP | Hsp70 | Bloomington Stock Center |

**Extended Data Figure 1.**
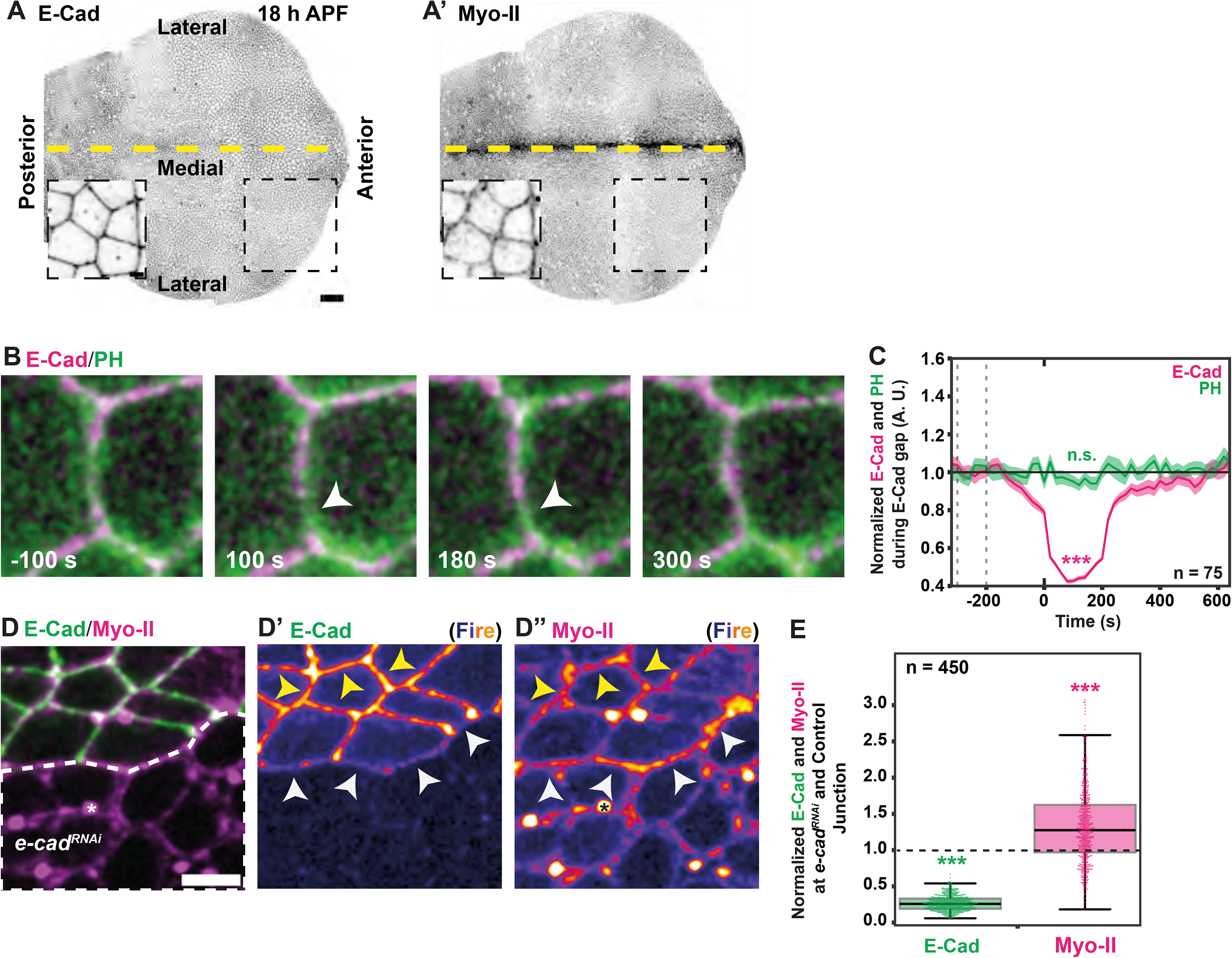
Junction dynamics and regulation of Myo-II level by E-Cad. **(A)** Pupal dorsal thorax images of E-Cad:GFP and Myo-II:mKate2x3 at 18 h APF as well as close-up. Dotted boxes indicate the anterior tissue region where the analyses were performed. Yellow dashed line: midline. **(B)** Time-lapse images of E-Cad:mKate2x3 and PH:GFP. White arrowheads mark the position of E-Cad gap and the absence of PH signal decrease. Timestamps are aligned to E-Cad gap onset. **(C)** Normalized E-Cad:mKate2x3 and PH:GFP signals (median ± SEM) aligned to local E-Cad decrease events (t = 0). E-Cad and PH intensities were normalized to their median baseline values between -300 and -200 s, indicated by dotted lines. Statistical significance was assessed using paired Wilcoxon signed-rank tests against baseline. n: total E-Cad gaps. Minimum number of animals analyzed for this quantification was 3. **(D)** E-Cad:GFP and Myo-II:mKate2x3 in control and *e-cad^RNAi^*cells. *e-cad^RNAi^* cells were identified by the expression of CAAX:BFP2 (not shown). Dotted line marks the position of the *e-cad^RNAi^* cells. E-Cad (D’) and Myo-II (D’’) are displayed with Fire LUT. To quantify Myo-II:mKate2x3 and E-Cad:GFP levels upon reduction of E-Cad function without confounding effects due to cortex detachment, Myo-II and E-Cad levels were measured at junctions between control and *e-cad^RNAi^*cells (white arrowheads) relative to control cell junction (yellow arrowheads). Asterisk indicates the position of the Myo-II labelled midbody. **(E)** Quantification of E-Cad:GFP and Myo-II:mKate2x3 levels at junctions between *e-cad^RNAi^* and control cells, normalized to neighboring control junctions. Box plots show the median, interquartile range (IQR), and whiskers at 1.5 × IQR; individual junctions are overlaid as dots. Statistical significance was assessed using Welch’s t-test. Number of animals analyzed for this quantification was 3. Scale bars: 50 µm (A); 2 µm (A close-up,B,D). Time is shown in seconds (s). Unless otherwise indicated, n and N indicate the number of junctions and animals, respectively. Asterisks indicate statistical significance: *p < 0.05, **p < 0.01, ***p < 0.001; ns, not significant.

**Extended Data Figure 2.**
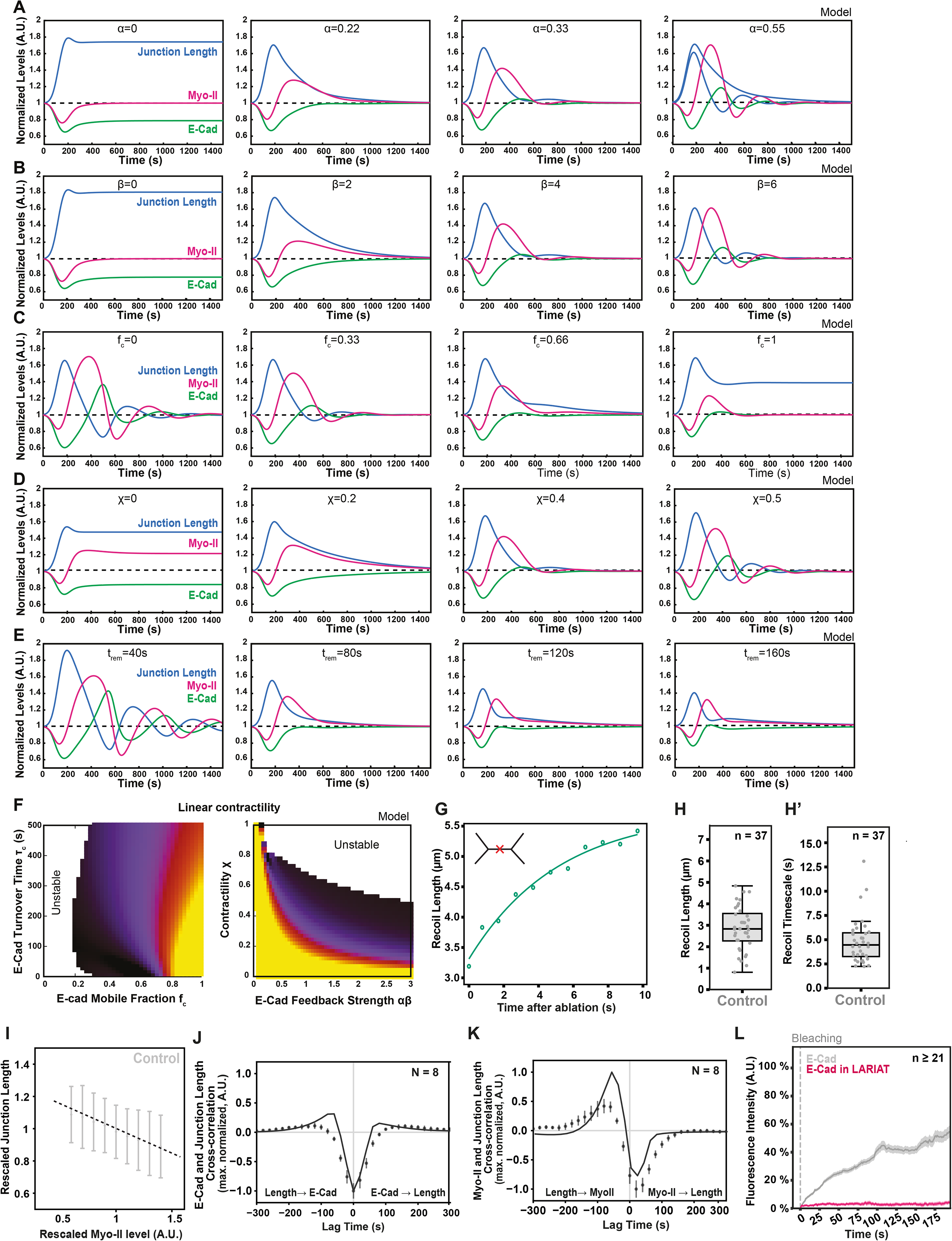
Sensitivity analysis of the model for predicting temporal responses to an E-Cad gap. **(A-E)** Temporal profiles of junctional length (blue), E-Cad level (green), and Myo-II level (magenta) in response to an E-Cad gap (see SI Theory); all homeostatic values are set to 1. For the sake of compactness, we show here already the full theory considering Cyst, although we found, as discussed in the theory, that the predictions are very close to the simplified model, considering the global feedback strength *γ* = *αβ*. (A) Response for increasing values of the Cyst to Myo-II coupling strength *α* (left to right), showing a lack of homeostatic response for low values of *α*, while large values of *α* lead to oscillations. (B) Response for increasing values of the E-Cad to Cyst coupling strength *β* (left to right), showing a lack of homeostatic response for low values of *β*, while large values of *β* lead to oscillations. (C) Response for increasing values of the E-Cad mobile fraction *fc* (left to right), showing oscillations for low mobile fractions and lack of homeostatic response for low immobile fractions. (D) Response for increasing values of Myo-II contractility *χ* (left to right), showing a lack of homeostatic response for low values of *χ*, while large values of *χ* lead to oscillations. (E) Response for increasing values of the timescale of junctional remodelling, *τ*_j_ (left to right), showing that very small values of *τ*_j_ lead to oscillations, while predictions are largely unaffected by values of *τ*_j_ above a given threshold. **(F)** Phase diagram for the amount of junctional length recovery, defined as junctional length 1000 s after the E-Cad gap. Large values (yellow) indicate poor junctional homeostasis. Recovery is shown as a function of model parameters in either linear or non-linear contractility models (see SI Theory for details). Unstable oscillatory junctions are indicated in white. This shows that, for most model parameters, there is a trade-off between the quality of junctional length homeostasis and the absence of oscillatory instability. **(G)** Representative scheme of how laser ablation experiments were analyzed to extract the timescale of short-term junctional relaxation. Junction length increases upon ablation, indicative of active tension within the junction, and recoils until it reaches a plateau. An experimental curve (dots) and the corresponding model fit (line) are shown in (G). **(H-H’)** Recoil length (H) and recoil timescale (H’) in control junctions, analyzed using the method explained in (G) (see SI Theory for details). Box plots show the median, interquartile range, and whiskers at 1.5 × IQR; individual junctions are overlaid as dots. n: the number of junction laser ablations analyzed. (**I)** Rescaled junction length as a function of rescaled Myo-II levels on the junction in control, showing a negative correlation from which we can infer the contractility parameter **χ** (see SI Theory for details). **(J,K)** Comparison of model predictions (solid lines) and experimental data (dots) for normalized cross-correlations between E-Cad and junction length (J), and Myo-II and junction length (K). Cross-correlation values were normalized to the maximum cross-correlation value. (L) FRAP recovery curves of E-Cad:GFP upon E-Cad:GFP clustering using LARIAT are shown as percentage of pre-bleach intensity (mean ± SEM). Time 0 corresponds to bleaching. n indicates the lowest number of FRAP measurements among the conditions shown. Time is shown in seconds (s). Unless otherwise indicated, n and N indicate the number of junctions and animals, respectively.

**Extended Data Figure 3.**
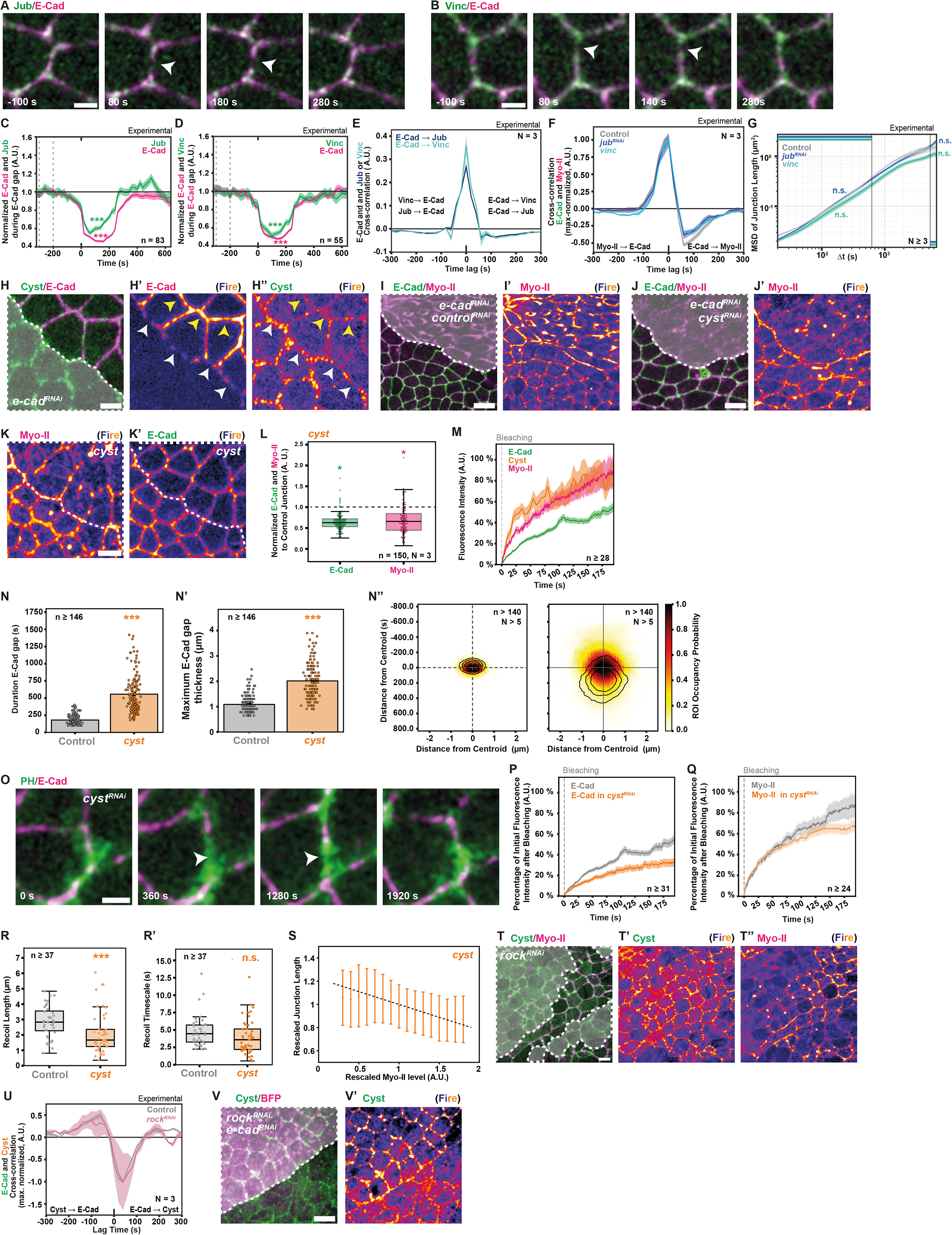
The RhoGEF Cyst controls E-Cad, Myo-II, and junction length dynamics. **(A,B)** Time-lapse images of Jub:GFP and E-Cad:mKate2x3 (A), and Vinc:GFP and E-Cad:mKate2x3 (B), in control tissues. White arrowheads mark E-Cad gap. Timestamps are aligned to E-Cad gap onset. **(C,D)** Normalized E-Cad:mKate2x3 and Jub:GFP (C), as well as E-Cad:mKate2x3 and Vinc:GFP (D), signals aligned to local E-Cad gap events (t = 0 s; median ± SEM). E-Cad and Jub (C), or E-Cad and Vinc (D), intensities were normalized to their median baseline values between -300 and -200 s, indicated by dotted lines. Statistical significance was assessed using paired Wilcoxon signed-rank tests against baseline. n: total E-Cad gaps. Minimum number of animals analyzed for this quantification was 3. **(E)** Cross-correlation between E-Cad:mKate2x3 and Jub:GFP and between E-Cad:mKate2x3 and Vinc:GFP intensities (mean ± SEM) in control junctions. Minimum number of junctions analyzed per animal was 161 junctions. **(F)** Normalized cross-correlation between E-Cad:GFP and Myo-II:mKate2x3 intensities (mean ± SEM) in control, *vinc* mutant, and *jub^RNAi^* junctions. Cross-correlation values were normalized to the maximum cross-correlation value. Minimum number of junctions analyzed per animal was 161 junctions. **(G)** MSD of junction length (mean ± SEM) over time intervals in control, *Vinc* mutant, and *jub^RNAi^* junctions. Statistical significance was assessed by comparing the mean short-timescale MSD (first 10%; 0-640 s) and long-timescale MSD (last 20%; 3960-6620 s) using Welch’s t-test. Minimum number of junctions analyzed per animal was 187 junctions. **(H-H’’)** Cyst:GFP and E-Cad:mKate2x3 in tissue containing *e-cad^RNAi^* clones. Dotted line (H) marks the position of the *e-cad^RNAi^*cells identified by the expression of CAAX:BFP2 (not shown). E-Cad (H’) and Cyst (H’’) displayed with Fire LUT. White arrowheads mark the position of junctions between control and *e-cad^RNAi^* cells, while yellow arrowheads indicate control cell junctions. **(I,I’)** E-Cad:GFP and Myo-II:mKate2x3 in tissue containing *control^RNAi^ (w^RNAi^)*, *e-cad^RNAi^* clones. Dotted line (I) marks the position of the *control^RNAi^ (w^RNAi^)*, *e-cad^RNAi^* cells identified by the expression of CAAX:BFP2 (not shown). Myo-II displayed with Fire LUT (I’). **(J,J’)** E-Cad:GFP and Myo-II:mKate2x3 in tissue containing *cyst^RNAi^*, *e-cad^RNAi^* clones. Dotted line (J) marks the position of the *cyst^RNAi^*, *e-cad^RNAi^* cells identified by the expression of CAAX:BFP2 (not shown). Myo-II displayed with Fire LUT (J’). **(K,K’)** E-Cad:GFP and Myo-II:mKate2x3 in tissue containing *cyst* mutant clones. Dotted line (K) marks the position of *cyst mutant* cells identified by the lack of expression of nls:GFP (not shown). Myo-II (K) and E-Cad (K’) displayed with Fire LUT. **(L)** Quantification of E-Cad:GFP and Myo-II:mKate2x3 levels at junctions of *cyst* mutant cells, normalized to neighboring control junctions. Box plots show the median, interquartile range, and whiskers at 1.5 × IQR; individual junctions are overlaid as dots. Statistical significance was assessed using Welch’s t-test. Minimum number of animals analyzed for this quantification was 3. **(M)** FRAP recovery curves of E-Cad:GFP, Myo-II:GFPx3, and Cyst:GFP are shown as percentage of pre-bleach intensity (mean ± SEM). Time 0 corresponds to bleaching (vertical dashed line). n indicates the lowest number of FRAP measurements among the conditions shown. **(N-N’’)** Quantification of E-Cad gap duration (N) and length (N’) in control and *cyst* mutant junctions. Bars show mean ± SEM; individual E-Cad gaps are overlaid as dots. ROI occupancy probability heat maps of E-Cad gaps in control (left) and *cyst* mutant junctions (right), centered on the gap centroid, are shown in (N’’). n: the minimum number of E-Cad gaps analyzed per condition. Statistical significance was assessed using Welch’s t-test. Minimum number of animals analyzed for this quantification was 3. **(O)** Time-lapse images of PH:GFP and E-Cad:mKate2x3 in *cyst^RNAi^* clones. White arrowheads mark PH localization at E-Cad gap. Timestamps are aligned to E-Cad gap onset. **(P,Q)** FRAP recovery curves of E-Cad:GFP (P) and Myo-II:GFPx3 (Q) at control and *cyst^RNAi^*junctions are shown as percentage of pre-bleach intensity (mean ± SEM). Time 0 corresponds to bleaching (vertical dashed line). n is the lowest number of FRAP measurements among the conditions shown. **(R,R’)** Recoil junction length (R) and recoil junction timescale (R’) upon junction laser ablation for control and *cyst* mutant junctions. Box plots show the median, interquartile range, and whiskers at 1.5 × IQR; individual junctions are overlaid as dots. n is the minimum number of junctions analyzed across all experimental conditions. Statistical significance was assessed using Welch’s t-test. **(S)** Rescaled junction length as a function of rescaled Myo-II levels on the junction in *cyst* mutant junctions (see SI Theory). **(T-T’’)** Cyst:GFP and Myo-II:mKate2x3 in tissue containing *rock^RNAi^*clones. Dotted line (T) marks the position of *rock^RNAi^* cells identified by the expression of CAAX:BFP2 (not shown). Cyst (T’) and Myo-II (T’’) are displayed with Fire LUT. **(U)** Normalized cross-correlation between E-Cad:mKate2x3 and Cyst:GFP intensities (mean ± SEM) in control and *rock^RNAi^* junctions. Cross-correlation values were normalized to the maximum cross-correlation value. Minimum number of junctions analyzed per animal was 67 junctions. **(V,V’)** Cyst:GFP and CAAX:BFP2 in *rock^RNAi^*, *e-cad^RNAi^* clones (V). Dotted line (V) marks the position of *rock^RNAi^*, *e-cad^RNAi^* cells identified by the expression of CAAX:BFP2 (not shown). Cyst (V’) is displayed with Fire LUT. Scale bars: 2 µm (A,O); 5 µm (H,K,V); 10 µm (I,T). Time is shown in seconds (s). Unless otherwise indicated, n and N indicate the number of junctions and animals, respectively. Asterisks indicate statistical significance: *p < 0.05, **p < 0.01, ***p < 0.001; ns, not significant.

**Extended Data Figure 4.**
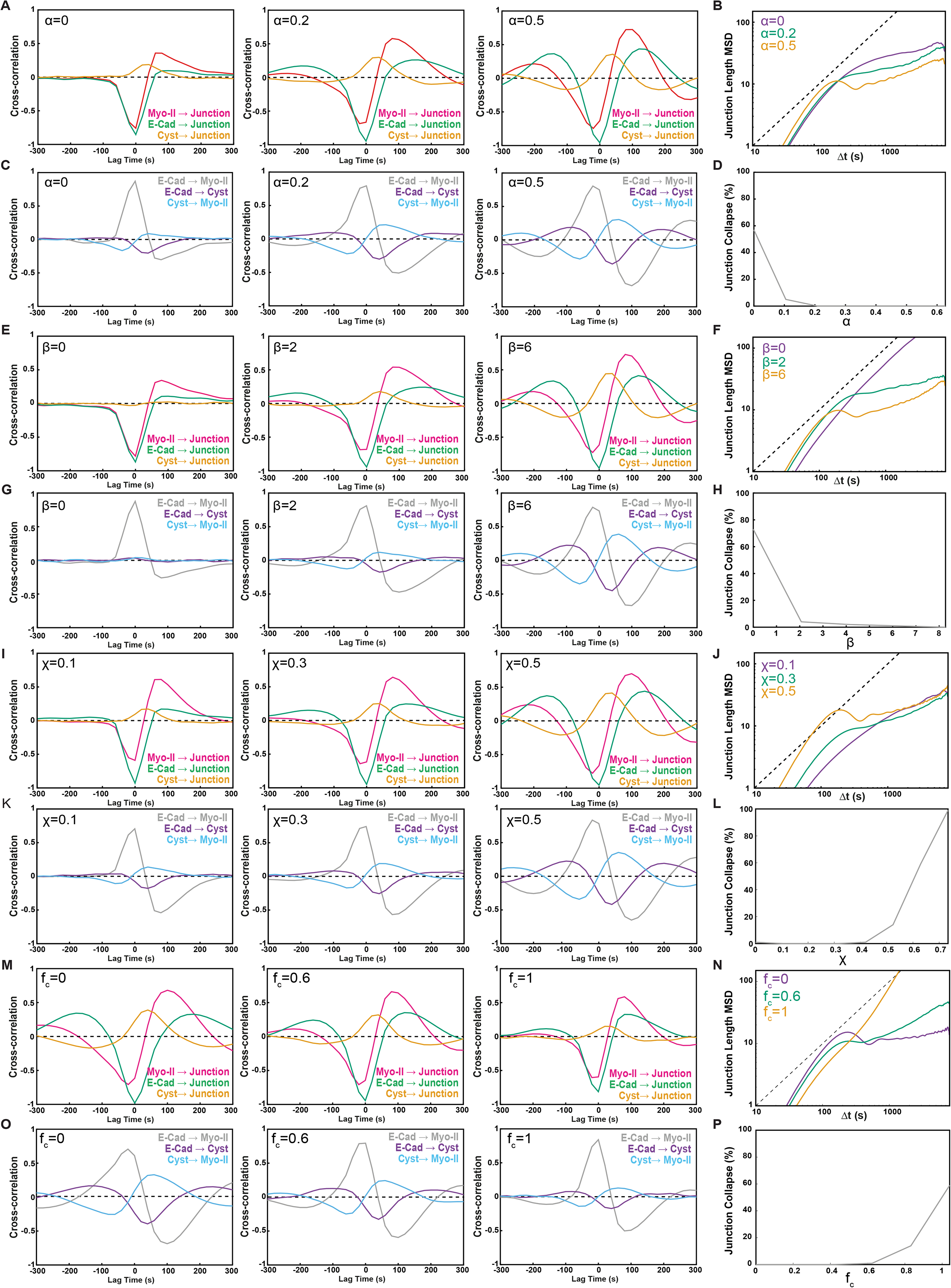
Sensitivity analysis of the model for cross-correlation and mean-square displacement predictions. In this figure, non-normalized cross-correlation functions are shown so that changes in the maximum correlation amplitude can be appreciated across parameter values. **(A-D)** Sensitivity of cross-correlation functions in the model to changes in the Cyst to Myo-II coupling strength *α*. Increasing values of *α* lead to increasing correlations between local concentrations (Myo-II, E-Cad, Cyst) and junctional length (A), as well as to reduced junction length fluctuations (B) and junction collapse or T1 (D). This also results in an increasingly negative anti-correlation from E-Cad to Myo-II (C). **(E-H)** Sensitivity of cross-correlation functions in the model to changes in the E-Cad to Cyst coupling strength *β*. Increasing values of *β* lead to increasing correlations between local concentrations (Myo-II, E-Cad, Cyst) and junctional length (E), as well as to reduced junction length fluctuations (F) and junction collapse or T1 (H). This also results in an increasingly negative anti-correlation from E-Cad to Myo-II (G). **(I-L)** Sensitivity of cross-correlation functions in the model to changes in the contractility *χ*. Increasing values of *χ* lead to increasing correlations between local concentrations (Myo-II, E-Cad, Cyst) and junctional length (I), as well as a complex effect on junction length fluctuations and junction collapse (J, L), as contractility is both required for the homeostatic response, but also drives fluctuations and Myo-II self-amplification (see SI Theory for details). **(M-P)** Sensitivity of cross-correlation functions in the model to changes in the E-Cad mobile fraction *f*_*c*_. Increasing values of *f*_*c*_ have relatively little impact on cross-correlations, except for rescaling the absolute magnitude of the correlations, because more immobile E-Cad is more sensitive to dilution effects (M, N). As discussed in the main text, increasing the mobile fraction both decreases short-term MSD and increases long-term MSD (N, P).

**Extended Data Figure 5.**
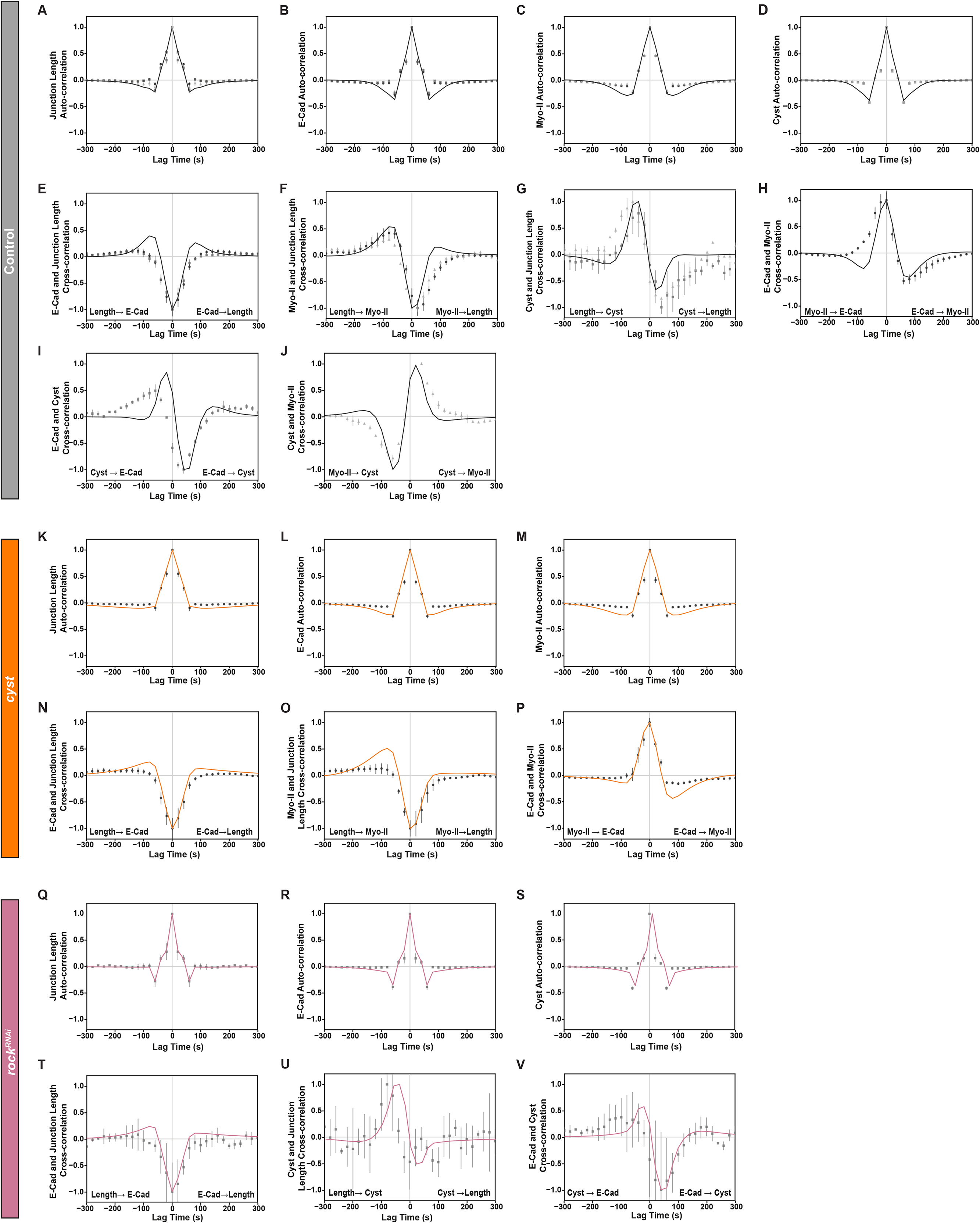
Data-model comparison for auto- and cross-correlation functions. Across this figure, experimental data are shown as dots, while model predictions are shown as solid lines; colors depend on the specific experimental condition or genotype. Some panels show two different datasets (two shades of grey), because they were extracted from two different experimental conditions. For instance, the cross-correlation of Cyst and junction length includes data from both E-Cad:mKate2x3/Cyst:GFP and Cyst:GFP/Myo-II:mKate2x3 conditions. All cross-correlation values were normalized to the maximum cross-correlation value. **(A-J)** Comparison between experimental data and the full model for control junctions. Auto-correlation functions are shown for junction length (A), E-Cad (B), Myo-II (C), and Cyst (D). Cross-correlation functions are shown for E-Cad and junction length (E), Myo-II and junction length (F), Cyst and junction length (G), E-Cad and Myo-II (H), E-Cad and Cyst (I), and Cyst and Myo-II (J). **(K-P)** Comparison between experimental data and the full model for *cyst* mutant junctions. Auto-correlation functions are shown for junction length (K), E-Cad (L), and Myo-II (M). Cross-correlation functions are shown for E-Cad and junction length (N), Myo-II and junction length (O), and E-Cad and Myo-II (P). **(Q-V)** Comparison between experimental data and the full model for *rock^RNAi^* junctions. Auto-correlation functions are shown for junction length (Q), E-Cad (R), and Cyst (S). Cross-correlation functions are shown for E-Cad and junction length (T), Cyst and junction length (U), and E-Cad and Cyst (V).

**Extended Data Figure 6.**
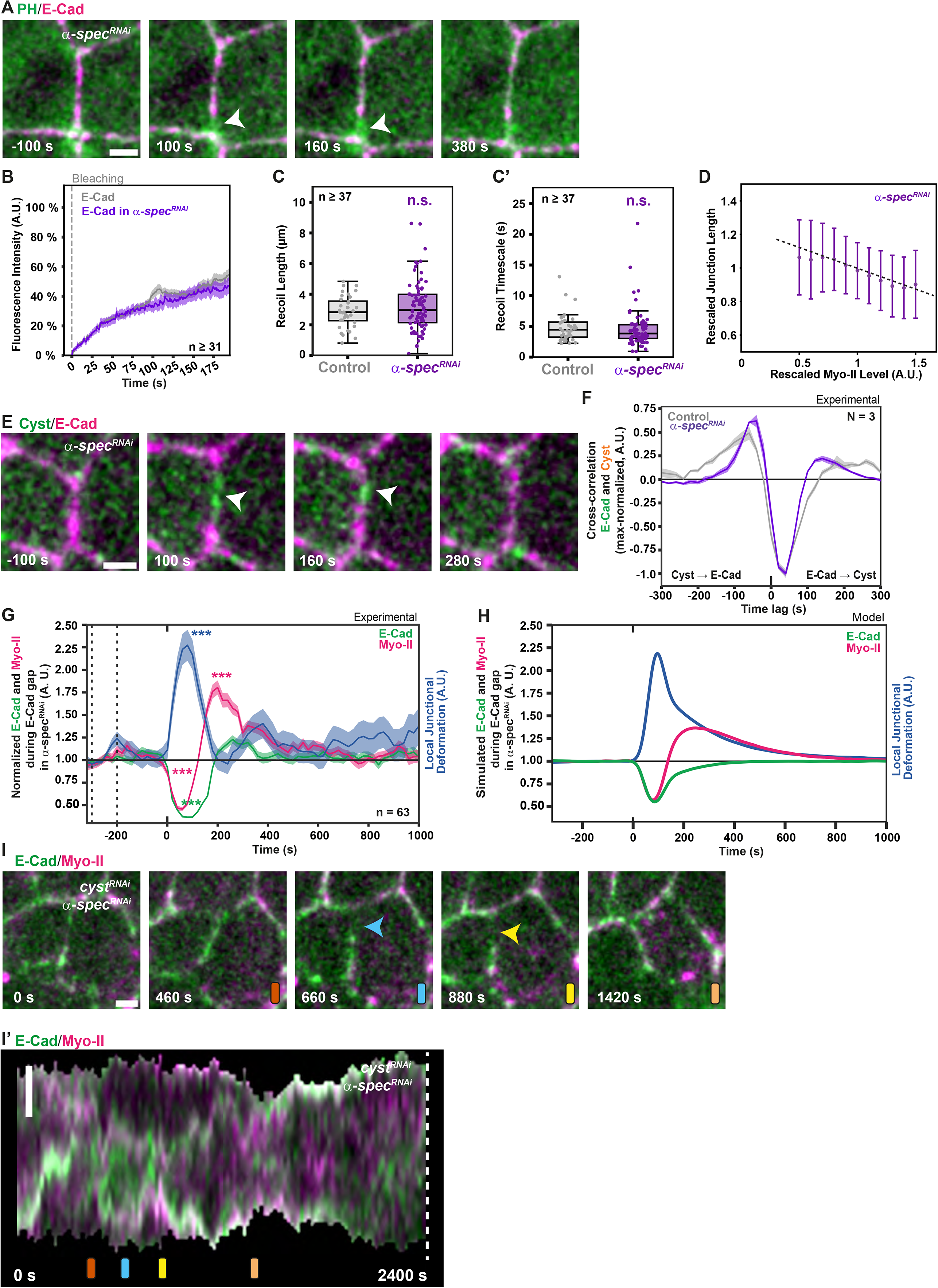
α-Spectrin controls short-term junction dynamics. **(A)** Time-lapse images of PH:GFP and E-Cad:mKate2x3 in an *α-spec^RNAi^*junction. White arrowheads mark the position of E-Cad gap and the absence of PH decrease. Timestamps are aligned to E-Cad gap onset. **(B)** FRAP recovery curves of E-Cad:GFP in control and *α-spec^RNAi^*junctions are shown as percentage of pre-bleach intensity (mean ± SEM). Time 0 corresponds to bleaching (vertical dashed line). n indicates the lowest number of FRAP measurements among the conditions shown. **(C-C’)** Recoil length (C) and recoil timescale (C’) upon junction laser ablation for control and *α-spec^RNAi^* junctions. Box plots show the median, interquartile range, and whiskers at 1.5 × IQR; individual junctions are overlaid as dots. n is the minimum number of junctions analyzed across all experimental conditions. Statistical significance was assessed using Welch’s t-test. **(D)** Rescaled junction length as a function of rescaled Myo-II levels on the junction in *α-spec^RNAi^*junctions (see SI Theory). **(E)** Time-lapse images of Cyst:GFP and E-Cad:mKate2x3 in an *α-spec^RNAi^* junction. White arrowheads mark the position of the E-Cad gap and Cyst upon gap formation and during closure. Timestamps are aligned to E-Cad gap onset. **(F)** Normalized cross-correlation between E-Cad and Myo-II intensities (mean ± SEM) in control and *α-spec^RNAi^*junctions. Cross-correlation values were normalized to the maximum cross-correlation value. Minimum number of junctions analyzed per animal was 161. Minimum number of animals analyzed for this quantification was 3. **(G)** Normalized E-Cad, Myo-II, and local junction deformation (median ± SEM) aligned to local E-Cad gap events (t = 0) in *α-spec^RNAi^* junctions. E-Cad and Myo-II intensities, as well as local junction deformation, were normalized to their median baseline values between -300 and -200 s, indicated by dotted lines. Statistical significance was assessed using paired Wilcoxon signed-rank tests against baseline. n: total E-Cad gaps. Minimum number of animals analyzed for this quantification was 6. **(H)** Simulated dynamics of E-Cad, Myo-II, and local junction deformation in *α-spec^RNAi^* junctions aligned to local E-Cad gap events (t = 0). **(I,I’)** Time-lapse images of E-Cad:GFP and Myo-II:mKate2x3 in a *cyst^RNAi^*, *α-spec^RNAi^*junction. Colored bars (orange, blue, yellow, peach) above each panel indicate the corresponding time points in the kymograph in (I’). Blue and yellow arrowheads mark the position of the E-Cad gap and the subsequent Myo-II accumulation. Kymograph of E-Cad:GFP and Myo-II:mKate2x3 along the junction shown in (I) over 2400 s (40 min) is shown in (I’). Scale bars: 2 µm (A, E, I-I’). Time is shown in seconds (s). Unless otherwise indicated, n and N indicate the number of junctions and animals, respectively. Asterisks indicate statistical significance: *p < 0.05, **p < 0.01, ***p < 0.001; ns, not significant.

**Extended Data Figure 7.**
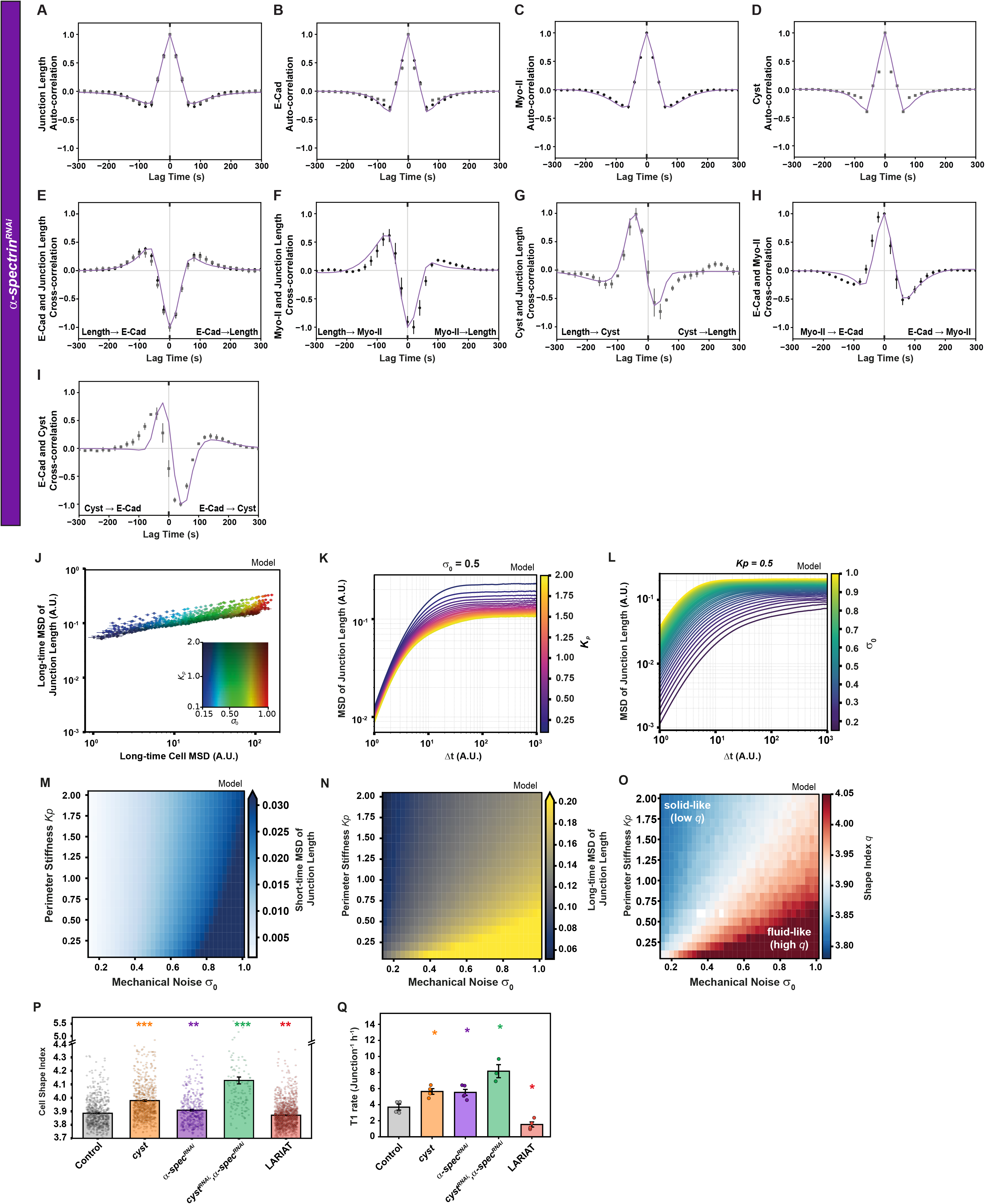
Modeling of the *α-spec^RNAi^* junction dynamics and Vertex simulation. From panels A to I, experimental data are shown as grey dots, while model predictions are shown in purple. Colors depend on the specific experimental condition (*α-spec^RNAi^*). All cross-correlation values were normalized to the maximum cross-correlation value. **(A-I)** Comparison between experimental data and the full model for *α-spec^RNAi^*cells. Auto-correlation functions are shown for junction length (A), E-Cad (B), Myo-II (C), and Cyst (D). Cross-correlation functions are shown for E-Cad and junction length (E), Myo-II and junction length (F), Cyst and junction length (G), E-Cad and Myo-II (H), and E-Cad and Cyst (I). **(J)** Long-time MSD of junction length as a function of long-time cell MSD in AVM simulations (log–log axes), illustrating the decoupling between cell motility (driven by *σ*_0_) and junction length fluctuations (controlled by *K*_p_ via the T1 rate). Color encodes (*K*_p_*, σ*_0_) as shown in the color key in Figure 4F. **(K)** MSD of junction length as a function of lag time Δ*t* (A.U) for all *K*_P_ values (0.1–2.0) at fixed *σ*_0_ = 0.5 (log–log axes). Color encodes *K_P_* using the plasma colormap (dark: low *K*_P_; bright: high *K*_P_). Curves overlap at short times, where dynamics are governed by *σ*_7_; at long times, higher *K*_P_ yields a lower plateau in junction length MSD. **(L)** MSD of junction length as a function of lag time Δ*t* (log–log axes) for all *σ*_0_ values (0.15–1.0) at fixed *K*_P_ = 0.5. Color encodes *σ*_0_ using the viridis colormap (dark: low *σ*_0_; yellow: high*Σ*_0_). Both short- and long-time junction length MSD increase with *σ*_0_, with the effect saturating at high *σ*_0_; all curves converge to a similar long-time plateau at fixed *K*_P_. **(M)** Phase diagram of short-time junction length MSD in (*σ*_0_, *K*_P_) parameter space. Short-time MSD increases monotonically with *σ*_0_ and is largely independent of *K*_P_, confirming that *σ*_0_ selectively drives short-time junction fluctuations. **(N)** Phase diagram of long-time junction length MSD in (*σ*_0_*, K*_P_) parameter space. Low *K*_P_ combined with high *σ*_0_ (lower right) yields the largest long-time MSD, consistent with a fluid, highly fluctuating regime; high *K*_P_ (top) suppresses long-time MSD regardless of *σ*_0_, in contrast to the *K*_P_ independent short-time behaviour in (M). **(O)** Phase diagram of shape index 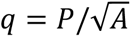 in (*σ*_0_*, K*_P_) parameter space (*σ*_0_: 0.15→1.0; *K*_P_: 0.1→2.0). Diverging colormap, *q* range 3.80–4.05, with the plot annotated directly with “solid-like (low *q*)” (top left) and “fluid-like (high *q*)” (bottom right). The solid–fluid boundary sweeps diagonally across parameter space: high *K*_P_ or low *σ*_0_ keeps the tissue solid-like; low *K*_P_ and high *σ*_0_ produce fluid-like behaviour. **(P)** Quantification of the cell shape index of control, *cyst* mutant, *α-spec^RNAi^*, *cyst^RNAi^ α-spec^RNAi^*, and LARIAT tissues. Bar plots show the mean ± SEM; individual cells are overlaid as dots. Statistical significance was assessed using Welch’s t-test with Holm adjustment, relative to control. Minimum number of animals analyzed for this quantification was 3. **(Q)** Quantification of the rate of T1 events per hour in control, *cyst* mutant, *α-spec^RNAi^*, *cyst^RNAi^ α-spec^RNAi^*, and LARIAT tissues. Bar plots show the mean ± SEM; individual animals are overlaid as dots. Statistical significance was assessed using Welch’s t-test with Holm adjustment, relative to control. Time is shown in seconds (s). Unless otherwise indicated, n and N indicate the number of junctions and animals, respectively.

**Extended Data Figure 8.**
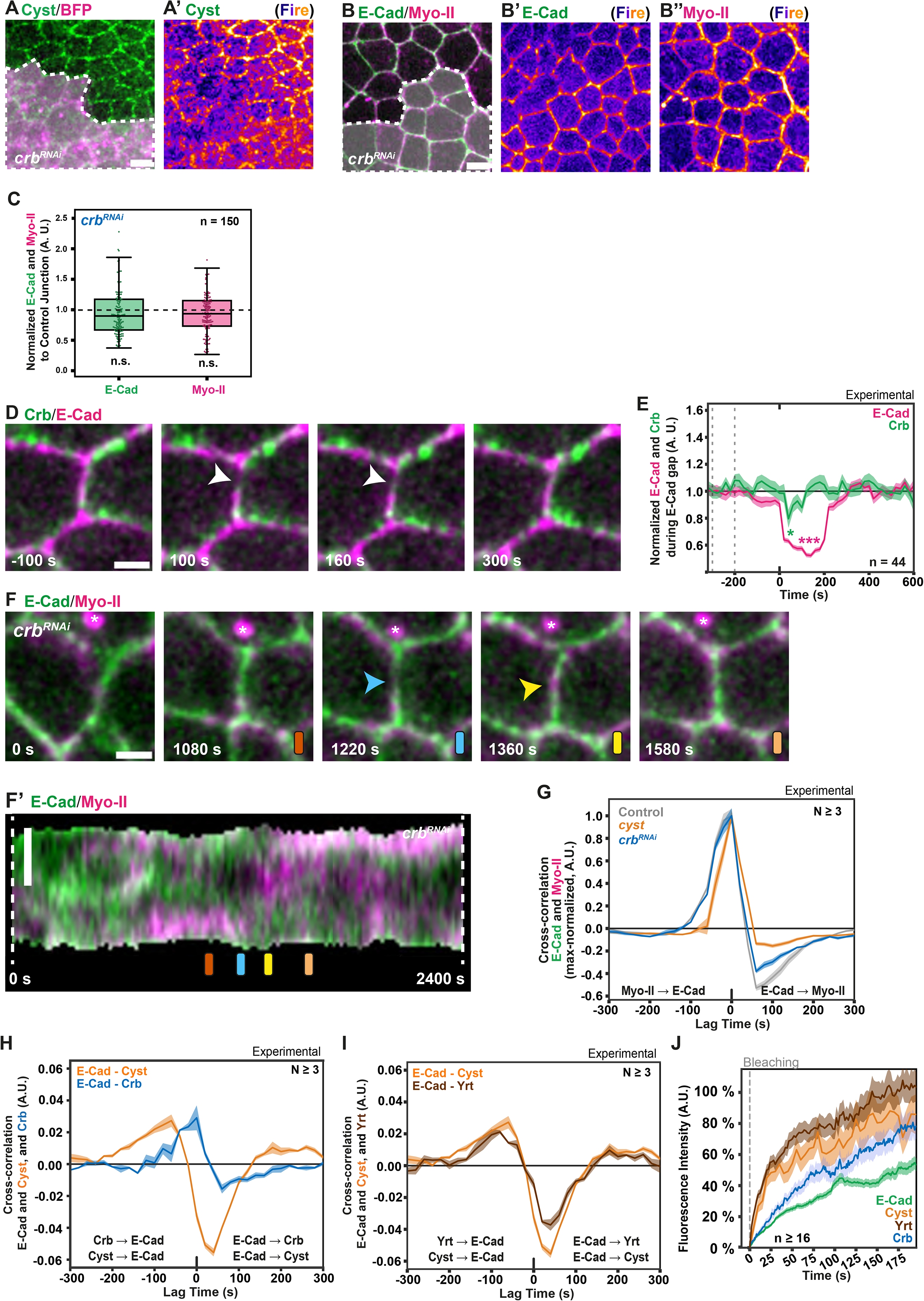
Analysis of Crb function in E-Cad gap regulation. **(A,A’)** Cyst:GFP and CAAX:BFP2 in tissue with *crb^RNAi^* cells. Dotted line (A) marks the position of *crb^RNAi^* cells identified by the expression of CAAX:BFP2 (not shown). Cyst:GFP displayed with Fire LUT (A’). **(B-B’’)** E-Cad:GFP and Myo-II:mKate2x3 in tissue with *crb^RNAi^*cells. Dotted line (B) marks the position of *crb^RNAi^* cells identified by the expression of CAAX:BFP2 (not shown). E-Cad:GFP (B’) and Myo-II:mKate2x3 (B’’) displayed with Fire LUT. **(C)** Quantification of E-Cad:GFP and Myo-II:mKate2x3 levels at junctions of *crb^RNAi^* cells, normalized to neighboring control junctions. Box plots show the median, interquartile range, and whiskers at 1.5 × IQR; individual junctions are overlaid as dots. Statistical significance was assessed using Welch’s t-test. Minimum number of animals analyzed for this quantification was 3. **(D)** Time-lapse images of Crb:GFP and E-Cad:mKate2x3 in control tissue. White arrowheads mark E-Cad gap and local decrease of Crb. Timestamps are aligned to E-Cad gap onset. **(E)** Normalized E-Cad:mKate2x3 and Crb:GFP intensities (median ± SEM) aligned to local E-Cad decrease events (t = 0). E-Cad and Crb intensities were normalized to their median baseline values between -300 and -200 s, indicated by dotted lines. Statistical significance was assessed using paired Wilcoxon signed-rank tests against baseline. n: total E-Cad gaps. Minimum number of animals analyzed for this quantification was 3. **(F,F’)** Time-lapse images of E-Cad:GFP and Myo-II:mKate2x3 at a *crb^RNAi^*junction (F). Colored bars (orange, blue, yellow, peach) above each panel indicate the corresponding time points in the kymograph in (F’). Asterisk indicates the position of the Myo-II labelled midbody. Blue and yellow arrowheads mark the formation of the E-Cad gap and the subsequent increase of Myo-II:mKate2x3. Kymograph of E-Cad:GFP and Myo-II:mKate2x3 along the junction shown in (F) over 2400 s (40 min, F’). Colored bars indicate the time points displayed in (F). **(G)** Normalized cross-correlation between E-Cad and Myo-II intensities (mean ± SEM) in control, *cyst* mutant, and *crb^RNAi^* junctions. Cross-correlation values were normalized to the maximum cross-correlation value. Minimum number of junctions analyzed per animal was 109 junctions. **(H)** Comparison of cross-correlations between E-Cad and Cyst intensities (orange) and E-Cad and Crb intensities (blue) in control junctions (mean ± SEM). Minimum number of junctions analyzed per animal was 120 junctions. **(I)** Comparison of cross-correlations between E-Cad and Cyst intensities (orange) and E-Cad and Yrt intensities (brown) in control junctions (mean ± SEM). Minimum number of junctions analyzed per animal was 75 junctions. **(J)** FRAP recovery curves of E-Cad:GFP, Myo-II:GFPx3, Crb:GFP, and Yrt:GFPx3 are shown as percentage of pre-bleach intensity (mean ± SEM). Time 0 corresponds to bleaching (vertical dashed line). n indicates the lowest number of FRAP measurements among the conditions shown. Scale bars: 10 µm (A,B); 2 µm (D,F,F’). Time is shown in seconds (s). Unless otherwise indicated, n and N indicate the number of junctions and animals, respectively. Asterisks indicate statistical significance: *p < 0.05, **p < 0.01, ***p < 0.001; ns, not significant.

**Extended Data Figure 9.**
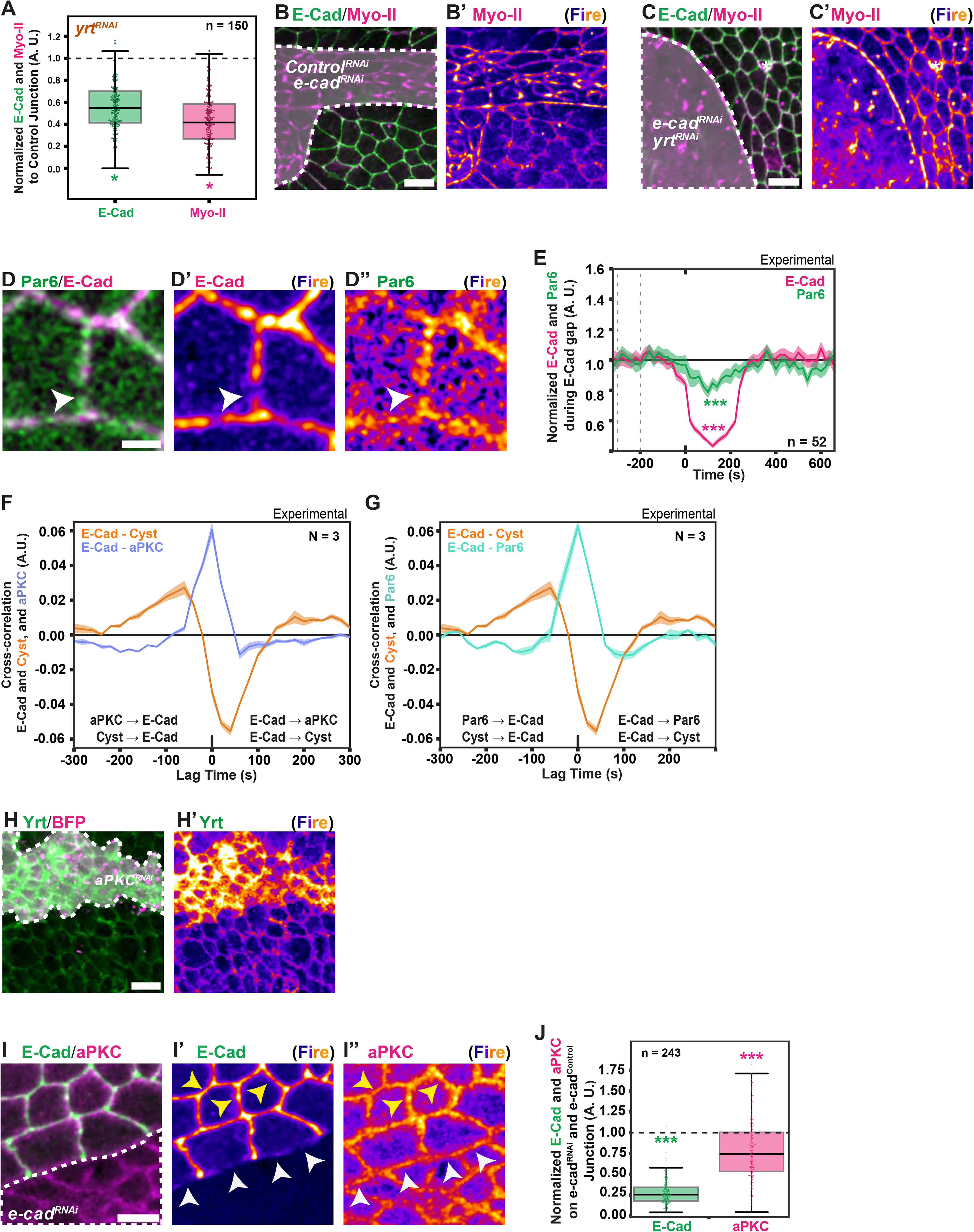
Analysis of Yrt and aPKC function in E-Cad gap regulation. **(A)** Quantification of E-Cad:GFP and Myo-II:mKate2x3 levels at junctions of *yrt^RNAi^* cells normalized to neighboring control junctions. Box plots show the median, interquartile range, and whiskers at 1.5 × IQR; individual junctions are overlaid as dots. Statistical significance was assessed using Welch’s t-test. Minimum number of animals analyzed for this quantification was 3. **(B,B’)** E-Cad:GFP and Myo-II:mKate2x3 in tissue with *control^RNAi^*(*w^RNAi^*), *e-cad^RNAi^* cells. Dotted line (B) marks the position of *control^RNAi^* (*w^RNAi^*), *e-cad^RNAi^* cells identified by the expression of CAAX:BFP2 (not shown). Myo-II (B’) displayed with Fire LUT. **(C,C’)** E-Cad:GFP and Myo-II:mKate2x3 in tissue with *yrt^RNAi^*, *e-cad^RNAi^* cells. Dotted line (C) marks the position of *yrt^RNAi^*, *e-cad^RNAi^* cells identified by the expression of CAAX:BFP2 (not shown). Myo-II (C’) displayed with Fire LUT. Asterisk indicates a microchaete. **(D-D’’)** Par6:GFP and E-Cad:mKate2x3 in control tissue (D). Par6:GFP (D’) and E-Cad:mKate2x3 (D’’) displayed with Fire LUT. White arrowhead marks E-Cad gap and the local decrease of Par6 signal. **(E)** Normalized E-Cad:mKate2x3 and Par6:GFP intensities (median ± SEM) aligned to local E-Cad decrease events (t = 0). E-Cad and Par6 intensities were normalized to their median baseline values between -300 and -200 s, indicated by dotted lines. Statistical significance was assessed using paired Wilcoxon signed-rank tests against baseline. n: total E-Cad gaps. Minimum number of animals analyzed for this quantification was 3. **(F)** Comparison of cross-correlations between E-Cad and Cyst intensities (orange) and E-Cad and aPKC intensities (violet) in control junctions (mean ± SEM). Minimum number of junctions analyzed per animal was 119 junctions. **(G)** Comparison of cross-correlations between E-Cad and Cyst intensities (orange) and E-Cad and Par6 intensities (cyan) in control junctions (mean ± SEM). Minimum number of junctions analyzed per animal was 113 junctions. **(H,H’)** Yrt:GFP and CAAX:BFP2 in tissue containing *aPKC^RNAi^*cells. Dotted line (H) marks the position of *aPKC^RNAi^* cells identified by the expression of CAAX:BFP2 (not shown). Yrt displayed with Fire LUT (H’). **(I-I’’)** E-Cad:GFP and aPKC:mScarlet in control and *e-cad^RNAi^*cells. *e-cad^RNAi^* cells were identified by the expression of CAAX:BFP2 (not shown). Dotted line marks the position of the *e-cad^RNAi^* cells. E-Cad (I’) and aPKC (I’’) displayed with Fire LUT. To quantify aPKC:mScarlet and E-Cad:GFP levels upon reduction of E-Cad function without confounding effects due to cortex detachment, aPKC:mScarlet and E-Cad levels were measured at junctions between control and *e-cad^RNAi^* cells (white arrowheads) relative to control cell junction (yellow arrowheads). **(J)** Quantification of E-Cad:GFP and aPKC:mScarlet levels at junctions between *e-cad^RNAi^* and control cells, normalized to neighboring control junctions. Box plots show the median, interquartile range, and whiskers at 1.5 × IQR; individual junctions are overlaid as dots. Statistical significance was assessed using Welch’s t-test. Minimum number of animals analyzed for this quantification was 3. Scale bars: 10 µm (B,C,H); 2 µm (D); 5 µm (I). Time is shown in seconds (s). Unless otherwise indicated, n and N indicate the number of junctions and animals, respectively. Asterisks indicate statistical significance: *p < 0.05, **p < 0.01, ***p < 0.001; ns, not significant.

**Extended Data Figure 10.**
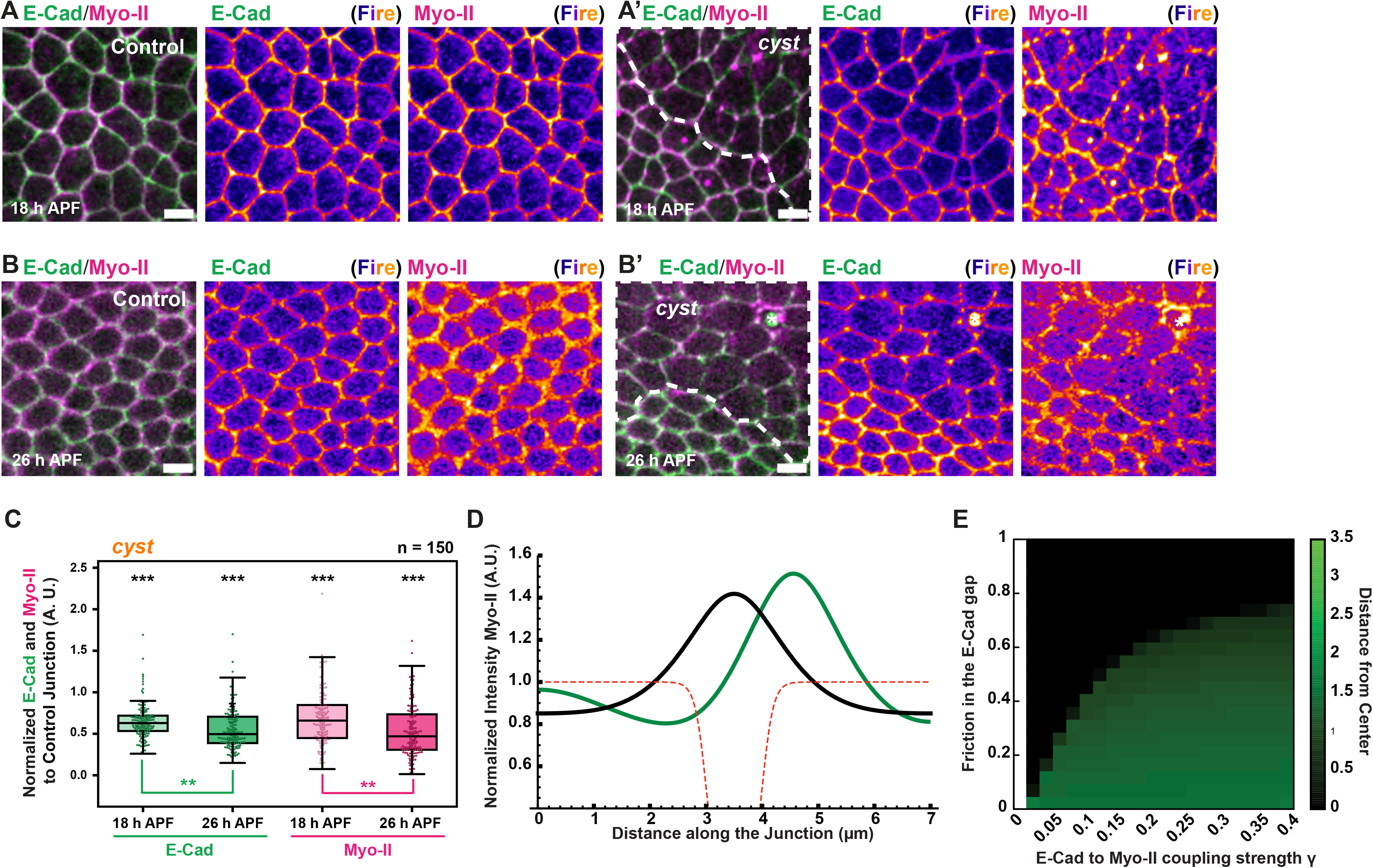
Epithelial mechano-response due to endogenous tissue anisotropic stress. **(A-B’)** E-Cad:GFP and Myo-II:mKate2x3 in control (A,B) and *cyst* mutant (A’,B’) cells at 18 (A,A’) and 26 h APF (B,B’). Dotted lines in (A’) and (B’) mark the positions of cyst mutant cells, identified by the lack of nls:GFP expression (not shown). E-Cad:GFP and Myo-II:mKate2x3 are also displayed with Fire LUT. Asterisk indicates a microchaete. **(C)** Quantification of E-Cad:GFP and Myo-II:mKate2x3 levels at 18 and 26 h APF at junctions of cyst mutant cells, normalized to neighboring control junctions. Box plots show the median, interquartile range, and whiskers at 1.5 × IQR; individual junctions are overlaid as dots. Statistical significance was assessed using Welch’s t-test. Minimum number of animals analyzed for this quantification was 3. **(D-E)** Simulated profiles and phase diagram of Myo-II localization in a spatial model of junctional mechanics, in the presence of an E-Cad gap (red dashed line). As discussed in SI Theory, we consider both the role of E-Cad in dictating friction on the actomyosin cortex, as well as in down-regulating local Myo-II concentration with coupling strength *γ*. For low coupling strength *γ* compared to the effect of friction, Myo-II tends to localize at the boundary of the E-Cad gap, via actomyosin flows from the low friction to the high friction region as observed during cytokinesis (shown as green curve in D, and as green region in the phase diagram of E, which quantifies the distance of the Myo-II peak from the center of the E-Cad gap). For strong coupling strength *γ*, Myo-II tends to accumulate in the center of the E-Cad gap, as observed for E-Cad gaps during interphase (shown as black curve in D and as a black region in the phase diagram of E). Scale bars: 5 µm (A-B’). Unless otherwise indicated, n and N indicate the number of junctions and animals, respectively. Asterisks indicate statistical significance: *p < 0.05, **p < 0.01, ***p < 0.001; ns, not significant.

## Supplementary Video Legends

**Supplementary Video 1. E-Cad and Myo-II dynamics in control tissue.** Time-lapse imaging of E-Cad:GFP (green) and Myo-II:mKate2x3 (magenta) in control tissue of the *Drosophila* dorsal thorax at 18 h APF. Images were acquired every 20 s for 40 min and are shown as a 2-µm-thick apical projection centered on the E-Cad:GFP signal. Yellow asterisks mark microchaetae. Time is shown in seconds (s). Scale bar: 10 µm.

**Supplementary Video 2. E-Cad dynamics upon optogenetic clustering using LARIAT.** Time-lapse imaging of E-Cad:GFP (gray) in the *Drosophila* dorsal thorax at 18 h APF following optogenetic clustering of E-Cad:GFP in a homozygous E-Cad:GFP background using the LARIAT system, which was expressed throughout the tissue. LARIAT activation was initiated before movie acquisition and maintained throughout the time lapse. Images were acquired every 20 s for 40 min and are shown as a 2-µm-thick apical projection centered on the dorsal thorax signal. Yellow asterisks mark microchaetae. Time is shown in seconds (s). Scale bar: 10 µm.

**Supplementary Video 3. E-Cad and Myo-II dynamics in *cyst* mutant tissue.** Time-lapse imaging of E-Cad:GFP (green) and Myo-II:mKate2x3 (magenta) in a large *cyst* mutant clone in the *Drosophila* dorsal thorax at 18 h APF. The *cyst* mutant clone was identified by the absence of nls:GFP expression (not shown). Images were acquired every 20 s for 40 min and are shown as a 2-µm-thick apical projection centered on the E-Cad:GFP signal. Yellow asterisks mark microchaetae. Time is shown in seconds (s). Scale bar: 10 µm.

**Supplementary Video 4. E-Cad and Myo-II dynamics in *α-spec^RNAi^* tissue.** Time-lapse imaging of E-Cad:GFP (green) and Myo-II:mKate2x3 (magenta) in a large *α-spec^RNAi^* clone in the *Drosophila* dorsal thorax at 18 h APF. The *α-spec^RNAi^* clone was identified by CAAX:BFP2 expression (not shown). Images were acquired every 20 s for 40 min and are shown as a 2-µm-thick apical projection centered on the E-Cad:GFP signal. Time is shown in seconds (s). Scale bar: 10 µm.

**Supplementary Video 5. E-Cad and Myo-II dynamics in *cyst^RNAi^*, *α*-*spec^RNAi^* tissue.** Time-lapse imaging of E-Cad:GFP (green) and Myo-II:mKate2x3 (magenta) in a large *cyst^RNAi^, α*-*spec^RNAi^*clone in the *Drosophila* dorsal thorax at 18 h APF. The *cyst^RNAi^, α*-*spec^RNAi^*clone was identified by CAAX:BFP2 expression (not shown). Images were acquired every 20 s for 40 min and are shown as a 2-µm-thick apical projection centered on the E-Cad:GFP signal. Time is shown in seconds (s). Scale bar: 10 µm.

