## Supplementary material for "Differential turnover of apicobasal regulators drives emergent mechano-response and shape homeostasis": Theory Note

**Supplementary Theory Note**

In this Supplementary Note, we provide additional details for our modelling results, both at the level of single-junction dynamics as well as full tissue vertex simulations.

### 1. Analytical modelling of single junctional dynamics

We aim to describe the mechano-chemical dynamics of a single epithelial junction [1–4]. Overall, our full model including the RhoGEF Cyst comprises of a set of 5 coupled equations:

$$\frac{dl}{dt} = \frac{-(l-l_0)-\chi\, F(m,G)}{\tau_{el}} + \sigma_l$$

$$\frac{dl_0}{dt} = \frac{-(l_0-l)}{\tau_j}$$

$$\frac{dm}{dt} = -\frac{m}{l}\frac{dl}{dt} - \frac{m-m_0-\alpha(G-G_0)}{\tau_m} + \sigma_m$$

$$\frac{dc_m}{dt} = -\frac{c_m}{l}\frac{dl}{dt} - \frac{c_m - f_c c_0}{\tau_c} + \sigma_c$$

$$\frac{dG}{dt} = -\frac{G}{l}\frac{dl}{dt} - \frac{G-G_0+\beta(c-c_0)}{\tau_G} + \sigma_G$$

The first equation describes the force balance of a junction, balancing dissipation, elastic stresses with rest length $l_0$ and active contractility $\chi$, while the second equation describes the adaptation over long time scales $\tau_j$ of the rest length $l_0$ – effectively setting up a model of short-term elasticity with long-term viscosity. The last three equations are conservation equation for chemical species (Myo-II, E-Cad, and Cyst) undergoing turnover on a junction of fluctuating length $l$. We also include a noise strength $\sigma_i$ on each of the equations, given recent reports of noisy junctional dynamics [5–9]. Note that $c_m$ describes the fraction of mobile E-cad, while the immobile fraction $c_i$ is simply inversely proportional to junctional length $\frac{dc_i}{dt} = -\frac{c_i}{l}\frac{dl}{dt}$, with the total concentration of E-Cad being $\mathrm{c} = \mathrm{c_m} + c_i$. Furthermore, we can non-dimentionalize all concentrations by their average values without loss of generality ($c_0 = G_0 = m_0 = 1$) as well as set the initial junctional length to 1. In the simplified model, we simply neglect Cyst and lump in a single feedback strength $\gamma$ directly from E-Cad to Myo-II:

$$\frac{dl}{dt} = \frac{-(l-l_0)-\chi\, F(m)}{\tau_{el}} + \sigma_l$$

$$\frac{dl_0}{dt} = \frac{-(l_0-l)}{\tau_j}$$

$$\frac{dm}{dt} = -\frac{m}{l}\frac{dl}{dt} - \frac{m-m_0+\gamma(c-c_0)}{\tau_m} + \sigma_m$$

$$\frac{dc_m}{dt} = -\frac{c_m}{l}\frac{dl}{dt} - \frac{c_m - f_c c_0}{\tau_c} + \sigma_c$$

Note that in both the full and the simplified models, we explicitly neglect molecular mechano-sensing, which would manifest in terms modulating the steady state concentration (or turnover time-scales) of E-Cad [10–13] and/or Myo-II ([14–19], for instance as a function of strain or strain rate. This is because, as we discuss in the main text, we want to use the most parsimonious model, to explore how differential dilution of components with intrinsically different dynamics can give rise, by itself, to emergent mechano-responses.

**Estimation of model parameters**

Firstly, we aim to constrain the timescale of each equation in the problem. For the chemical species, FRAP experiments can give direct access to $\tau_m$, $\tau_c$ and $\tau_G$ as well as validate our assumption of first-order kinetics for turnover. Indeed, the recovery of Myo-II and the RhoGEF Cyst was well-fitted by a single-exponential with little immobile fraction, allowing us to estimate $\tau_m = 87 \pm 3\ s$ and $\tau_G = 43 \pm 4\ s$. Furthermore, the E-Cad recovery was well-fitted by an exponential with immobile fraction, allowing us to jointly estimate $\tau_c = 81 \pm 5\ s$ as well as the mobile fraction $f_c = 0.55 \pm 0.02$. Furthermore, the elastic timescale for junctional response $\tau_{el}$ can be extracted by laser ablation and tracking of junctional recoil, which was again well-described by an exponential, leading to $\tau_{el} = 5s \pm 2\ s$. The timescale of junctional remodelling $\tau_j$ has been previously measured in *Drosophila* embryo as $\tau_j = 60s$ [20,21], which we took in our system, although we have checked by sensitivity analysis that the predicted gap recovery dynamics are qualitatively robust to changes in this parameter (Extended Data Fig. 2E). Furthermore, given that we use throughout normalized cross-correlation functions, the exact values of noise levels $\sigma_l$, $\sigma_m$ and $\sigma_c$ does not have a key effect on the cross-correlation functions, although they rescale the magnitude of the junction length MSD. For simplicity, we set $\sigma_m = \sigma_c = 0.05$, consistent with the short-term variability we see in the Myo-II and E-Cad signal. We then fit $\sigma_l = 0.01$ to get the correct scale for the MSD curves, although we stress again that the trends of noise buffering, and scaling exponents of the MSD curves are not strongly affected by this choice.

Next, we wished to experimentally assess the coupling strengths in the system: the contractility $\chi$ which couples Myo-II to junctional length, the negative feedback $\beta$ from E-Cad to Cyst and the positive feedback $\alpha$ from Cyst to Myo-II. Following a number of previous works [10,22–26], we take active junctional tension to be proportional to contractility $\chi$ as well as to active Myo-II level up to a saturation. We found that this saturation does not qualitatively change the nature of the predicted dynamics and cross-correlation, but stabilizes the junction against extreme divergences, consistent with previous works [22,27,28], and we therefore follow them by taking $F(m) = 2\frac{m-1}{1+m}$ for the simplified model. Importantly, because the timescale of junctional elasticity $\tau_{el}$ is much shorter than all other timescales in the system, we can assume that Myo-II contractility nearly balances junctional elasticity, i.e. $(l - l_0) \approx -\chi F(m)$. This can be tested by plotting junctional length (subtracting its temporal average as a proxy for $l_0$ as the latter evolves much slower) as a function of Myo-II, revealing a noisy, yet consistent negative relationship between the two (Extended Data Fig. 2I). From this, we could extract $\chi = 0.3 \pm 0.05$ in control junctions, and $\chi = 0.25$ in *cyst* mutant junctions. This decrease of contractility in *cyst* mutant tissue is consistent with its role in Myo-II recruitment at the junctions (Extended Data Fig. 3L). Note that in the full model, we have taken the same functional form, but also considering Cyst: $F(m, G) = 2\frac{Gm-1}{1+Gm}$. This is motivated by the fact that contractility requires active Myo-II, which is to first order proportional both to the total amount of Myo-II on the junction and of the concentration of Cyst to activate it.

We also find that although individual values of $\alpha$ and $\beta$ are important to determine the relative magnitude of Cyst accumulation in E-Cad gaps, the overall response is mainly sensitive to their product $\alpha\beta$. This can be made exact if taking the limit of very fast Cyst turnover, in which the product $\alpha\beta$ maps exactly onto the coupling strength $\gamma$ from the simplified model considering only Myo-II and E-Cad. As discussed in the main text, we fit $\alpha$ and $\beta$ (or $\gamma$ in the simplified model) by fitting the dynamics of junctional recovery after a gap, via a simple least-square minimization, running 100 simulations for each parameter regime. This leads to a best-fit of $\gamma = 1.5$ in the simplified E-Cad/Myo-II model, and $\alpha = \beta = 1$ in the full model with Cyst. Interestingly, these values, which are inferred from the control cell dynamics, can be compared to results from perturbation experiments. As discussed in the main text, E-Cad downregulation via RNAi (by a magnitude of $\Delta c = 73\%$, see Extended Data Fig. 1E) leads increased Myo-II at the junction (Extended Data Fig. 1D), as well as increased Cyst (Extended Data Fig. 3H-H''). From this, we can estimate the predicted increase of Myo-II as $\alpha\beta\Delta c$ (full model) or $\gamma\Delta c$ (simplified model). Experimental measurements lead to $\alpha\beta = \gamma = 0.5$. Although these values are lower than the parameter estimation from the best-fit, it is important to note that in perturbation experiments, the decrease in E-Cad is much larger than the physiological decrease seen in gaps, which given the fact that we have a purely linear model for the sake of parsimony, would tend to under-estimate these coefficients as observed here.

Finally, we also tested in mutant conditions how some of the key parameters change compared to control cells. In particular, laser ablations in both *α-Spec$^{RNAi}$* and *cyst* mutant cells revealed a very similar $\tau_{el} = 5s$, arguing this parameter is not changed compared to control cells (Extended Data Fig. 6C,C'). The absolute amount of recoil was also similar in control vs *α-Spec$^{RNAi}$* cells, although it slightly decreased in *cyst* mutant cells (Extended Data Fig. 3R,R'), consistent with the role of Cyst in promoting Myo-II contractility in the model. Furthermore, we also performed FRAP experiments and analysis for E-Cad and Myo-II in *cyst* mutant and *α-Spec*$^{RNAi}$ cells, which revealed that the mobile fraction of E-Cad was largely unchanged in *α-Spec*$^{RNAi}$ cells ($f_c = 0.5$, Extended Data Fig. 6B), but smaller in *cyst* mutant cells ($f_c = 0.4$, Extended Data Fig. 3P). Contrary to other conditions, *cyst* mutant cells displayed a non-negligible immobile fraction in Myo-II upon FRAP experiments, and we therefore added an immobile fraction of Myo-II (20%, Extended Data Fig. 3Q) in simulations.

As a summary, when performing simulations for different conditions, all parameters were kept constant, except the ones described here:

- for *cyst* mutant cells, we input the experimentally observed increase in immobile fraction of E-Cad and Myo-II. Furthermore, the effect of Cyst on Myo-II contractility is encapsulated by the parameter $\alpha$, so we reduce it in simulations. We found that a 33% reduction yielded good prediction for the hole recovery dynamics (Fig. 3G,H), as well as cross-correlation functions (Extended Data Fig. 5K-P).
- For the *Rok*$^{RNAi}$ condition (Extended Data Fig. 5Q-V), we simply decreased contractility $\chi$ by 50%, keeping all other parameters constant.
- for *α-Spec*$^{RNAi}$ cells, we kept all parameters exactly the same as in control condition, as we did not find significant differences in FRAP recovery rate nor recoil upon junction ablations (Extended Data Fig. 6B-F). α-Spec has been shown to have a role on junctional mechanics, providing structural support and elasticity to junctions [29]. Given that we observe increased short-term noise in the MSD of *α-Spec*$^{RNAi}$ cells, even at very short-time scales (i.e. 20s, faster than any of the turnover times of species considered), we therefore doubled the amount of mechanical noise $\sigma_l$ in the *α-Spec*$^{RNAi}$ simulations, leaving all other parameters constant for all the simulations shown in the main figures. An alternative which we also explored is that *α-Spec*$^{RNAi}$ cells could be described by decreasing the parameter $\tau$, corresponding to junctional remodeling. Interestingly, decreasing $\tau$ in the model is sufficient to transition to oscillatory or overshooting modes of junctional dynamics (Extended Data Fig. 2E), something we can observe small signatures for in the data. For instance, junctional recovery after an E-Cad gap is slightly faster than control, and followed by small overshooting phases, as seen for instance by E-Cad levels increasing above 1 around 300 s after initial gap formation (Extended Data Fig. 6G). However, we did not include it for the sake of parsimony in our model, given that increased short-term noise $\sigma_l$ could already recapitulate the core experimental observations (cross-correlations and MSD, see Fig. 4B,C and Extended Data Fig. 7A-I).
- For the *cyst* mutant, *α-Spec*$^{RNAi}$ conditions, we simply combine the previous scenarios, all parameters are the ones from *cyst* mutant cells, except the short-term mechanical noise $\sigma_l$, which is tripled compared to controls (Fig. 4B,C).

- For LARIAT condition, we kept all parameter constants, simply changing the E-Cad mobile fraction to near zero ($f_c = 0.05$, Extended Data Fig. 2L). This is already sufficient, by itself, to lead to an increase in short term noise (Fig. 2G), which is buffered at long-time scales, as evidenced from the fact that the LARIAT MSD, when normalized by control MSD, is larger than 1 for time-scales shorter than 400 s, but smaller than 1 afterwards. We note that this slightly under-estimates the increase in short-term noise, so we also explored the possibility of slightly larger $\sigma_l$ in LARIAT (for instance due to the fact that E-Cad is strongly spatially clustered in this condition). Importantly, this does not change the core results, as even with increased $\sigma_l$, we still find strong noise buffering and long-term MSD in LARIAT significantly smaller to control (Fig. 2I-I'').

**Test of the model**

As discussed in the main text, we have computed a number of independent quantities to quantitatively compare model and experiments.

Firstly, we computed auto- and cross-correlation functions. The cross-correlation function of two signal $f(t)$ and $g(t)$ is computed as $C_{f,g}(\Delta t) =< (f(t)-< f >_t)(g(t+\Delta t)-< g >_t) >_{t,N}$, where $T$ and $<>_{t,N}$ refers to averaging over all time points t and samples $N$. Autocorrelation functions are the same expression, simply correlation the signal with itself. In all cases, we compute the time derivatives (calculated after smoothing over the nearest time-steps) of the quantities of interest (junctional length, local Myo-II Cyst or E-Cad levels), and normalize auto-correlations by their values at $\Delta t = 0$, and all cross-correlations $C_{f,g}(\Delta t)$ by $\sqrt{C_{f,f}(\Delta t = 0)C_{g,g}(\Delta t = 0)}$ (i.e. geometric means of respective auto-correlations functions at $\Delta t = 0$, ensuring that all correlation functions are bounded between -1 and 1). These are for instance shown in Fig. 1B, C, Extended Data Fig. 3E,F,U, Extended Data Fig. 4, Extended Data Fig. 6, Extended Data Fig. 8I,J and Extended Data Fig. 9F,G. This normalization for cross-correlation is particularly suited to compare the amount of cross-correlation between different markers within a single condition, for instance showing that E-Cad and Cyst have a much stronger anti-correlation than E-Cad and Crb in control cells (Extended Data Fig. 8I). However, when comparing across different conditions, this can give misleading information: for instance, in *α-Spec$^{RNAi}$* cells, all correlations between markers are stronger, as they are more junctional fluctuations, but this does not reflect any difference between intrinsic biochemical interactions. To normalize out the effect of the fluctuations, we thus also normalize the cross-correlation function by their maximal values, so that the maximum is always 1 or -1. This is indicated as "Max normalized", and shown in Fig. 2E, Fig. 3C,D, I, J, Fig. 4D, Extended Data Fig. 2J,K, Extended Data Fig. 5 and Extended Data Fig. 7A-I. With this max normalization, the effect of increased junction length fluctuation is removed, so that the anti-correlation peak between E-Cad and Myo-II is the same between control and *α-Spec*$^{RNAi}$ in simulations, mirroring the data of Fig. 4D.

Secondly, we calculated the MSD of the junction length, which quantifies the fluctuations in junctional length at different time scales: $MSD(\Delta t) =< (l(t) - l(t+\Delta t))^2 >_{t,N}$. For a purely diffusive process, i.e. no long-term regulation of junctional length/homeostasis, we expect the MSD to increase linearly as a function of time, while a long-term plateau or sublinear scaling of the MSD are evidence of regulated behavior. Interestingly, we can also use the short-term behavior of the MSD (i.e. sub-minute time-scale) as a measure for the local stochasticity/fluctuating strength experienced in the system, as these are time-scales faster than E-Cad or Myo-II dynamics, and thus at which regulation cannot occur. Both correlation functions and MSD are calculated exactly in the same manner in both data and model. In the model, we run simulations in the presence of stochastic noise $\sigma_i$ for 2h (the timescale of our experimental movies). The exact values of $\sigma_i$ have very little effect on the shape of the cross-correlation, and only set the magnitude of the short-term MSD evolution. We also include some small amount of measurement noise on the intensities $m, c$ which do not impact the MSD, nor the functional shapes of the cross-correlation functions – only the maximal values of cross-correlation – as for instance the Cyst:GFP signal is noisier/with more background noise than for Myo-II and E-Cad fluorescence signals.

In the previous modelling set-up, we test how the junction evolves with respect to stochastic noise over long-time scales. We also complemented this with short term noise-free simulations, where we explicitly introduce a junctional dilation at a specific time point, to better match the quantifications where we focus exclusively on gaps (Fig. 2D, 3E and Extended Data Fig. 6H). Briefly, we run exactly the same set of equation as above, but simply apply in the first equation a transient force of duration $T = 60\ s$ driving transient dilation in the junction $f_0 \propto e^{-\frac{(t-100)^2}{2T^2}}$. We then compute how the junctional length, but also E-Cad, Myo-II and Cyst levels, respond over time to this perturbation.

**Extensions and simplifications of the model**

To get additional insights into the mechanisms at play for junctional homeostasis, we can simplify the model further, in particular by assuming that Cyst dynamics is fast, so that we can write directly the inhibition of E-Cad on Myo-II: $\frac{dm}{dt} = -\frac{m}{l}\frac{dl}{dt} - \frac{m - m_0 + \alpha\beta(c - c_0)}{\tau_m}$.

We first consider the effect of only junctional elasticity, but without feedback, to see whether purely mechanical effects can stabilize junctions. Equations for this read:

$$\frac{dl}{dt} = \frac{l_0 - l - \chi\, F(m)}{\tau_{el}}$$

$$\frac{dm}{dt} = -\frac{m}{l}\frac{dl}{dt} - \frac{m - m_0}{\tau_m}$$

$$\frac{dc_m}{dt} = -\frac{c_m}{l}\frac{dl}{dt} - \frac{c_m - f_c c_0}{\tau_c}$$

Performing a linear stability analysis around the steady state $c_0 = m_0 = l_0 = 1$, we find that the Jacobian matrix reads:

$$J = \begin{matrix} -\frac{1}{\tau_{el}} & -\frac{\chi}{\tau_{el}} & 0 \\ \frac{1}{\tau_{el}} & \frac{\chi}{\tau_{el}} - \frac{1}{\tau_m} & 0 \\ \frac{1}{\tau_{el}} & \frac{\chi}{\tau_{el}} & -\frac{1}{\tau_c} \end{matrix}$$

For simplicity in the analytical expressions, we will non-dimensionalize all timescales by $\tau_{el} = 1$ here. In this case of absent feedback, E-cad plays no role in the dynamics, and the relevant eigenvalues associated with length and Myo-II dynamics read $\lambda_\pm = \frac{1}{2}\left(-\frac{1}{\tau_m} - 1 + \chi \pm \sqrt{\left(\frac{1}{\tau_m} + 1 - \chi\right)^2 - 4/\tau_m}\right)$, which has an oscillatory instability upon a critical contractility $\chi = 1 + 1/\tau_m$, with critical frequency $\lambda_i = \frac{1}{\sqrt{\tau_m}}$, as described in [16,26,30,31]. This shows that even purely elastic junctions, something which is not realistic, as previous measurements have shown significant viscoelasticity [20], can undergo spontaneous instabilities linked to the fact that shrinking a junction leads to concentration of Myo-II, which tends to constrict junctions even further.

It is also theoretically instructive to neglect junctional elasticity, so that no other sources of feedback apart from differential dilution can stabilize junctions. This can arise for fluid tissues, where there are no energy barriers to T1 transitions [7,32,33].

$$\frac{dl}{dt} = \frac{-\chi\, F(m)}{\tau_{el}}$$

$$\frac{dm}{dt} = -\frac{m}{l}\frac{dl}{dt} - \frac{m - m_0 + \alpha\beta(c - c_0)}{\tau_m}$$

$$\frac{dc_m}{dt} = -\frac{c_m}{l}\frac{dl}{dt} - \frac{c_m - f c_0}{\tau_c}$$

We first investigate the scenario of fully mobile E-Cad ($f = 1$), where differential turnover must play a key role. From this model, we can get analytical insights into the stability of junctions with and without feedbacks. Performing a linear stability analysis around the steady state $c_0 = m_0 = l_0 = 1$, we find that the Jacobian matrix reads

$$J = \begin{pmatrix} 0 & -\frac{\chi}{\tau_{el}} & 0 \\ 0 & \frac{\chi}{\tau_{el}} - \frac{1}{\tau_m} & -\frac{\alpha\beta}{\tau_m} \\ 0 & \frac{\chi}{\tau_{el}} & -\frac{1}{\tau_c} \end{pmatrix}$$

Again, for simplicity in subsequent analytical expressions, we will non-dimensionalize all timescales by $\tau_{el} = 1$. In the absence of feedback $\alpha\beta = 0$, plotting the eigenvalues of this matrix reveals an instability (finite real part of eigenvalues without imaginary parts) above a critical value of the contractility $\chi > 1/\tau_m$. Indeed, only one eigenvalue can have a positive value and reads $\lambda = -\frac{1}{\tau_m} + \chi$. Note that in the absence of feedback, parameters linked to E-Cad do not enter in the expression, as the rest of the system of equation is decoupled from it. In the presence of feedback $\alpha\beta > 0$, the threshold for junctional instability is increased, consistent with a stabilizing role for feedback, but eigenvalues can also have imaginary parts, meaning that the another, oscillatory, instability can be triggered, as seen in numerical simulations. At the trace-zero onset of instability, the frequency of these oscillations can be derived analytically as $\omega^2 = \frac{\alpha\beta}{\tau_m^2} + \frac{\alpha\beta}{\tau_m \tau_c} - \frac{1}{\tau_c^2}$. Further instabilities still occur above a critical value of contractility, which is a complex expression of other parameters. Slower Myo-II turnover tend to stabilize junctions, while E-Cad has a non-monotonous effect as seen in the full numerical simulations: starting from very fast E-Cad turnover time $\tau_c$, increasing it tends to stabilize junctions, but above a certain value, it starts to destabilize them by creating a secondary oscillatory instability. This threshold of E-Cad turnover occurs at $\frac{\tau_c}{\tau_m} = \frac{1}{2}(\sqrt{\frac{4}{\alpha\beta} + 1} - 1)$, from which a few interesting points can be noted: i) this threshold is always strictly positive, and depends on the ratio of E-Cad to Myo-II turnover, showing as discussed in the main text that this depends on the differential turnover of junctional components, ii) in the absence of feedback $\alpha\beta = 0$, this threshold can never be reached, and oscillatory instabilities are not observed.

We can also compute in this simplified model how differential dilution leads to junction length compensation, by testing the response to an instantaneous stretch of the junction by a factor $S$ – so that we run the system of equation above with initial condition $l(0) = S, c_m(0) = \frac{f_c}{S}, c_i(0) = \frac{1-f_c}{S}, m(0) = \frac{1}{S}$. Solving for steady state solution $l_\infty$, we find $l_\infty^2 - 2\chi\tau_m \left(1 - \frac{\alpha\beta\tau_c}{\tau_m}\right)(l_\infty - 1) - S^2 = 0$. This reveals that in the absence of feedback ($\alpha\beta = 0$), the long-term junctional length will always be larger than the original stretch ($l_\infty > S$) as seen in the numerical simulations of Extended Data Fig. 2A,B, due to Myo-II being temporally diluted by the stretch. This also reveals that even with feedback, for this model in the absence of any immobile fraction, a slow E-Cad turnover is necessary for compensation after junctional stretching, but that this compensation is always imperfect ($l_\infty > 1$), as shown in Fig. 2B-B''.

Next, we investigated the effect of E-Cad immobile fraction ($f_c < 1$). We found that this critically changed the dynamics, because this provides memory to the system at infinite times – under the current assumption that mobile and immobile pools of E-Cad are fully independent. Indeed, if the junction didn't fully recover yet ($l > 1$), the total E-Cad pool always remains under its homeostatic value due to immobile pool dilution ($c < 1$), leading to a small yet persistent activation of Myo-II that slowly closes the junction to $l = 1$. Quantitatively, this means that immobile E-Cad pool function as an effective elasticity in the system, with timescale of homeostatic return to original values of junctional length with a timing that scales as $\frac{\tau_c}{1-f_c}$ – meaning that it takes an amount of time that diverges with lower and lower immobile fraction $1 - f_c$, irrespective of the amount of stretch $S$. Given that $c_i = \frac{1-f_c}{l}$, the Jacobian of the system with immobile fraction (a 4-by-4 matrix to take into account $c_i$ as a fourth species) reads:

$$J = \begin{matrix} 0 & -\frac{\chi}{\tau_{el}} & 0 & 0 \\ 0 & \frac{\chi}{\tau_{el}} - \frac{1}{\tau_m} & -\frac{\alpha\beta}{\tau_m} & -\frac{\alpha\beta}{\tau_m} \\ 0 & f_c \frac{\chi}{\tau_{el}} & -\frac{1}{\tau_c} & 0 \\ 0 & (1-f_c)\frac{\chi}{\tau_{el}} & 0 & 0 \end{matrix}$$

Finally, for theoretical completeness, we can also consider an alternative full model where both a negative and positive regulator directly regulate Myo-II (e.g. E-Cad directly downregulating Myo-II without intermediary and an activator $a$ with timescale $\tau_a$ upregulating Myo-II):

$$\frac{dl}{dt} = \frac{-\chi\, F(m)}{\tau_{el}} \quad \frac{dl_0}{dt} = \frac{-(l_0 - l)}{\tau_j}$$

$$\frac{dm}{dt} = -\frac{m}{l}\frac{dl}{dt} - \frac{m - m_0 + \alpha\beta(c - c_0) - \gamma(a - a_0)}{\tau_m}$$

$$\frac{dc}{dt} = -\frac{c}{l}\frac{dl}{dt} - \frac{c - c_0}{\tau_c} \quad \frac{da}{dt} = -\frac{a}{l}\frac{dl}{dt} - \frac{a - a_0}{\tau_a}$$

The point of this model is to explore a fully symmetrical situation, where the inhibitors and activators are subject to no regulation other than turnover and dilution, and regulate directly Myo-II. We also for the sake of simplicity omit immobile fractions. Testing again for the response to an instantaneous stretch of the junction by a factor $S$, we find a steady state junctional length of $l_\infty^2 - 2\chi\tau_m\left(1 - \frac{\alpha\beta\tau_c}{\tau_m} + \frac{\gamma\tau_a}{\tau_m}\right)(l_\infty - 1) - S^2 = 0$. This demonstrates, as stated in the main text, that the system needs faster activator than inhibitors ($\tau_c \gg \tau_a$) for junctional length to partially return to its pre-stretch values (compensation), whereas the converse leads to aggravation of stretch, due to activators being more diluted upon stretch.

**Spatial extension of the model**

Although we have concentrated here on a spatially averaged model of a junction, a possible future extension would to be to combine our model of differential dilution with previous works on the role of Myo-II and E-Cad in generating cytoskeletal flows in space and time. Previously, we had considered the role of E-Cad in generating friction on the AJ[23], where local dilution of E-Cad $c(x,t)$ resulted in spatially modulated friction coefficient $\zeta c(x,t)$, and thus self-generated actomyosin flows in the neighbors of dividing cells. This was demonstrated via an active gel model for actomyosin:

$$\frac{dm}{dt} = -\nabla vm - \frac{m - m_0}{\tau_m} + D\Delta m$$

$$-\zeta c(x,t)v(x,t) + \eta\Delta v = -\chi\nabla m$$

where $\eta$ is the viscosity of the actomyosin gel, $D$ its diffusion coefficient and $v(x,t)$ the velocity field representing actomyosin advection. This type of model can be readily combined with our simplified model, where steady-state Myo-II levels are modulated by E-Cad, which then enters both the force balance equation (via friction) and the conservation equation:

$$\frac{dm}{dt} = -\nabla vm - \frac{m - m_0 + \gamma(c - c_0)}{\tau_m} + D\Delta m$$

$$-\zeta cv(x,t) + \eta\Delta v = -\chi\nabla m$$

For $\gamma = 0$, our previous model[23] had shown that a gap in E-Cad results in low friction, and thus triggers self-generated actomyosin flows from the gap, to the boundary between the E-Cad high and E-Cad low domains (Extended Data Fig. 10D). This results in depletion of Myo-II from the E-Cad gap. However, positive values of $\gamma$ go in the opposite direction, as they favor Myo-II accumulation within the gap. To reconcile these findings, we generated a phase diagram to quantify whether the peak of MyoII tends to be in the center of the gap, or on its side, finding a transition above a critical value of $\gamma$ compared to the amount of friction decrease in the E-Cad gap (Extended Data Fig. 10D-E), which could suggest that junctions during or outside of cytokinesis could be in different regions of the phase diagram. Interestingly, we had also shown before that the size of the E-Cad gap matters, as large gaps allow more easily the formation of actomyosin flows[23]. This could also explain why the large E-Cad gaps generated by the cytokinetic furrow give rise to actomyosin flows away from the gap, while smaller E-Cad gaps studied in this work during interphase are repaired by local Myo-II accumulation.

Having characterized junction dynamics at the level of individual length and tension (contractility) variables, we next turn to tissue-scale mechanics.

### 2. **Active vertex model for epithelial monolayers**

To understand junction-scale mechanochemical dynamics from tissue-level observables, we utilize the active vertex model (AVM), a standard framework for describing the collective mechanics and motility of confluent epithelial monolayers [7,35–37]. In this framework each cell *i* of the tissue is represented as a convex polygon whose vertices are shared with neighboring cells. The mechanical state of the entire tissue is governed by a free energy functional that penalizes deviations in each cell's geometry from a preferred cell area $A_0$ and a preferred cell perimeter $P_0$ (Fig. 4E):

336 $$E = \sum_i K_A(A_i - A_0)^2 + K_P(P_i - P_0)^2$$

The first term penalizes deviations of cell area $A_i$ from a preferred area $A_0$, with modulus $K_A$. This reflects the resistance of epithelial cells to volumetric compression, arising from cytoplasmic incompressibility, basal membrane elasticity, and osmotic pressure. The second term penalizes deviations of the cell perimeter $P_\mathrm{i}$ from a preferred perimeter $P_0$, with modulus $K_P$. This captures the elastic cost of deforming the actomyosin cortical ring, which resists both stretching and compression of cell-cell junctions. Together, the competition between area and perimeter regulation produces a rigidity transition controlled by the dimensionless shape index $q_0 = P_0/\sqrt{A_0}$: below a critical value $p^* \approx 3.81$, the tissue behaves as a solid with finite shear modulus and junctions that resist rearrangement, whereas above $p^*$ the tissue is fluid-like with vanishing shear modulus and cells freely exchange neighbors via T1 transitions [7,35–37]. A central assumption underlying these models is that each cell actively maintains $K_P$, $P_0$, $K_A$, and $A_0$ at precise values, yet the molecular mechanisms by which any of these parameters are established, how they respond to mechanical perturbations, and what determines their homeostatic values in a living tissue remain largely unknown [38–40]. Note that a direct mapping from our junctional model to the vertex model would be to take an energy with homeostatic junction length instead of perimeter, of the type $E = \sum_i K_A(A_i - A_0)^2 + \sum_i \sum_j K_P\left(l_{ij} - l_0\right)^2$, where $l_{ij}$ is the junction *j* of cell *i*, so that each junction has its own preferred length $l_0$ arising from our mechano-chemical mechanism. However, since the perimeter is a sum over its bounding edges, the classical AVM is closely related to a sum of squared edge lengths. Indeed, these two different variants of the AVM have been shown to reproduce the similar rigidity transition[32,34], so we use the more classical version of the AVM to link better to the past literature.

**Equations of motion and cell activity**

The dynamics of each vertex position $r_\alpha$ of a cell *i* (and shared by other cells $\mathcal{N}(\alpha)$) are governed by an overdamped equation of motion [37,41], that balances viscous drag against forces derived from the energy functional and an active noise term $\sigma_0$ (Fig. 4E'):

$$\mu \frac{\mathrm{d}r_\alpha}{\mathrm{d}t} = -\nabla_\alpha E + \frac{1}{3} \sum_{k \in \mathcal{N}(\alpha)} \sigma_0 \widehat{n_i}(t)$$

where $\mu$ is the effective mobility coefficient of the vertex against the substrate and neighboring cells, $\sigma_0$ is the self-propulsion speed of the cell *i* contributing to the vertex noise, and $\widehat{n_i}(t) = (cos\,\theta_i\,, sin\,\theta_i)$ is the instantaneous polarity unit vector of cell *i*. The polarity angle $\theta_i$ of each cell evolves according to an angular diffusion process:

$$\frac{d\theta_i}{dt} = \eta_i(t) \qquad \langle \eta_i(t)\eta_j(t') \rangle = 2D_r \delta_{ij} \delta(t - t')$$

where $\eta_i(t)$ is a white noise term and $D_r$ is the rotational diffusion coefficient of cell polarity[7,37,41]. The ratio $\sigma_0/D_r$ determines the persistence length of cell trajectories and, together with $p_0$, controls whether the tissue is in a solid or fluid state [7,42].

Importantly, in a confluent epithelium each vertex is shared by exactly three cells under normal topology (away from T1 transitions). The net active force at a vertex therefore receives contributions from all three surrounding cells. This distributed activity rule means that: (i) if neighboring cells have aligned polarity, the net active force at the shared vertex is large and coherent, promoting coordinated tissue flow; (ii) if neighboring cells have disordered polarity, contributions cancel and the net force at the vertex is suppressed. The parameter $\sigma_0$ therefore sets the single-cell propulsion (or the junctional vertex noise) magnitude, while the effective tissue-level activity depends on the degree of polarity alignment, which is itself controlled by $D_r$.

In the limit of highly disordered polarity driven by large rotational diffusion ($D_r$), cell orientations randomize rapidly, leaving $\sigma_0$ to act simply as an isotropic active noise amplitude that fluidizes the tissue without imparting a macroscopic migration direction. Conversely, in the low-$D_r$ limit, cell orientations remain highly persistent, fostering strongly correlated velocity fields and directed collective migration.

To allow for macroscopic tissue remodeling, the AVM handles cellular neighbor exchanges through T1 topological transitions. Specifically, if a junction shrinks below a threshold length $\ell^*$, it undergoes an instantaneous flip to form a new, perpendicularly oriented junction. Because these rearrangements dictate tissue flow, the T1 transition rate acts as a critical experimental observable that sharply increases as the monolayer crosses from a solid to a fluid state [7,35–37].

Note that we explicitly neglect here more complex modes of feedbacks, such as junctional or area mechano-sensing, which have been increasingly explored in recent years [6,15,43–45], as we seek to understand the most minimal form of shape homeostasis, following our junctional model.

**Tissue-scale simulations with Vertex Model**

Tissue-scale simulations were performed using the Active Vertex Model (AVM) implemented in the CellGPU framework [41] following the energy functional and equations of motion described above (see Active vertex model for epithelial monolayers). Neighbour exchanges, which are required for tissue fluidity, occur via T1 transitions: when a junction shortens below a threshold of 0.04 (in units of $\sqrt{A_0}$), it is instantaneously flipped to form a new perpendicular contact between the two previously non-adjacent cells.

All simulations used tissues of $N$ = 36 cells within a fully periodic square domain. At this size, a tissue in a disordered hexagonal-like packing contains approximately 108 junctions, providing sufficient junctional statistics for MSD analysis while keeping the computational cost of the 700-point parameter sweep tractable. Throughout, the preferred area $A_0$ = 1, area modulus $K_A$ = 1, vertex mobility $\mu$ = 1, preferred shape index $q_0^* = \frac{P_0}{\sqrt{A_0}} = 3.8$, and rotational diffusion coefficient $D_r$ = 1 were held fixed. Differences in tissue behaviour therefore arise solely from changes in two parameters: the perimeter modulus $K_P$, which sets the energetic cost of deviating from the preferred cell perimeter and reflects the effective junctional stiffness controlled by the E-Cad/Cyst/Myo-II mechanochemical circuit (Fig. 4E, Extended Data Fig. 7K); and the mechanical noise amplitude $\sigma_0$, which corresponds to the self-propulsion speed in the equations of motion above; with $D_r$ = 1 fixed, the polarity randomises on the timescale of a single time unit so that $\sigma_0$ acts as an isotropic active noise amplitude driving short-timescale junction fluctuations, rather than producing directed migration (Fig. 4E', Extended Data Fig. 7L).

Each simulation was initialized from a random Voronoi tessellation of uniformly distributed seed points. To remove artefacts of this initial configuration, the tissue was equilibrated for 20,000 integration steps (100 simulation time units at $\Delta t$ = 0.005) with mechanical noise already active, allowing cell shapes and packing to relax to a disordered steady state consistent with the imposed $K_P$ and $\sigma_0$. Production runs then continued for a further 400,000 steps (2,000 time units), with vertex positions, cell areas, perimeters, and junction connectivity recorded every 10 steps (every 0.05 time units), yielding 40,000 frames per trajectory. This production run length was chosen to obtain robust statistics for junction-length fluctuations and T1 rearrangement rates within each parameter regime, rather than to capture the long-term collective migration patterns typically probed in longer AVM simulations. For each $(K_P, \sigma_0)$ combination, 20 independent realisations were run from different random initial configurations and noise seeds; all reported metrics are ensemble means with uncertainty quantified as the standard error across the 20 realisations.

To systematically explore how tissue mechanics depends on model parameters, we swept $K_P \in [0.1, 2.0]$ in steps of 0.1 and $\sigma_0 \in [0.15, 1.0]$ in steps of 0.025, producing a parameter grid. This grid spans the solid-like regime (high $K_P$, low $\sigma_0$), where junctions resist deformation and T1 transitions are rare, through to the fluid-like regime (low $K_P$, high $\sigma_0$), where cells exchange neighbours frequently and junctional fluctuations are large, covering both sides of the rigidity transition at $p_0^* \approx 3.81$. Simulated observables from this grid were compared directly with the corresponding experimental measurements across genetic conditions, without fitting model parameters to individual conditions (Fig. 4F-K, Extended Data Fig. 7K-O).

**Quantification of AVM Simulation Observables**

The following quantities were computed for each $(K_P, \sigma_0)$ combination from the production trajectories and averaged across the 20 realisations.

*Mean shape index:* The mean shape index $q = \langle \frac{P}{\sqrt{A}} \rangle$ was computed by averaging over all $N$ cells at steady state. This serves as the primary readout of tissue mechanical state, reflecting how elongated cells are relative to an equivalent circle and is directly comparable to the shape indices measured from segmented experimental images (Extended Data Fig. 7O).

*Junction-length MSD:* To characterise the dynamics of individual cell–cell contacts, we computed the mean squared displacement (MSD) of junction lengths as a function of lag time $\tau$:

$$C(\tau) = \langle [\ell(t + \tau) - \ell(t)]^2 \rangle_{t,junctions}$$

where $\ell(t)$ is the length of a given junction at time *t*. For each junction, the evolution of length over time was extracted from the recorded vertex positions and the MSD computed using a fast Fourier transform (FFT)-based autocorrelation algorithm that efficiently handles the full lag-time range. Junctions created or destroyed by T1 transitions were included only for the portions of the trajectory during which they existed. Per-realization MSD curves were averaged over all junctions, and the ensemble mean and standard error were computed across all the realizations (Fig. 4F,H,J). Two summary scalars were extracted: a short-time

MSD (mean over the first 0.2% of the lag range, capturing ballistic-to-diffusive crossover dynamics; Extended Data Fig. 7M) and a long-time MSD (mean over the final 20% of the lag range, capturing the plateau or slow-diffusion regime; Extended Data Fig. 7N), enabling quantitative comparison with the long-time and short-time junction MSD values extracted from experimental kymograph data.

*Cell centroid MSD:* Cell centre-of-mass positions were tracked across frames and the MSD of cell displacements computed using the same FFT-based algorithm. This provides a complementary measure of collective cell motility alongside the junction-length fluctuations, and is shown in Extended Data Fig. 7J alongside the junction MSD.

*T1 transition rate:* Neighbor-exchange events were identified by monitoring changes in cell adjacency between successive saved frames. The T1 rate was defined as the total number of detected events divided by the number of cells and the total production time, giving units of events per cell per simulation time unit. This rate characterizes the timescale of tissue remodeling (Fig. 4J) and its trend is therefore directly comparable to the experimentally measured T1 rates per junction per unit time after appropriate unit conversion (Fig. 4K).
